# Comparative genome and transcriptome analyses revealing interspecies variations in the expression of fungal biosynthetic gene clusters

**DOI:** 10.1101/2020.04.17.047639

**Authors:** Hiroki Takahashi, Maiko Umemura, Masaaki Shimizu, Akihiro Ninomiya, Yoko Kusuya, Syun-ichi Urayama, Akira Watanabe, Katsuhiko Kamei, Takashi Yaguchi, Daisuke Hagiwara

## Abstract

Filamentous fungi produce various bioactive compounds that are biosynthesized by a set of proteins encoded in biosynthetic gene clusters (BGCs). For an unknown reason, large parts of the BGCs are transcriptionally silent under laboratory conditions, which has hampered the discovery of novel fungal compounds. The transcriptional regulation of fungal secondary metabolism is not fully understood from an evolutionary viewpoint. To address this issue, we conducted comparative genomic and transcriptomic analyses using five closely related species of the *Aspergillus* section *Fumigati*: *Aspergillus fumigatus, Aspergillus lentulus, Aspergillus udagawae, Aspergillus pseudoviridinutans*, and *Neosartorya fischeri*. From their genomes, 298 secondary metabolite (SM) core genes were identified, with 27.4% to 41.5% being unique to a species. Compared with the species-specific genes, a set of section-conserved SM core genes was expressed at a higher rate and greater magnitude, suggesting that their expression tendency is correlated with the BGC distribution pattern. However, the section-conserved BGCs showed diverse expression patterns across the *Fumigati* species. Thus, not all common BGCs across species appear to be regulated in an identical manner. A consensus motif was sought in the promoter region of each gene in the 15 section-conserved BGCs among the *Fumigati* species. A conserved motif was detected in only two BGCs including the *gli* cluster. The comparative transcriptomic and *in silico* analyses provided insights into how the fungal SM gene cluster diversified at a transcriptional level, in addition to genomic rearrangements and cluster gains and losses. This information increases our understanding of the evolutionary processes associated with fungal secondary metabolism.

**Author summary:** Filamentous fungi provide a wide variety of bioactive compounds that contribute to public health. The ability of filamentous fungi to produce bioactive compounds has been underestimated, and fungal resources can be developed into new drugs. However, most biosynthetic genes encoding bioactive compounds are not expressed under laboratory conditions, which hampers the use of fungi in drug discovery. The mechanisms underlying silent metabolite production are poorly understood. Here, we attempted to show the diversity in fungal transcriptional regulation from an evolutionary viewpoint. To meet this goal, the secondary metabolisms, at genomic and transcriptomic levels, of the most phylogenetically closely related species in *Aspergillus* section *Fumigati* were compared. The conserved biosynthetic gene clusters across five *Aspergillus* species were identified. The expression levels of the well-conserved gene clusters tended to be more active than the species-specific, which were not well-conserved, gene clusters. Despite highly conserved genetic properties across the species, the expression patterns of the well-conserved gene clusters were diverse. These findings suggest an evolutionary diversification at the transcriptional level, in addition to genomic rearrangements and gains and losses, of the biosynthetic gene clusters. This study provides a foundation for understanding fungal secondary metabolism and the potential to produce diverse fungal-based chemicals.

## Introduction

Filamentous fungi produce various small molecules known as secondary metabolites (SMs, also known as natural products), which are thought to contribute to their survival in environmental niches (1, 2). Fungal SMs are biosynthesized by enzyme sets, which include backbone and tailoring enzymes. The backbone enzymes are represented by non-ribosomal peptide synthetase (NRPS) and polyketide synthase (PKS) (3). The backbone gene and additional genes encoding a tailor enzyme, transcriptional regulator, and efflux pump are often arrayed in a biosynthetic gene cluster (BGC). Fungi, including phytopathogens and human pathogens, possess large numbers of SM gene cluster in their genomes, which indicates a potent ability to produce a myriad of metabolites that could be used to impact humans (4–6).

Comparative genomic studies were conducted and revealed that SM gene clusters are species-specific or narrowly taxonomically distributed within a certain group of species. Lind et al. (7) showed that 91.6%–96.1% of SM gene clusters are species-specific among *Aspergillus niger, Aspergillus oryzae, Aspergillus nidulans*, and *Aspergillus fumigatus*, and that none of the clusters is shared by all four species. This is in sharp contrast to primary metabolic genes, which are 7.5%–15.4% species-specific (7). Comparisons between more closely related species using *A. fumigatus* and *Neosartorya fischeri* or *Aspergillus novofumigatus*, which all belong to *Aspergillus* section *Fumigati* revealed that 30.3% or 70.5% of *A. fumigatus* SM genes were shared, respectively (8,9). Comprehensive genomic studies in *Aspergillus* sections *Nigri* and *Flavi* also showed overlaps of SM genes at specific rates. These reports provide an evolutionary insight into how secondary metabolic pathways evolve and degenerate in filamentous fungi (5,6).

For deeper insights into fungal SM gene variation, intraspecies variations among the BGCs were investigated using the genomes of 66 *A. fumigatus* strains (10). The evolutionary traits of SM gene clusters, such as genetic polymorphisms, genomic rearrangements, gene gains or losses, and horizontal gene transfers (Interspecies diversification), were identified and may affect fungal SM production. Indeed, intraspecies microevolutions in SM gene clusters have been reported in *A. fumigatus* for fumitremorgin, trypacidin, and fumigermin and in *A. flavus* for aflatoxin B1 (11–14). Such interspecies variations may determine the ecological properties of the fungi, because bioactive compounds play protective and weaponized roles in competitive niches (3).

The fungal SM gene clusters, in general, are transcriptionally silent under laboratory condition, which makes it difficult for us to comprehensively explore fungal SMs and to understand ecological role of the SMs (14). For example, genomic study revealed that more than 40 SM backbone genes were found in *Aspergillus fumigatus, Aspergillus niger*, and *Aspergillus oryzae*, 74.2 to 91.4% of which were not expressed or expressed at ultimately low level in any of the cell types, hyphae, resting conidia, or germinating conidia (15). The reason why a large part of SM genes are silent under the laboratory controlled conditions remains to be addressed. One plausive explanation for this is unknown ecological cues that cannot reproduced under laboratory conditions and are involved in triggering silent fungal SM production. This may occur because of the loss of transcriptional ability owing to the diversification of machinery regulating SM gene clusters. Although many researchers have attempted to verify this hypothesis (13,14,16,17), whether the SM gene clusters in their genomes are “alive” from an evolutionary perspective has been poorly studied.

In the present study, we attempted to investigate whether the fungal SM gene clusters that are conserved among different species are transcriptionally regulated in an identical manner. We performed comparative genomics combined with transcriptome analyses focusing on SM genes using five closely related *Aspergillus* section *Fumigati* species. Comparisons of the transcriptomes generated under four different conditions revealed that the section-conserved (SC) SM core genes were transcriptionally more active than the specie-specific SM core genes. Among the species, expression profiles of the SC BGCs were diverse, suggesting that SM had independently evolved at the transcriptional level in the distributed species. These findings increase our understanding of how the transcriptional regulation of fungal SMs has diversified during the course of evolution.

## Results

### Genomic sequences of five different species of *Aspergillus* section *Fumigati*

Genetically closely related fungal species that belong to the same section, *Aspergillus* section *Fumigati*, were used for a comparative genomic study. The genome data of *A. fumigatus* (18), *Aspergillus lentulus* (19), *Aspergillus udagawae* (20), and *Neosartrya fischeri* (*Aspergillus fischeri*) (21) are available at the NCBI and were retrieved for this study (Fig 1A). In addition to the four strains, we sequenced *Aspergillus pseudoviridinutans* IFM 55266 (22), which also belongs to *Aspergillus* section *Fumigati*. The numbers of proteins in the fungi are summarized in Table 1. In total, 6,277 proteins were orthologous in the five *Fumigati* strains (Fumigati-conserved; Fig 1B), and among them, 3,598 proteins were shared by the other 17 available *Aspergillus* genomes (Asp-conserved). Notably, 3,017, 3,840, 4,276, 4,431, and 3,316 proteins were species-specific to *A. fumigatus, A. lentulus, A. udagawae, A. pseudoviridinutans*, and *N. fischeri*, respectively (Fig 1B). A genomic synteny analysis revealed that most of the *A. fumigatus* genomic region is covered by sequences of *A. lentulus, A. udagawae, A. pseudoviridinutans*, and *N. fischeri* (Fig 1C). The numbers of syntenic genes were 8,486 (86.23% of *A. fumigatu*s genes), 8,375 (85.24%), 8,336 (84.84%), and 8,559 (87.11%), respectively. This suggested that *N. fischeri* is the closest relative to *A. fumigatu*s among the *Fumigati* species, which was supported by the phylogenetic tree shown in Fig 1A.

**Table 1.** The numbers of protein-coding and *SM* core genes in the fungal strains used in this study.

| Strains | Genome size [M] | Predicted proteins | NRPS or NRPS-like | PKS or PKS-like | Hybrid | TS | Total |
| --- | --- | --- | --- | --- | --- | --- | --- |
| *A. fumigatus* Af293 | 29.38 | 9,840 | 18 | 15 | 1 | 5 | 39 |
| *A. lentulus* IFM 54703<sup>T</sup> | 30.96 | 11,205 | 24 | 18 | 4 | 5 | 51 |
| *A. udagawae* IFM 46973<sup>T</sup> | 32.19 | 11,792 | 31 | 35 | 3 | 6 | 75 |
| *A. pseudoviridinutans* IFM 55266<sup>T</sup> | 33.20 | 11,927 | 33 | 38 | 5 | 6 | 82 |
| *N. fischeri* NRRL 181<sup>T</sup> | 32.55 | 10,406 | 28 | 17 | 1 | 5 | 51 |

**Fig 1.**
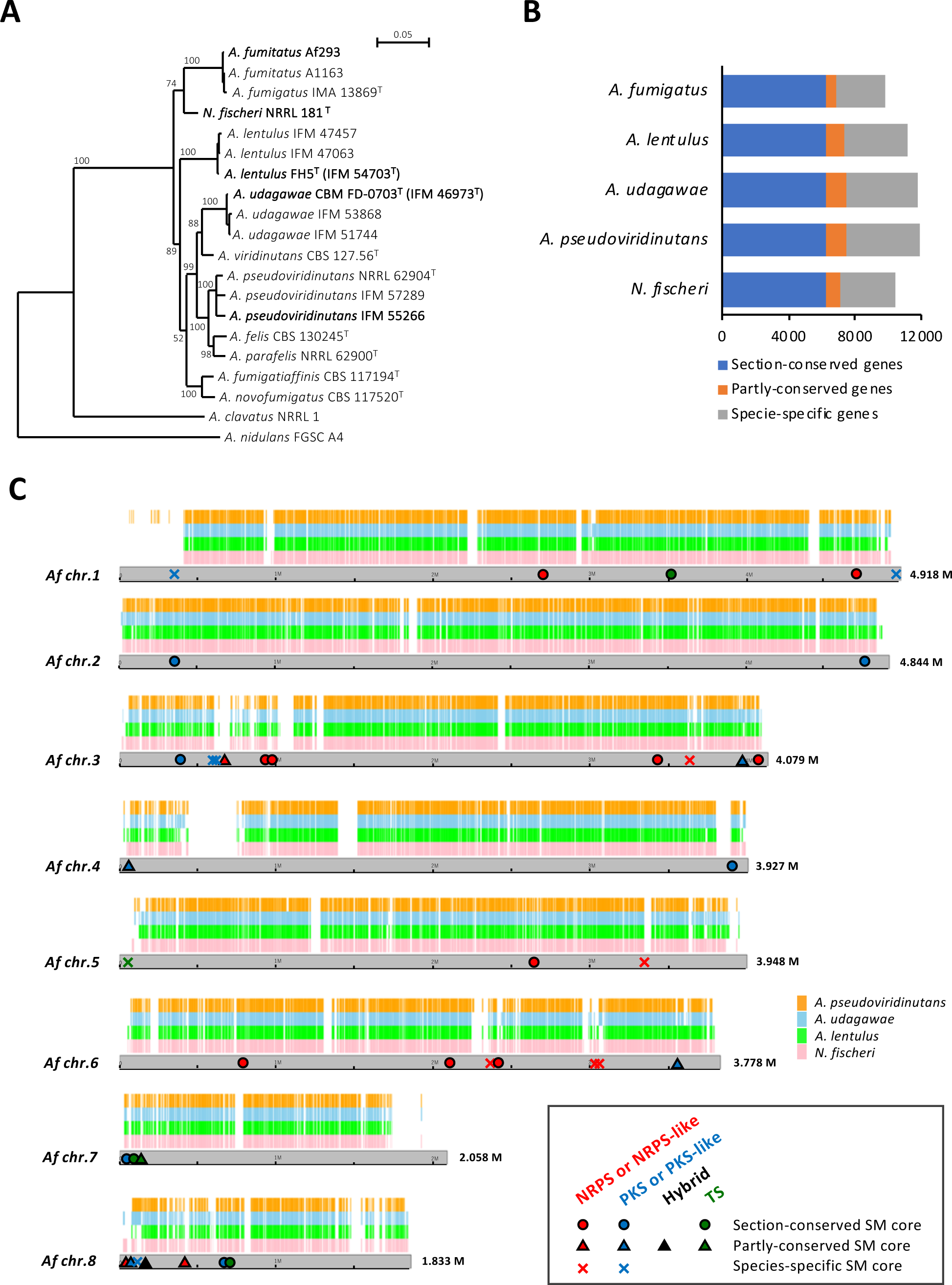
The *Aspergillus* section *Fumigati* strains used for the comparative genomic analysis. (A) Phylogenetic tree of 20 strains from 12 species. The tree was constructed by Clustal X with a neighbor-joining analysis using partial β-tubulin and calmodulin gene sequences. The distances between sequences were calculated using Kimura’s two-parameter model. The bootstrap was conducted with 1,000 replications. The main strains used in the study are indicated in bold. (B) The numbers of genes that are conserved across the section or that are species specific. (C) Whole-genome synteny plot with positions of *A. fumigatus*’ SM core genes. The syntenic genes are mapped to *A. fumigatus*’ chromosomes. The positions of the section-conserved, partly-conserved, and species-specific SM core genes of *A. fumigatus* are indicated with circles, triangles, and crosses, respectively. Genes encoding NRPS or NRPS-like protein are colored in red, PKS or PKS-like proteins are in blue, and TS proteins are in green.

### Comparative genomics regarding SM core genes

The SM core genes encoding PKS, NRPS, a PKS-NRPS hybrid, and terpene synthase (TS) were identified from the genome data using a combination of a BLAST program and the PKS/NRPS Analysis Web-site (http://nrps.igs.umaryland.edu/), as well as manual inspection. In total, 39, 51, 75, 82, and 51 genes were identified in *A. fumigatus, A. lentulus, A. udagawae, A. pseudoviridinutans*, and *N. fischeri*, respectively (Table 1, S1 Table). Compared with *A. fumigatus*, 24, 23, 23, and 27 of the SM proteins were conserved, having identities greater than 80%, in *A. lentulus, A. udagawae, A. pseudoviridinutans*, and *N. fischeri*, respectively, which revealed the high overlapping of SM core genes among the species belonging to the same section (Fig 2A). Notably, 19 genes were shared among all five species, which we termed the SC SM core genes. Meanwhile, there were 11, 13, 30, 34, and 14 species-specific SM core genes in *A. fumigatus, A. lentulus, A. udagawae, A. pseudoviridinutans*, and *N. fischeri*, respectively (Fig 2B). In total, 298 SM core genes were identified and grouped into 160 orthologous types. A cladogram was generated based on a binary matrix (presence/absence of the SM proteins), which revealed that *A. fumigatus* and *N. fischeri* are most closely related on the basis of the SM core protein distribution across species (Fig 2C). This relationship resembled the genetic phylogeny (Fig 1A), which suggests that the diversification of SM core genes occurred along with speciation inside the *Fumigati* section.

**Fig 2.**
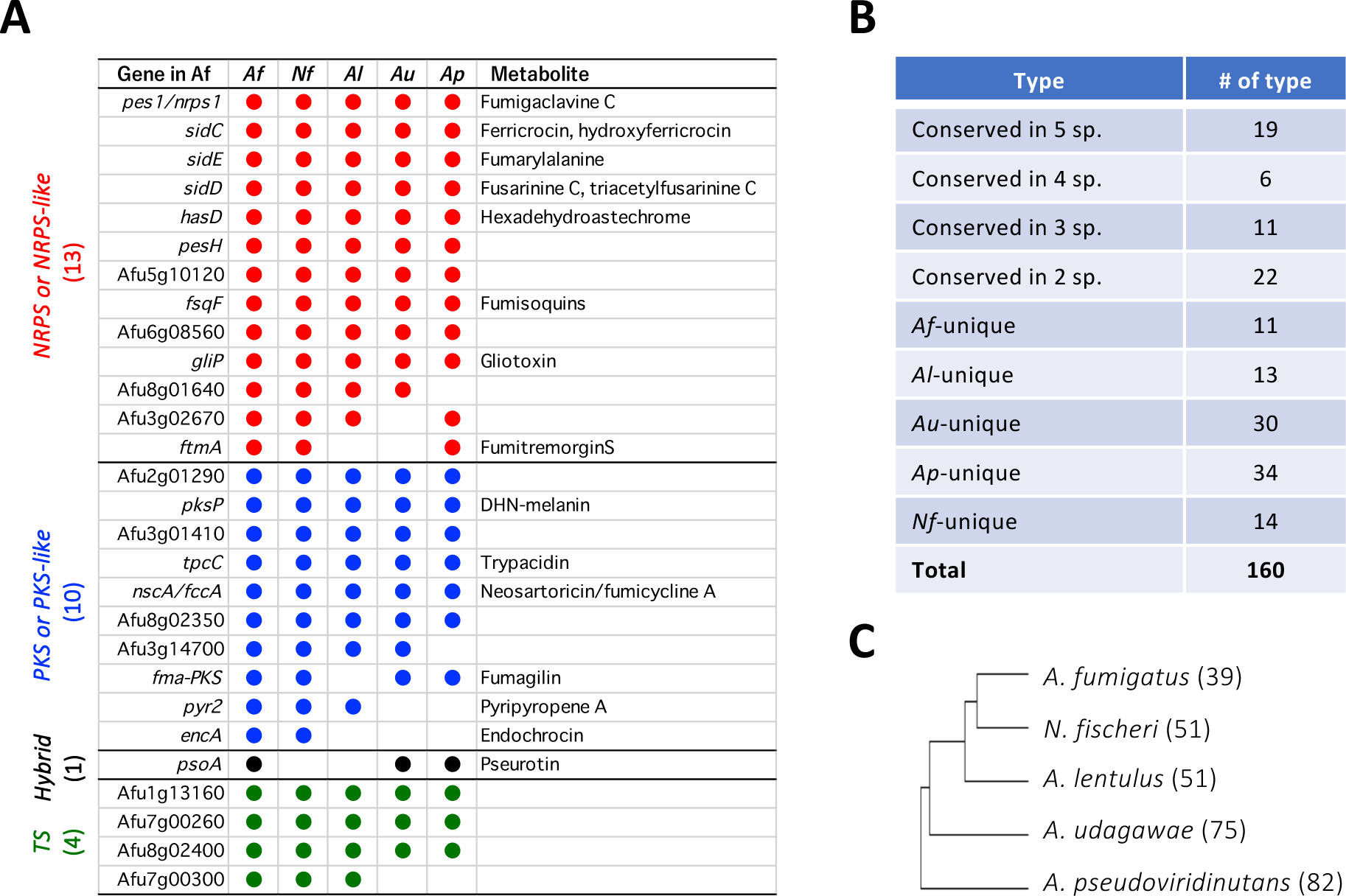
The SM core genes conserved across *Aspergillus* section *Fumigati*. (A) Summary of *A. fumigatus* SM core genes that are conserved in the other *Fumigati* species. *Af*: *A. fumigatus*; *Nf*: *N. fischeri*; *Al*: *A. lentulus*; *Au*: *A. udagawae*; *Ap*: *A. pseudoviridinutans*. (B) Summary of the numbers of SM gene types. (C) A cladogram was constructed using a binary matrix (presence/absence of the PKSs and NRPSs) with Cluster 3.0 (http://bonsai.hgc.jp/~mdehoon/software/cluster/software.htm#ctv). The tree and heat map were constructed using Tree View. The numbers of SM core genes are shown in parentheses behind the species’ names.

### Characterization of BGCs for SC SM genes

The 19 SC SM gene types include 10 NRPSs, 6 PKSs, and 3 TSs, among which 10 genes were previously characterized in *A. fumigatus* as being involved in the biosynthesis of fumigaclavine C (23), ferricrocin (24), fumarylalanine (25), fusarinine C (24), hexadehydroastechrome (26), fumisoquins (27), gliotoxin (28), DHN-melanin (29), trypacidin (12,30), and neosartoricin/fumicyclines (31,32) (Table 2). It is notable that the BGCs for 10 NRPSs and 6 PKSs were well conserved among the species in terms of gene composition, gene order, and encoded protein similarity (S1 Fig), and they are hereafter designated SC-BGC1 to SC-BGC15, with SC-BGC6 including two SC SM core genes.

**Table 2.**
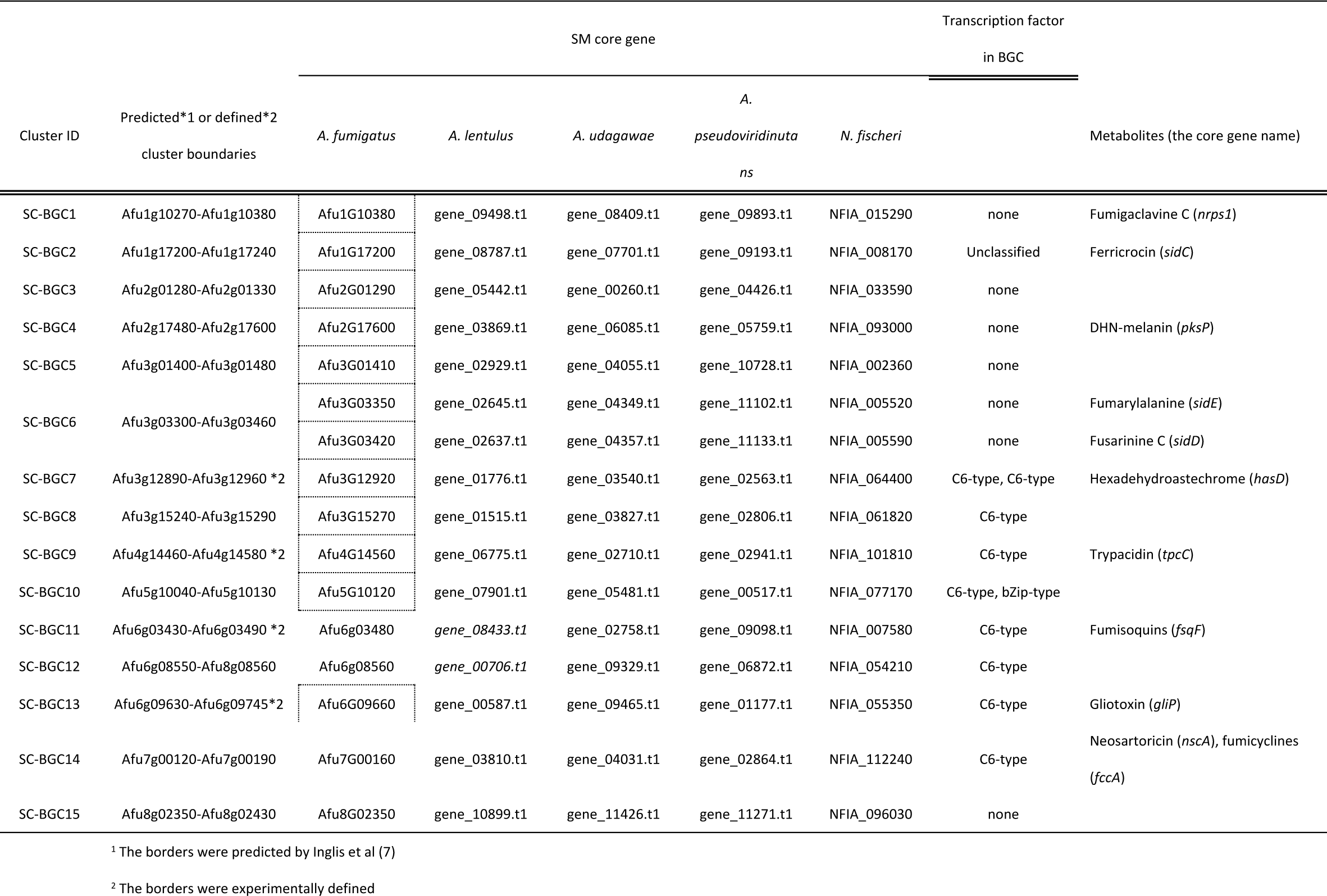
List of section-conserved biosynthetic gene clusters (BGCs)

### Comparative transcriptome analysis of SM core genes

To gain comprehensive insights into SM gene expression, the fungal strains were cultivated under four different medium-based conditions, potato dextrose broth (PDB), Czapek-Dox (CD), Sabouraud broth (SB), and the asexual stage on potato dextrose agar (PDA). In *A. fumigatus*, the median Transcripts Per Kilobase Millions (TPMs) of Asp-conserved genes (n = 3,598) were 51.9, 35.8, 21.1, and 64.9 for PDB, CD, SB, and PDA cultivations, respectively, which were much higher than those of Af_unique genes (n = 3,017) (Fig 3A). This was also the case for the other species, *A. lentulus, A. udagawae, A. psuedoviridinutans, and N. fischeri*. These data showed that a set of genes that are well conserved among *Aspergilli* was transcriptionally more active than the species-specific genes in all the species. Interestingly, *Fumigati*-unique genes (n = 61) that are conserved only among the five *Fumigati* species also showed low expression levels (Fig 3A). Thus, widely conserved genes tended to be expressed with more frequency and at higher intensities than the narrowly distributed genes.

**Fig 3.**
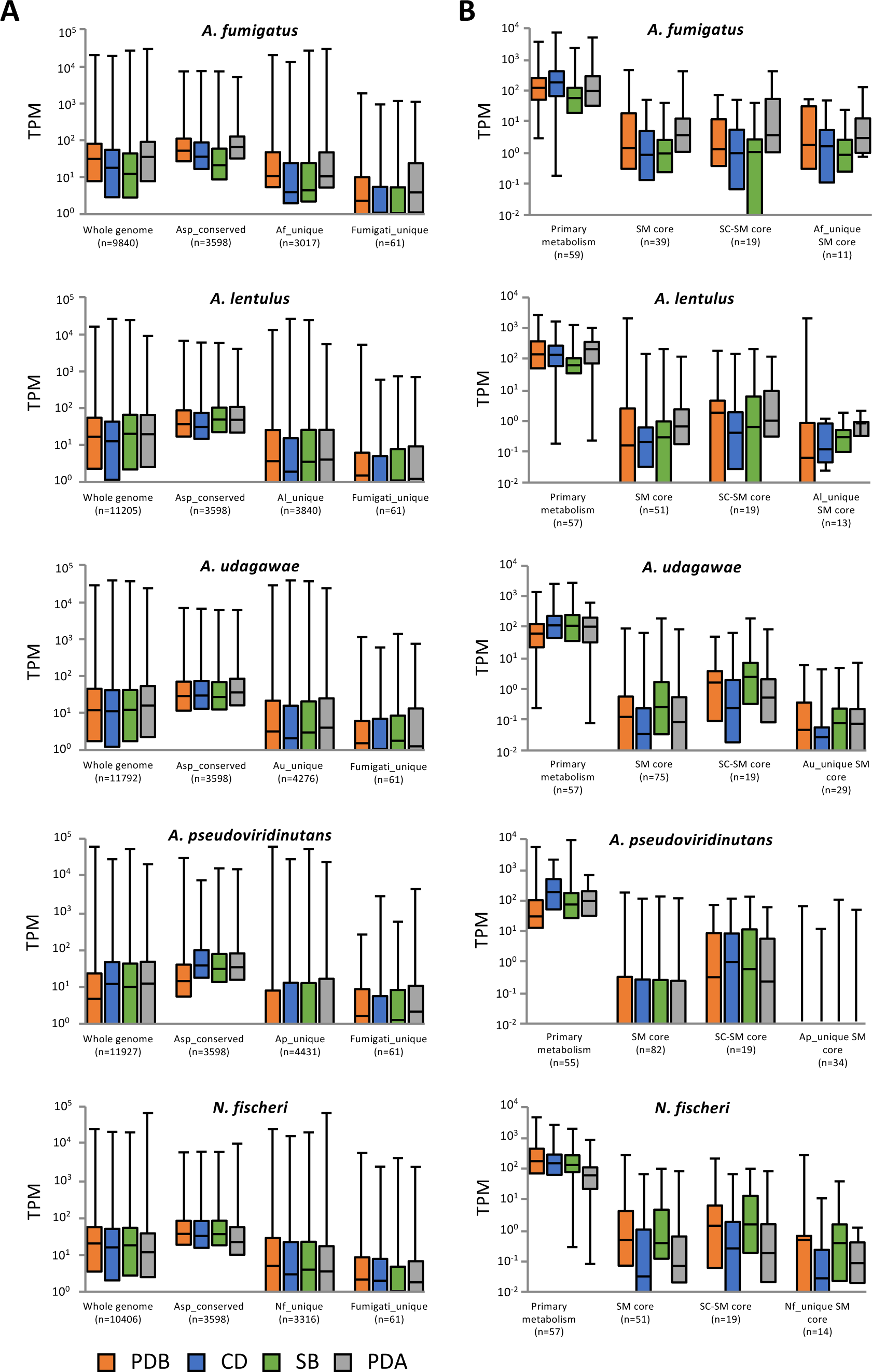
The distribution of the gene expression levels as assessed by the transcriptome analysis is shown using a box plot. (A) Gene expression patterns of the whole genome, *Asp*-conserved genes, species-specific genes, and *Fumigati*-unique genes under four different conditions (PDB, CD, SB, and PDA) are shown for each species. The *Fumigati*-unique genes are those present in the *Fumigati* species but not in other species, including *Aspergillus* fungi. (B) Expression patterns of genes involved in primary metabolism (glycolysis, TCA cycle, and the ergosterol biosynthesis pathway), SM core genes, SC SM core genes, and species-specific SM core genes are shown for each species. If the minimum values were below 10^0^ (A) or 10^−2^ (B), then the minimum plot is not shown under the box column.

Average expression levels for genes involved in primary metabolism, including glycolysis, the TCA cycle, and ergosterol biosynthesis, were relatively high in all the species and under all the tested conditions (Fig 3B). In contrast to the primary metabolic genes, the SM core genes were much less expressed in each species. The distributions of the expression levels were compared between SC and species-specific SM core genes. A set of species-specific SM core genes was found to be transcriptionally less active than SC SM core genes in *A. lentulus, A. udagawae, A. pseudoviridinutans*, and *N. fischeri* (Fig 3B).

When a gene with an expression level greater than 1/20^th^ of the mean TPM was considered as being expressed, 69.2% (27/39), 29.4% (15/51), 20% (15/75), 23.5% (19/81), and 19.6% (10/51) of the SM core genes were expressed under any of the conditions tested in *A. fumigatus, A. lentulus, A. udagawae, A. pseudoviridinutans*, and *N. fischeri*, respectively (Fig 4A). This data highlighted that the expression rates of SM core genes were low in these species, except in *A. fumigatus*. Notably, 36.8%–52.6% of the SC SM genes were expressed in *A. lentulus, A. udagawae, A. pseudoviridinutans*, and *N. fischeri* (Fig 4B), whereas only small percentages (6.7% to 17.6%) of the species-specific SM genes were expressed under the same conditions (Fig 4C). Interestingly, most *A. fumigatus* species-specific SM genes (8/11) were expressed under any of the tested conditions.

**Fig 4.**
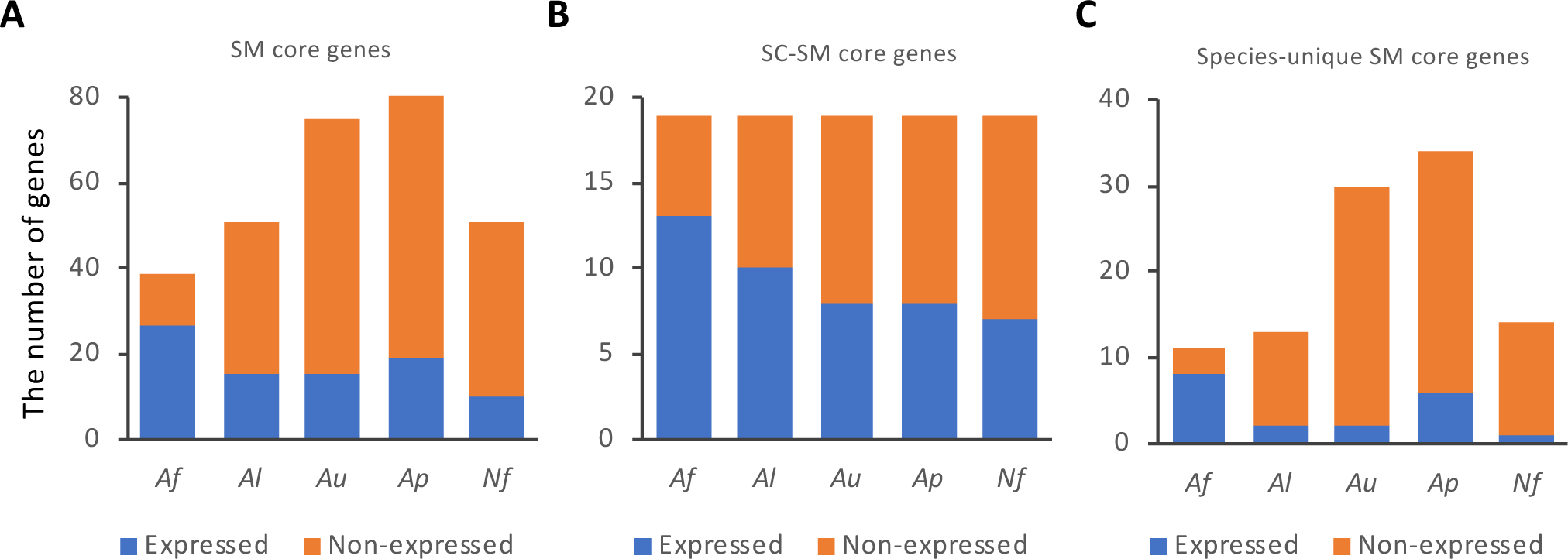
The numbers of expressed and non-expressed SM core genes. Genes with TPMs higher than 1/20^th^ of the mean TPM under either conditions were regarded as expressed genes, while the remaining were non-expressed genes. The numbers of expressed and non-expressed (A) SM core genes, (B) SC SM core genes, and (C) species-specific SM core genes are shown for each species. *Af*: *A. fumigatus*; *Nf*: *N. fischeri*; *Al*: *A. lentulus*; *Au*: *A. udagawae*; *Ap*: *A. pseudoviridinutans*.

### Variation in expression patterns of SC BGCs across the species

To investigate how the SC BGCs were transcriptionally regulated in the closely related species, we first sought to identify gene clusters whose expression levels were coordinately regulated under specific condition(s) using a MIDDAS-M program (33). Consequently, 4, 3, 5, 6, and 6 clusters that contained one or multiple SM core genes were detected to be coordinately expressed in *A. fumigatus, A. lentulus, A. udagawae, A pseudoviridinutans*, and *N. fischeri*, respectively (S2 Table). Among them, 3, 2, 3, 3, and 3 clusters were SC BGCs, whereas 0, 1, 1, 2, and 0 clusters were species-specific BGC, respectively. Notably, SC BGC7 (*hasD*) was detected to be coordinately expressed in all five species. The expression levels of the component genes in all the SC BGCs were depicted using a heat map (Fig 5). This highlighted that SC BGC4 (*pksP*) was exclusively expressed in *A. fumigatus* and *A. pseudoviridinutans*, and that SC BGC9 (*tpcC*) was only expressed in *A. fumigatus* on PDA. The SC BGC14 (*fccA*) was expressed in *A. pseudoviridinutans*, although its expressions levels were low in the other species. To identify the BGCs whose expressions were regulated in similar manners, a correlation analysis was performed among the five species for each BGC (Fig 6). High correlations among the five species were found for three SC BGCs (BGC1, −12, and −15), whereas there were no apparent correlations between any combinations of the species for BGC8, −10, and −14. Thus, three BGCs were differentially expressed across the species, while three others were regulated in a similar manner.

**Fig 5.**
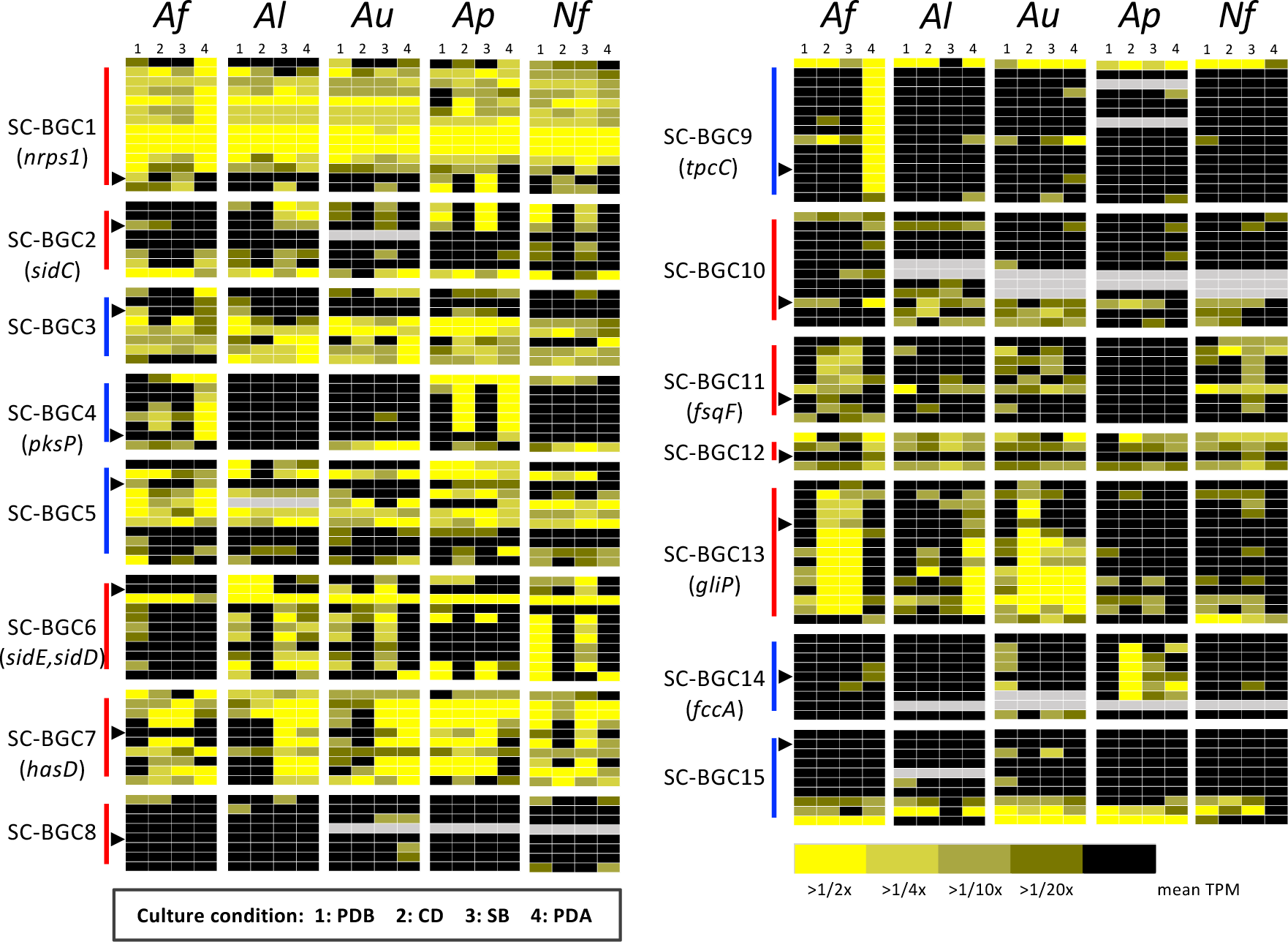
Heat map revealing expression profiles in the SC BGCs. The colors of bars between the BGC IDs and panels indicate the types of SM core genes, as follows: red: *NRPS* or *NRPS-like*; blue: *PKS* or *PKS-like*. A black triangle indicates SM core gene expression in the BGC. Grey panels indicate the absence of the corresponding gene. *Af*: *A. fumigatus*; *Nf*: *N. fischeri*; *Al*: *A. lentulus*; *Au*: *A. udagawae*; *Ap*: *A. pseudoviridinutans*.

**Fig 6.**
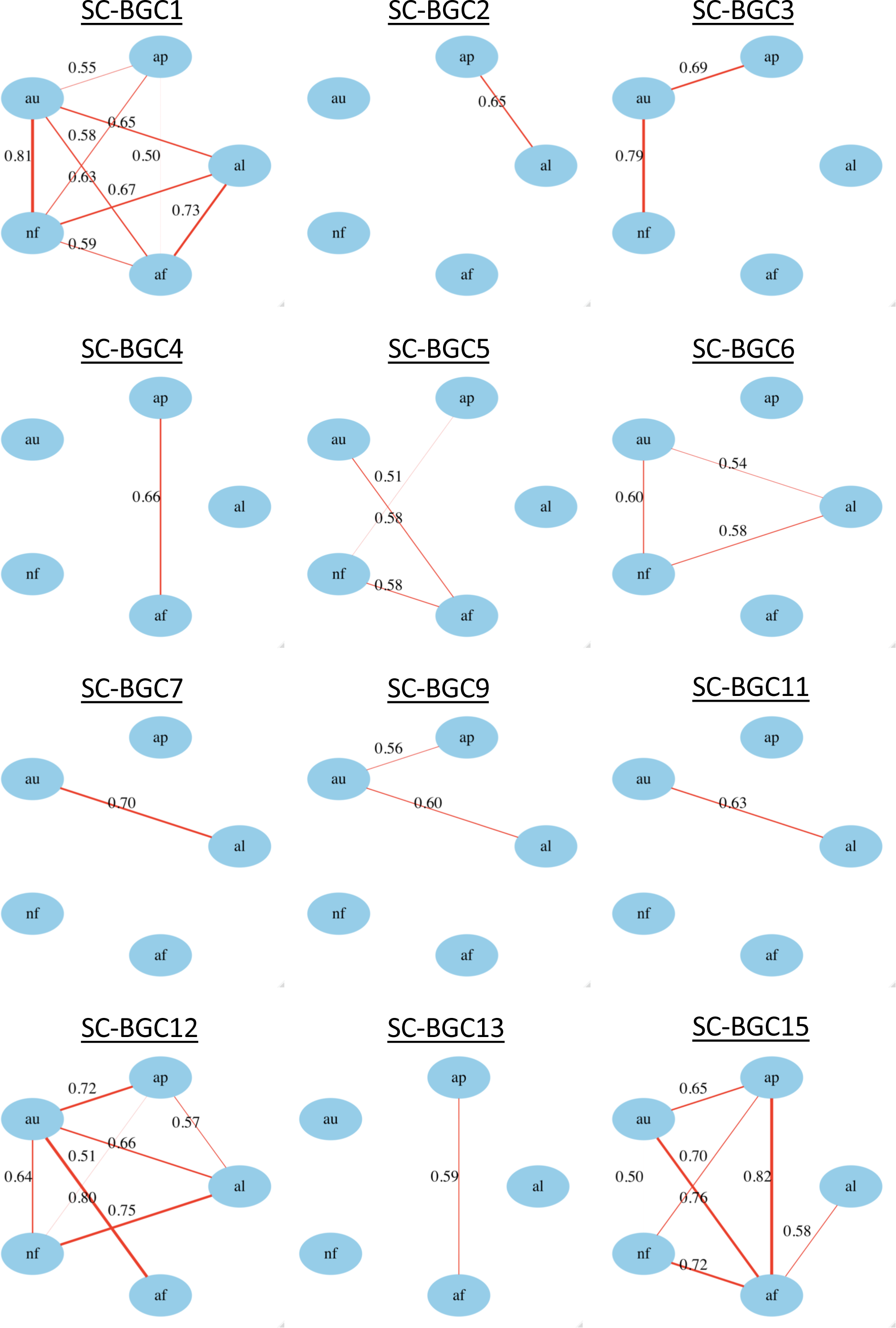
Correlations of gene expression patterns in the SC BGCs between species. Pearson’s correlation coefficients (PCCs) of the gene expressions (log2 TPM) were calculated in each SC BGC using the TPM values of all the component genes under the four different culture conditions. The relationships with PCC *r* ≥0.5 are indicated with red lines, with the line thickness corresponding to the magnitude of the correlation. There were no correlations between any pairwise combinations of SC BGC8, −10, and −14, among the species; therefore, they are not shown. *af*: *A. fumigatus*; *nf*: *N. fischeri*; *al*: *A. lentulus*; *au*: *A. udagawae*; *ap*: *A. pseudoviridinutans*.

### Comparative *in silico cis*-element analysis of SC BGCs

To gain a deeper insights into interspecies variations in the transcriptional regulation of SM gene clusters, we computationally searched DNA-binding sites for cluster-specific transcription factors (TFs). In total, 9 of the 15 SC BGCs included one or two putative TFs, which could act as cluster-specific transcriptional regulators (Table 1, S1 Fig). Notably, eight of the nine SC BGCs contain a C6-type TF. Thus, we focused on the SC BGCs (BGC7–14) harboring a C6-type TF to investigate whether there are potent DNA-binding sites in the promoter regions of each component gene in a cluster, and whether and how they are conserved among the closely related species. Because C6-type TFs reportedly bind to inverted CGG triplets spaced with several nucleotides, like CGG(N_x_)CCG, we sought to identify such bipartite motifs conserved in the promoter regions of component genes of the clusters using BioProspector (34) (S2A–H Fig). When the gap length was set at 3 bp, a CGG triplet was found in SC BGC13 (*gli*) and SC BGC14 (*nsc*). Notably, the palindromic sequence TCGG(N_3_)CCGA was found in SC BGC13, whereas SC BGC14 contains the non-palindromic sequence TCGG(N_3_)TTT(G/A), which is likely to be a variant of the inverted CGG triplet. The consensus sequence of each cluster was highly conserved among the five closely related species. When the gap was set variably from 1 to 10 nucleotides, CGG triplets were detected in SC BGC9 (*tpc*) and SC BGC11 (*fsq*) as well. However, these sequences were only partially conserved among the species. Thus, interspecies-conserved *cis*-elements for C6-type TFs were detected only in SCBGC13 out of the eight BGCs tested.

### Diverse sequences in the promoter regions of *gli* genes among the related species

The SC BGC13 (*gliP*), comprising 13 genes, is responsible for gliotoxin production and is well studied in *A. fumigatus* (28,35,36). The *gli* cluster is regulated by GliZ, a C6-type TF, and the DNA-binding site has been proposed as TCGG(N_3_)CCGA (37), which is identical to the palindromic sequence motif we detected. The palindromic sequence was found in 9 of 13 *gli* genes with perfect matches, and they are positioned approximately 100-bp upstream of the translational initiation site in each gene (Fig 7). Interestingly, although the positions of the consensus motifs are conserved in the five related species, sequences of the proximal regions are diverse, particularly in *gliL, gliM, gliG*, and *gliN* (S3A–M Fig). These variations in the regions near the *cis*-element may affect the transcriptional regulators’ access. These results suggested that sequence diversification had frequently occurred in the promoter region during the course of evolution, which might result in variations in the expression patterns of fungal SM BGCs.

**Fig 7.**
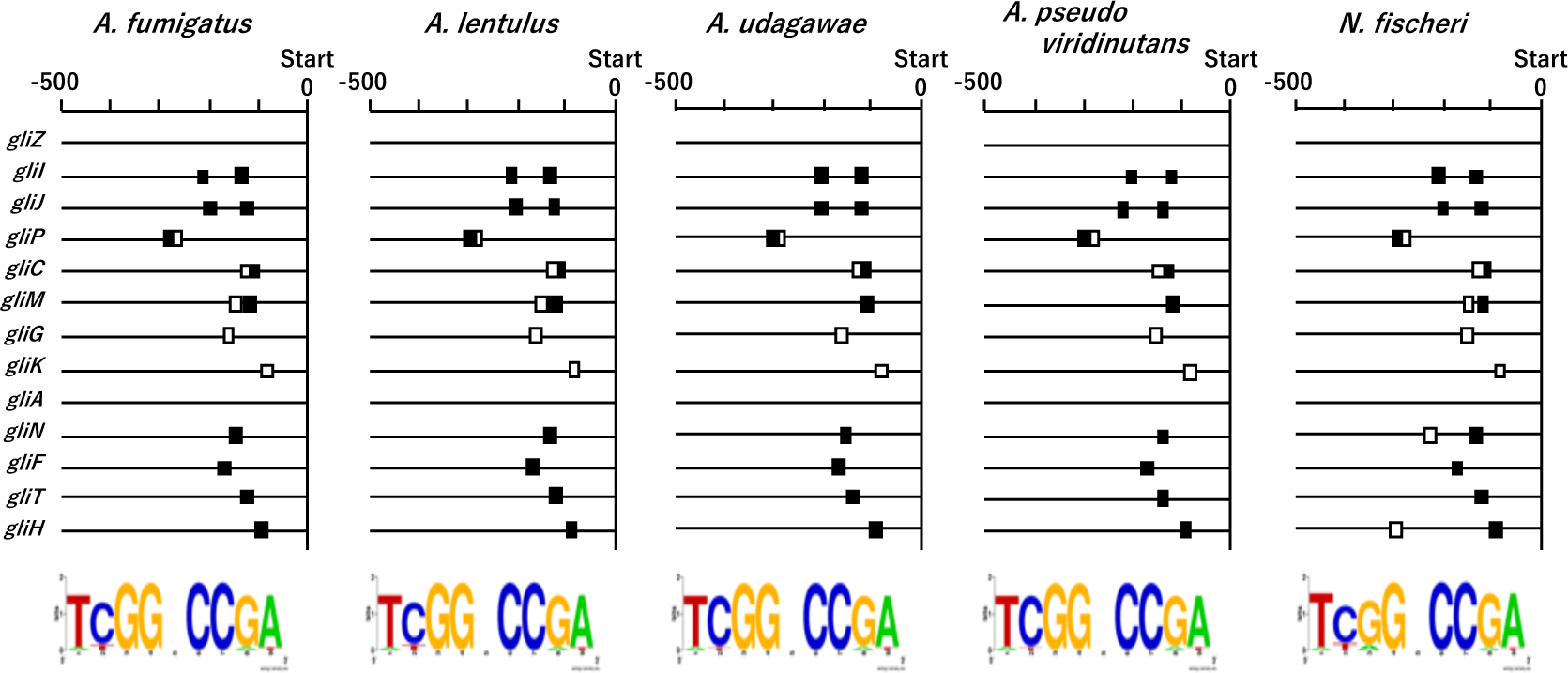
The consensus motifs in the promoter regions of the *gli* genes. The consensus motifs were detected in the 13 *gli* genes in each species using BioProspector (Release 2), and representative motifs are shown under the positioning map. Positions of the primary motif detected (<u>TCGG</u>NNN<u>CCGA</u>) is indicated by black boxes and the others by white boxes. –500, 500-bp upstream from the translational initiation site of each gene.

## Discussion

Since the first fungal genomes were published, the potential of fungi to produce a wide variety of secondary metabolites has become generally accepted. Comparative genomics studies regarding fungal SMs have been intensively conducted during this decade, revealing the genetic diversity, universality, and plasticity of SM gene clusters in fungal genomes (5,6,9,10,38,39). Variations in SMs produced by fungi allow their metabolites to be used for medical and biotechnological applications.

In the present study, we *de novo* sequenced the genome of *A. pseudoviridinutans* and compared genomes of five species of section *Fumigati*. The interspecies comparisons allowed us to determine the SM gene clusters that are conserved across the species (SC) and that are unique to the species (species-specific). In *A. fumigatus*, there are 19 SC SM core genes, more than half of which had been identified as encoding metabolites, including well-studied siderophores, DHN-melanin, and gliotoxin. In contrast, only 3 of 11 species-specific SM core genes had been characterized. The most recently identified gene, *Afu1g01010*, is involved in fumigermin biosynthesis in strain ATCC 46645, but not in strain Af293 owing to multiple SNPs inside the ORF (13). On the basis of the results, we hypothesized that SC BGCs are transcriptionally more active and functional than species-specific BGCs, enabling us to easily access and preferentially characterize the metabolites derived from the SC BGCs under laboratory conditions. However, the transcriptome data revealed that both sets of SM core genes were expressed at comparable rates of ∼70% in *A. fumigatus* (Fig 4B, C). In contrast to *A. fumigatus*, the rates (6.7% to 17.6%) and magnitudes of expressed species-specific SM core genes were low in the other four species (Figs 3B and 4C). These data suggested that the narrowly distributed SM genes tended to be less preferentially expressed. During the course of evolution, transcriptional regulation may be a potential target for the degeneration of secondary metabolism, although further clarification is required.

Here, the expression patterns of SC BGCs in each species were found to be varied across species despite the highly conserved gene contents and orders (S1 Fig). For example, the SC BGC6, which is responsible for siderophore production, was highly expressed in *N. fischeri* in PDB, whereas the expression was lower or partial in *A. fumigatus* and *A. pseudoviridinutans* (Fig 5). The SC BGC14, which was identified as a fumicycline producing cluster, was expressed in *A. pseudoviridinutans* on CD medium, while that of *A. fumigatus* was hardly expressed under any conditions. This BGC14 is remarkably induced in *A. fumigatus* when cocultured with *Streptomyces rapamycinicus*, resulting in the production of fumicycline (32). Given that *A. pseudoviridinutans* produces this metabolite in mono-cultures of CD, the molecular mechanisms underlying the transcriptional activation might be different between the two species. High-level expressions of *pks* and *tpc* clusters were observed in *A. fumigatus* on PDA but not in PDB, CD, and SB liquid media. The metabolites DHN-melanin and trypacidin accumulate in the conidia, which was consistent with the expression profiles determined in the different cultivation styles (41,42). Notably, the *A. lentulus, A. udagawae*, and *N. fischeri* strains tested here were unable to produce many conidia on PDA or in liquid media, which accounted for the lack of *pks* and *tpc* cluster expression in the fungi. In contrast, a moderate level of conidiation was observed in *A. pseudoviridinutans* on PDA, which could explain the high-level expression of the *pks* cluster in the fungus on PDA. However, the *tpc* cluster was not expressed on PDA; therefore, it could be assumed that the *tpc* cluster of *A. pseudoviridinutans* had become transcriptionally silent after the divergence from *A. fumigatus*. The *gli* cluster was highly expressed in *A. fumigatus* and *A. udagawae* in CD. However, the culture extract from *A. udagawae* contained no detectable gliotoxin, while it was highly produced by *A. fumigatus* (data not shown). It is possible that post-transcriptional regulation affects the production of the fungal metabolites in media.

The MIDDAS-M analysis revealed that the cluster covering *gene_01461*.*t1* to *gene_01473*.*t1*, which contains 13 genes including species-specific SM core genes *gene_01472*.*t1* and *gene_01473*.*t1*, in *A. lentulus* was highly expressed (S4A, B Fig). This cluster was orthologous to the terrein (*ter*) cluster of *Aspergillus terreus* in terms of gene content and order (S4A Fig). The flanking genes of the *ter* cluster are not conserved, suggesting that this cluster has been translocated between the genomes by horizontal gene transfer. Interestingly, *A. terreus* can produce terrein in PDB and PDA (42), and *A. lentulus* also produces large amounts of terrein under these conditions (S2C Fig). The consistent conditions required for terrein production suggest that the *ter* cluster is regulated in a similar manner in *A. lentulus* and *A. terreus*. This indicates that the fungal SM gene cluster has been evolutionarily transferred and that it retains its transcriptional regulatory mechanism in different hosts.

In *A. fumigatus*, the transcriptional regulation of the *gli* cluster has been studied, and GliZ, a C6-type TF, plays a pivotal role in cluster-wide regulation (37). We found that the promoter sequences of the *gli* genes were somehow diverse across the section *Fumigati* strains, despite the highly conserved consensus sequence. Variations in the sequences of the promoter regions might affect the efficiency of RNA polymerase binding to the region, the positioning of the transcription start point, and the appearance of an unsuitable initiation codon, which could consequently lead to changes in the expression of the *gli* cluster. In addition to the *gli* cluster, seven SC BGCs putatively contain a C6-type TF. Unexpectedly, no bipartite palindromic motifs, such as CGG(N_x_)CCG, were predicted using bioinformatics tools. The *has* (SC BGC7) and *fsq* (SC BGC11) clusters are responsible for the production of hexadehydro-astechrome and fumisoquins, respectively (26,27). In these reports, overexpression of the C6-type TFs (HasA and FsqA) resulted in transcriptional activation of the clusters. Therefore, further investigations of common *cis*-element among genes in the clusters are needed to determine whether the TFs regulate the clusters in a manner identical to that of the *A. fumigatus* cluster.

Our genomic study provides a comprehensive catalog of SM genes in *A. lentulus, A. udagawae*, and *A. pseudoviridinutans*, which had not been adequately compiled previously, while those of *A. fumigatus* and *N. fischeri* have been well investigated. Larsen et al. reported auranthine, cyclopiazonic acid, neosartorin, pyripyropene A, and terrein as major metabolites of *A. lentulus* (43), which was supported by the presence of the corresponding core genes (*cpaA, nsrB, pyr2*, and *terAB*, respectively) in the genome. On the basis of the SM gene list, 18 of 39 SM core genes in *A. fumigatus* were identified as being involved in the production of known metabolites. With the exception of these genes, as well as *cpaA, nsrB, pyr2*, and *terAB*, no SM core genes have been assigned as encoding known metabolites in the other *Fumigati* strains tested here. Because some of the unstudied SM genes were highly expressed under laboratory conditions, it is still possible to characterize genes and identify novel metabolites from these genomes.

In conclusion, we combined comparative genomic and transcriptomic analyses to study variations in transcriptional activities of fungal BGCs across closely related species. This research provides a perspective on how the BGC distribution may be correlated with the tendency for silent secondary metabolism production. The transcriptional regulatory pattern for common BGCs could differ even among closely related species. On the basis of our findings, we proposed that the diversification of transcriptional regulation could drive the evolution and degeneration of SM gene clusters. Further efforts to characterize such transcriptional diversity will expand our understanding of the evolutionary processes affecting fungal secondary metabolism.

## Materials and Methods

### Fungal strains

The strains *A. fumigatus* Af293, *A. lentulus* IFM 54703, *A. udagawae* IFM 46973, *A. pseudoviridinutans* IFM 55266, and *N. fischeri* NRRL 181 were provided through the National Bio-Resource Project, Japan (http://www.nbrp.jp/) and are preserved at the Medical Mycology Research Center, Chiba University. The genomes of *A. lentulus* IFM 54703 (19) and *A. udagawae* IFM 46973 (20) were previously sequenced, and the data were retrieved from the NCBI database (https://www.ncbi.nlm.nih.gov/). The strain *A. pseudoviridinutans* IFM 55266 was isolated from a patient in Japan and identified using tubulin and calmodulin partial sequences (22).

### Culture conditions

All the strains were grown in liquid PDB (BD Difco, Franklin Lakes, NJ, USA), SB (BD Difco), and CD (BD Difco) at 37°C for 5 d by inoculating each culture with three 0.5-cm^2^ agar plugs. For asexual stage culturing, the mycelia, which were cultured in PDB at 37°C for 3 d, were harvested using a miracloth, washed with distilled water, and then placed onto PDA plates (BD Difco) for another 2 d of culturing at 37°C.

### Molecular phylogenetic analysis

A phylogenetic tree of *A. fumigatus* and related species was constructed using partial β-tubulin and calmodulin gene sequences. The strains and sequence identification numbers used for the analysis are listed in S3 Table. The sequence alignments and phylogenetic tree construction based on a neighbor-joining analysis (44) were performed using Clustal X software (45). The distances between sequences were calculated using Kimura’s two-parameter model (46). A bootstrap was conducted with 1,000 replications (47). A genome synteny analysis was conducted as described previously (6).

### Genome sequencing and gene prediction for *A. pseudoviridinutans*

The genomic DNA of *A. pseudoviridinutans* was extracted from a 2-d-old culture using phenol-chloroform and NucleoBond buffer set III (TaKaRa, Shiga, Japan). The DNA was fragmented in an S2 sonicator (Covaris, MA, USA), and then purified using a QIAquick gel extraction kit (Qiagen, CA, USA). A paired-end library with insert sizes of 700 bp was performed using a NEBNext Ultra DNA library prep kit (New England BioLabs, MA, USA) and NEBNext multiplex oligos (New England BioLabs) in accordance with the manufacturer’s instructions. Mate-paired libraries with insert sizes of 3.5 to 4.5 kb, 5 to 7 kb, and 8 to 11 kb were generated using the gel selection-based protocol of the Nextera mate pair kit (Illumina, San Diego, CA, USA) and a 0.6% agarose gel in accordance with the manufacturer’s instructions. The quality of the libraries was determined by an Agilent 2100 Bioanalyzer (Agilent Technologies, Santa Clara, CA, USA). The 100-bp paired-end sequencing was performed by Hiseq 1500 (Illumina) using the HiSeq reagent kit v1, in accordance with the manufacturer’s instructions.

The Illumina reads sets were trimmed using Trimmomatic (ver. 0.33), and sequencing adapters and sequences with low-quality scores were removed (48). The read sets were then assembled using Platanus (ver. 1.2.1) (49). Protein-coding genes of *A. pseudoviridinutans, A. lentulus*, and *A. udagawae* were predicted using the FunGap pipeline (50).

### Identification of SM core genes

To identify the *NRPS* and *PKS* genes of *A. lentulus, A. udagawae, A. pseudoviridinutans*, and *N. fischeri*, a set of proteins for each strain were queried using the BLASTP program against multiple *Aspergillus* genome data available in AspGD (http://www.aspgd.org/). The proteins showing high similarity levels to the known *A. fumigatus* NRPS or PKS were considered for further verification. The amino acid sequences of the candidate proteins were manually aligned with the sequence of the authentic NRPS or PKS. The motifs were confirmed using PKS/NRPS Analysis Web-site (http://nrps.igs.umaryland.edu/) programs. The orthologous relationships were determined using the BLASTP program with the criteria of more than 80% identities and more than 80% coverage of either protein sequence. A cladogram was constructed based on a binary matrix (presence/absence of the PKSs and NRPSs) using Cluster 3.0 (http://bonsai.hgc.jp/~mdehoon/software/cluster/software.htm#ctv). The tree and heat map were constructed using Tree View (51).

### RNA sequencing (RNA-seq) and data analysis

Each strain was cultured, harvested, and ground into a fine powder using a mortar and pestle. Total RNA was isolated using Sepazol-RNA Super G (Nacalai, Kyoto, Japan) in accordance with the manufacturer’s instructions. The RNA isolation was carried out with two biological replicates, and they were pooled for preparation of the RNA-seq libraries. The RNA-seq libraries were constructed using a KAPA mRNA Hyper Prep Kit (Nippon Genetics, Tokyo, Japan), in which mRNA was purified by poly-A selection, the second-strand cDNA was synthesized from the mRNA, the cDNA ends were blunted and polyAs added at the 3′ ends, and appropriate indexes were ligated to the ends. The libraries were PCR amplified, and the quantity and quality were assessed using a Bioanalyzer (Agilent Technologies). Each pooled library was sequenced using Illumina Hiseq 1500. The gene expression levels were estimated using a previously described method (52). Briefly, the sequencing reads cleaned by Trimmomatic (48) were mapped to the reference genomes using STAR (ver. 2.4.2a) (53). A raw read count was conducted using HTSeq (ver. 0.5.3p3) (54), and transcript abundances were estimated as TPMs (55). The expressed genes of PKSs, NRPSs, and TSs were identified using the 1/20^th^ mean TPM criterion.

### MIDDAS-M analysis

Binary logarithms of RPKM values generated under the four culture conditions were analyzed using the MIDDAS-M algorithm to detect gene clusters in which certain condition(s) coordinately expressed or repressed the genes (33). Briefly, the induction ratio of each gene was evaluated for every pairwise combination of the four culture conditions. After *Z*-score normalization, the gene cluster expression scores were calculated for each gene using the algorithm. The maximum cluster size was set as 30. The threshold to detect clusters was set to the value corresponding to false positive rate of 0, which was evaluated from data in which the original gene order was randomly shuffled. The threshold values were 1.7E5, 2.5E5, 1.9E05, 5.9E5, and 4.6E5 in *A. fumigatus, A. lentulus, A. udagawae, A. pseudoviridinutans*, and *N. fischeri*, respectively.

### SC BGC expression level correlations among species

The SC BGC gene expression pattern correlations between species were evaluated using Pearson’s correlation coefficients (PCCs) of the gene expressions (log2 TPM) and the R programming language (56). To calculate the PCC in each SC BGC, the TPM values of all the component genes under the four different conditions were used. The relationships having PCC *r* ≥0.5 were visualized using DiagrammeR (ver. 1.0.5) (57).

### Extraction of compounds and HPLC analysis

For terrein detection, 5 μL of culture supernatant was subjected to HPLC analysis, which was performed using an Infinity1260 modular system (Agilent Technologies) consisting of an autosampler, high-pressure pumps, a column oven, and a photo diode array detector with InfinityLab Poroshell 120 EC-C18 column (particle size: 2.7 μM; length: 100 mm; internal diameter: 3.0 mm) (Agilent Technologies). Running conditions were as follows: gradient elution, 5%–100% acetonitrile in water over 30 min; flow rate, 0.8 mL min^-1^; detection wavelength; 254 nm. Terrein production was identified by comparing retention times and UV spectra with those of an authentic standard purchased from Cayman Chemical Company (Ann Arbor, MI, USA).

## Data availability

The whole-genome sequences of *A. pseudoviridinutans* IFM 55266 have been deposited at DDBJ/EMBL/GenBank under the accession numbers BHVY01000001–440. The raw RNA-seq data have been submitted to the DDBJ Short Read Archive under accession number PRJDB7496.

## Author contributions

HT, MU, and DH designed the research; HT, MU, AN, MS, YK, SU, TY, and DH performed experiments; HT, AW, KK, and TY contributed new materials/tools; HT, MU, AN, MS, YK, SU, TY, and DH analyzed data; and HT, MU, AN, MS, TY, and DH wrote the manuscript.

## ACKNOWLEDGMENTS

This study was supported by the National Bioresource Project (to HT and TY), by AMED under grant numbers JP19fm0208024 (to HT, AW, and DH) and 19jm0110015 (to HT, AW, KK, TY, and DH), and by a grant from the Institute for Fermentation, Osaka (to DH). We would like to thank Dr. Atsushi Iwama, Dr. Motohiko Ohshima, and Dr. Atsunori Saraya (Chiba University) for technical support with the Illumina HiSeq 1500, and Dr. Teigo Asai (The University of Tokyo) for fruitful discussions on potential metabolites in the strains. We thank Lesley Benyon, PhD, from Edanz Group (www.edanzediting.com/ac) for editing a draft of this manuscript.

## Supporting information

**S1 Fig**. Comparison of SC-BGCs among the *Fumigati* species. The genes predicted to be cluster components are indicated with blue arrows, while those indicated with white arrows are outside the cluster. Orange arrows indicate SM core genes, and red arrows indicate transcription factors. Black arrows indicate genes that show identities lower than 80% compared with the corresponding gene in *A. fumigatus*.

**S2 Fig**. The conserved motifs in the SC BGCs in each species identified using BioProspector (Release 2). *Af*: *A. fumigatus*; *Nf*: *N. fischeri*; *Al*: *A. lentulus*; *Au*: *A. udagawae*; *Ap*: *A. pseudoviridinutans*.

**S3 Fig**. Alignments of promoter sequences. The sequences 500-bp upstream from initiation sites for each *gli* gene were aligned using Clustal X. The estimated consensus motifs are indicated by red characters, and the positions are indicated by boxes. If another gene in the sequence has a coding region, then the region is indicated by a dashed blue-lined box. The direction of gene transcription is indicated by a blue arrow. The sequences’ ATG sites are shown in white characters. *Af*: *A. fumigatus*; *Nf*: *N. fischeri*; *Al*: *A. lentulus*; *Au*: *A. udagawae*; *Ap*: *A. pseudoviridinutans*.

**S4 Fig**. Characterization of the *ter* cluster of *A. lentulus*. (A) Structures of *ter* clusters for *A. lentulus* and *A. terreus*. The SM core genes are indicated by orange arrows. The genes (*terG, terH, terI*, and *terJ*) whose involvement in terrein production remains obscure in *A. terreus* are indicated by light grey arrows. (B) The expression profiles of *ter* genes in *A. lentulus* are shown using a heat map. (C) The production of terrein in *A. lentulus*. The strains were cultivated in PDB, SB, CD, and PDA (Asex), and the ethyl acetate-derived culture extracts were analyzed using HPLC. Terrein (**1**) production was identified by comparison with the standard (Std.). The corresponding peaks are indicated by arrows, and the UV spectrum from a PDA culture is indicated by a yellow-lined box.

**S1 Table**. The secondary metabolite (SM) core genes

**S2 Table**. The coodinately expressed secondary metabolite (SM) gene clusters identified using the MIDDAS analysis

**S3 Table**. The sequences used to construct the phylogenetic tree

