## Supplementary material for "Comparative genome and transcriptome analyses revealing interspecies variations in the expression of fungal biosynthetic gene clusters": Fig S1

**A**

SC-BGC1

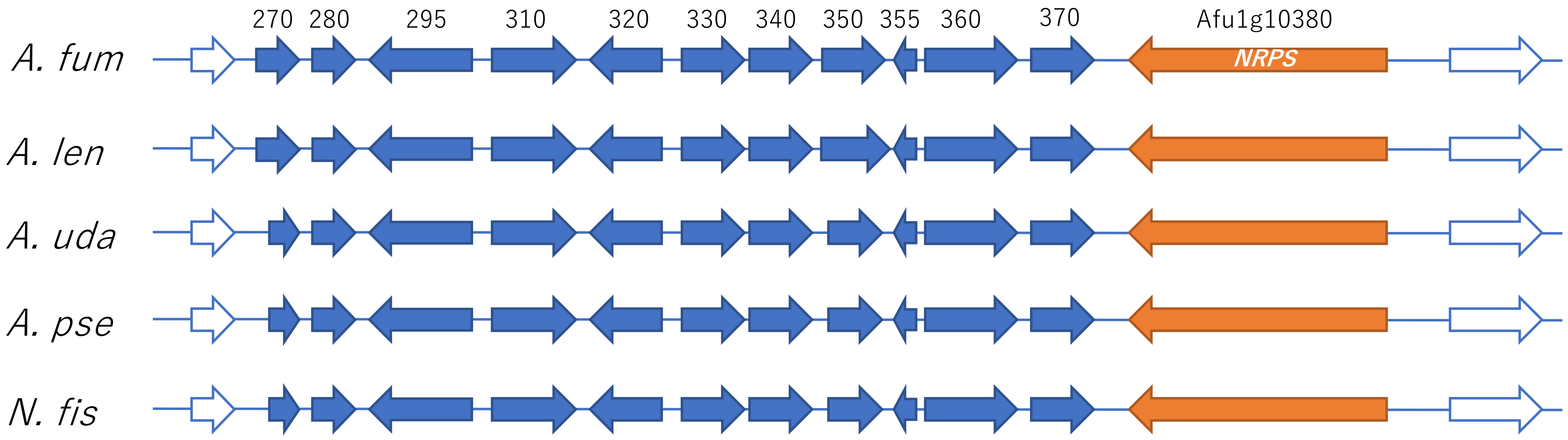
270
280
295
310
320
330
340
350
355
360
370
Afu1g10380
NRPS
A. fum
A. len
A. uda
A. pse
N. fis

**B**

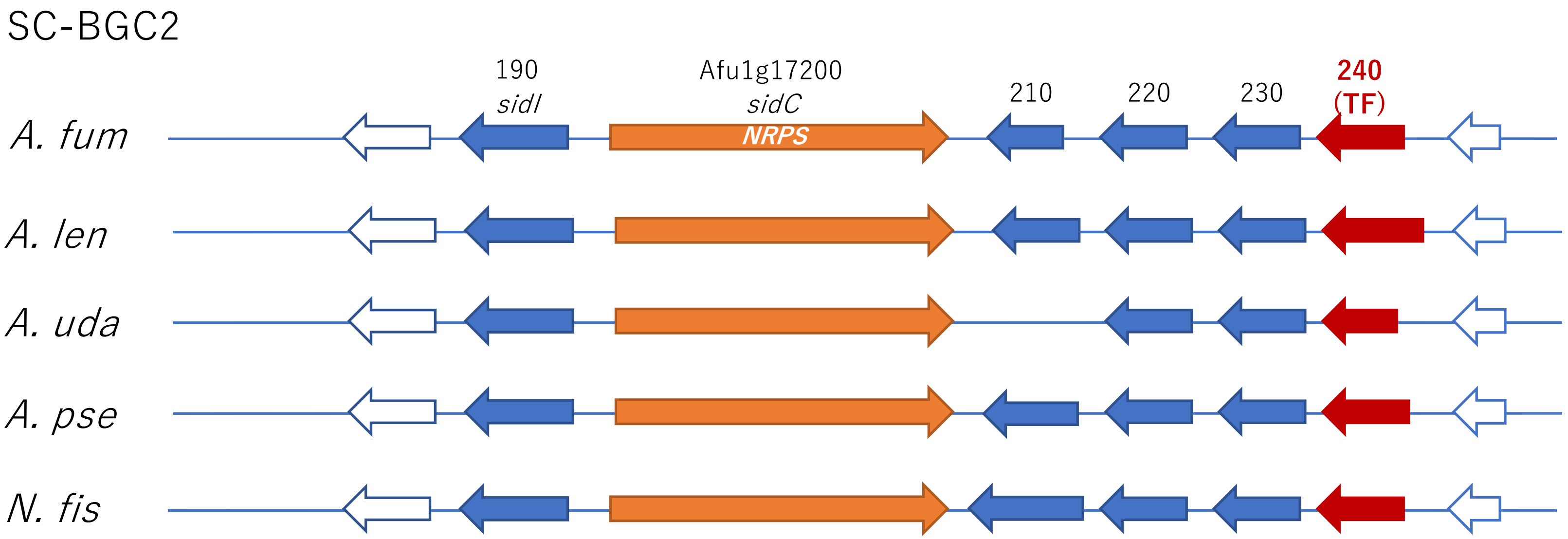
SC-BGC2
190
*sidI*
Afu1g17200
*sidC*
*NRPS*
210
220
230
240
(TF)
*A. fum*
*A. len*
*A. uda*
*A. pse*
*N. fis*

**C**

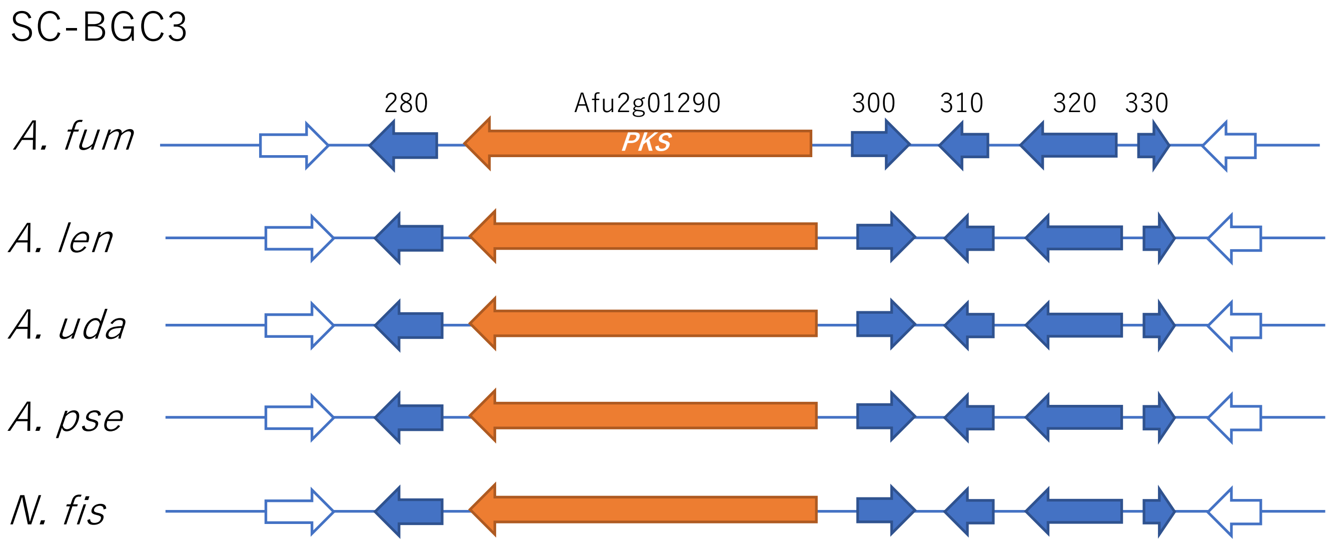
SC-BGC3
280
Afu2g01290
300
310
320
330
PKS
A. fum
A. len
A. uda
A. pse
N. fis

**D**

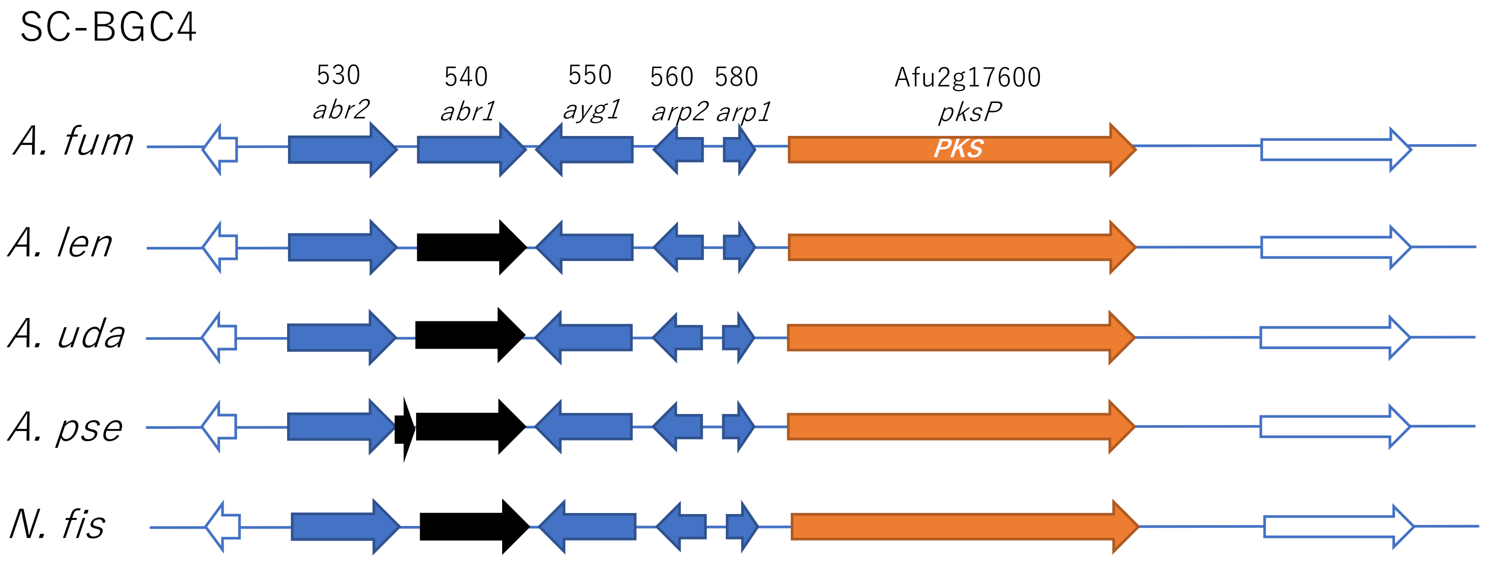
SC-BGC4
530
abr2
540
abr1
550
ayg1
560
arp2
580
arp1
Afu2g17600
pksP
PKS
A. fum
A. len
A. uda
A. pse
N. fis

**E**

SC-BGC5

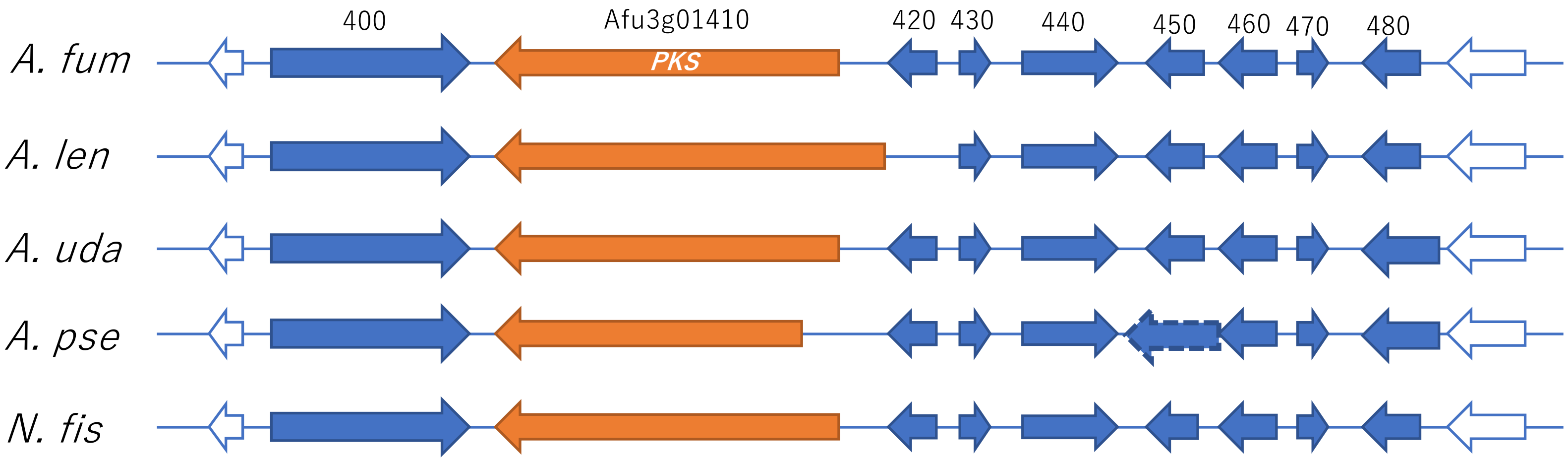
400
Afu3g01410
PKS
420
430
440
450
460
470
480
A. fum
A. len
A. uda
A. pse
N. fis

F

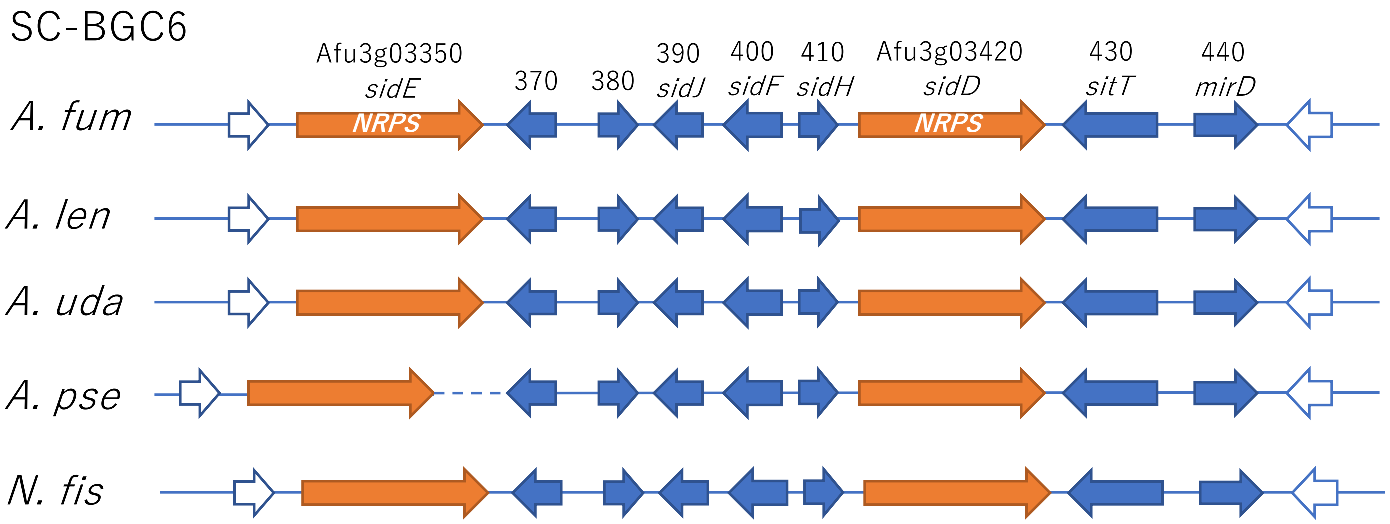
SC-BGC6
Afu3g03350
sidE
370
380
390
sidJ
400
sidF
410
sidH
Afu3g03420
sidD
430
sitT
440
mirD
A. fum
NRPS
NRPS
A. len
A. uda
A. pse
N. fis

**G**

SC-BGC7

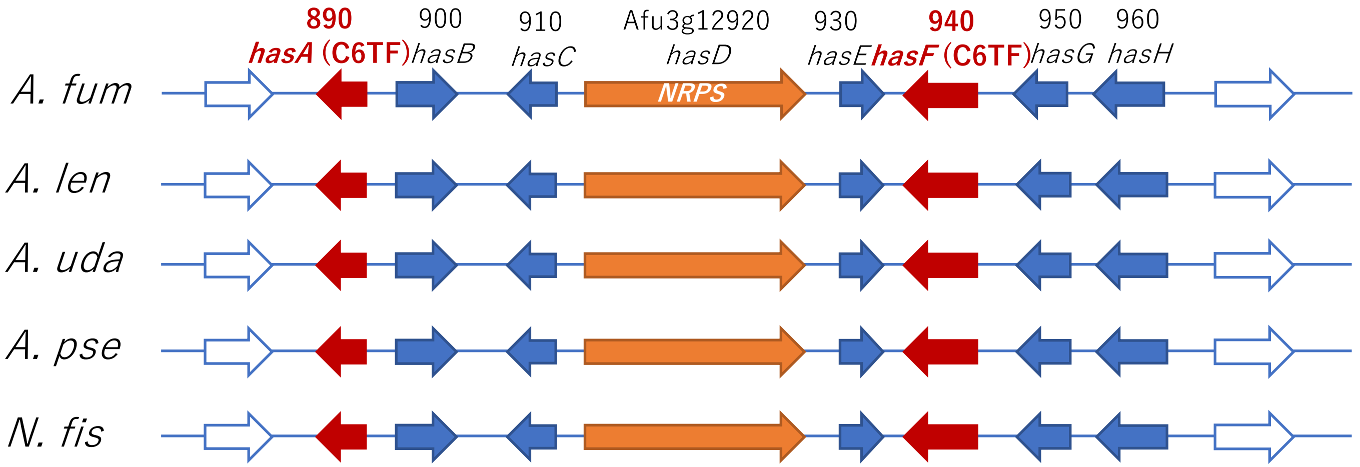
890
*hasA* (C6TF)
900
*hasB*
910
*hasC*
Afu3g12920
*hasD*
NRPS
930
*hasE*
940
*hasF* (C6TF)
950
*hasG*
960
*hasH*
*A. fum*
*A. len*
*A. uda*
*A. pse*
*N. fis*

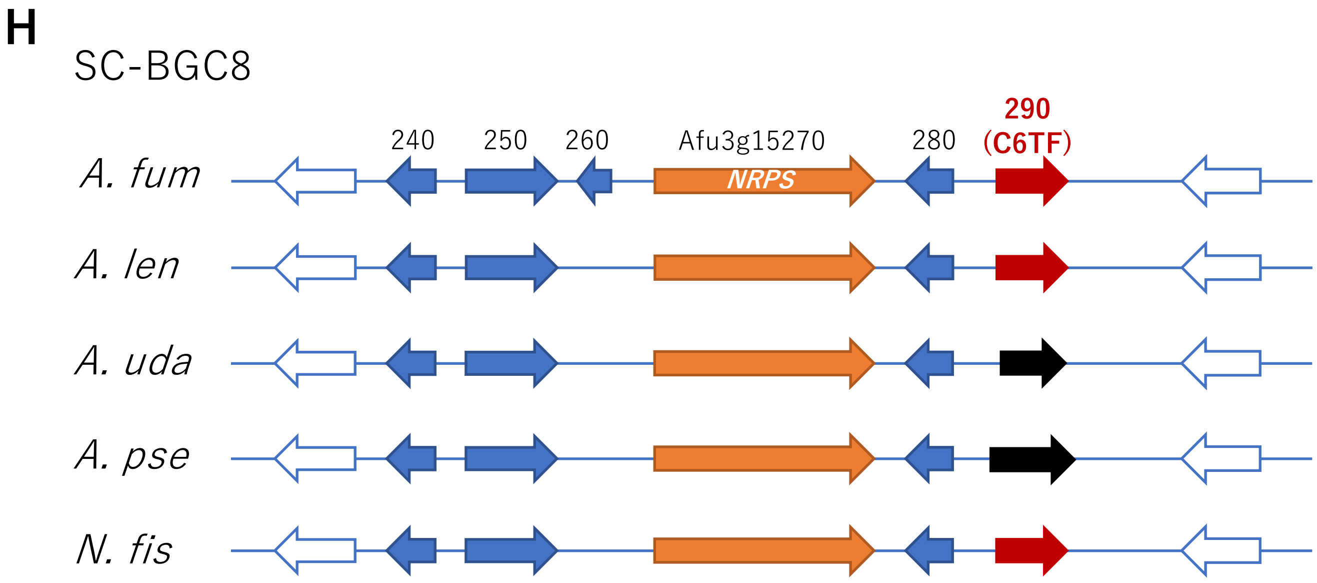
H
SC-BGC8
A. fum
A. len
A. uda
A. pse
N. fis
240
250
260
Afu3g15270
NRPS
280
290
(C6TF)

I

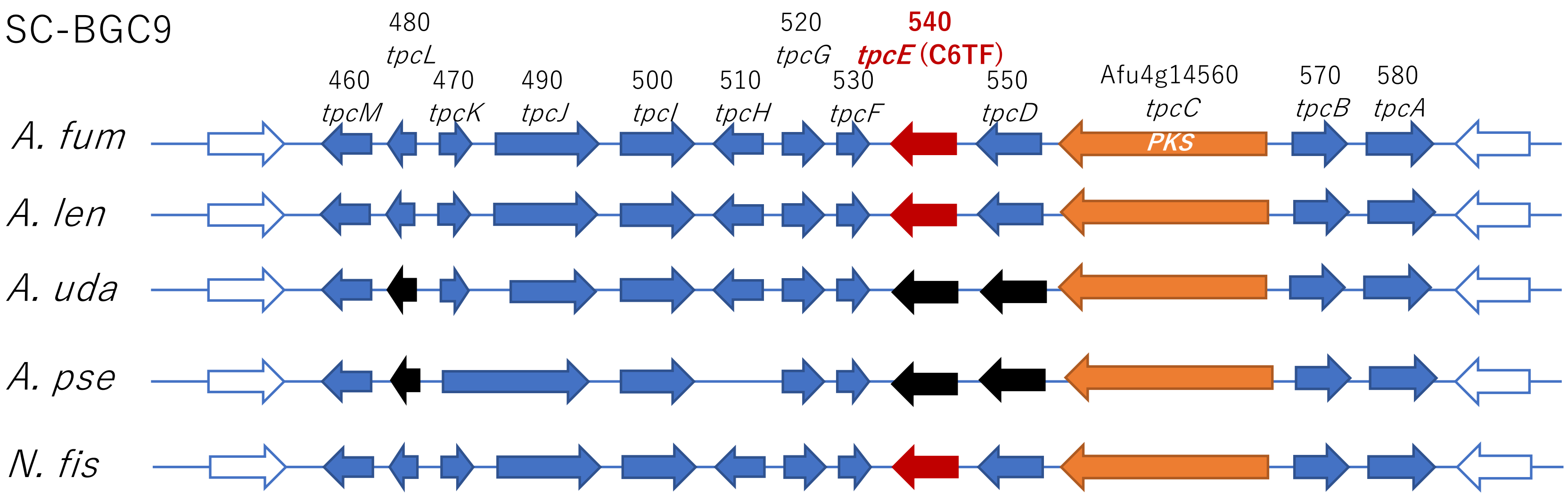

SC-BGC9
480
tpcL
520
tpcG
540
tpcE (C6TF)
460
tpcM
470
tpcK
490
tpcJ
500
tpcI
510
tpcH
530
tpcF
550
tpcD
Afu4g14560
tpcC
570
tpcB
580
tpcA
PKS
A. fum
A. len
A. uda
A. pse
N. fis

**J**

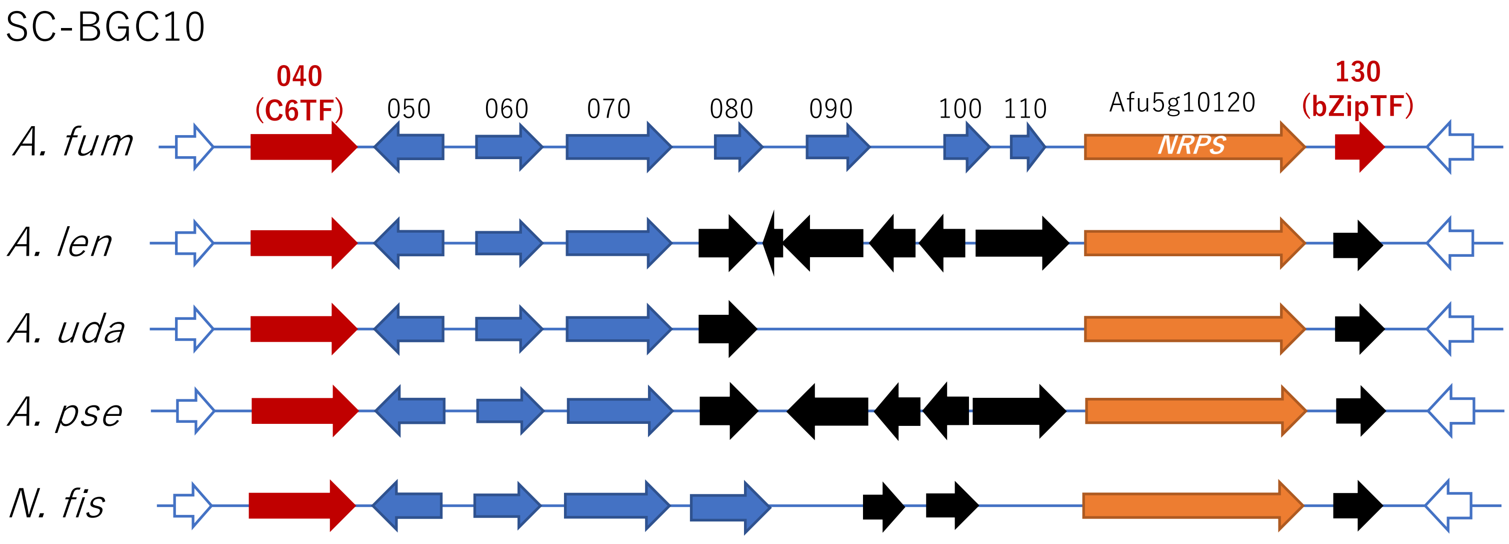
SC-BGC10
040
(C6TF)
050
060
070
080
090
100
110
Afu5g10120
NRPS
130
(bZipTF)
A. fum
A. len
A. uda
A. pse
N. fis

**K**

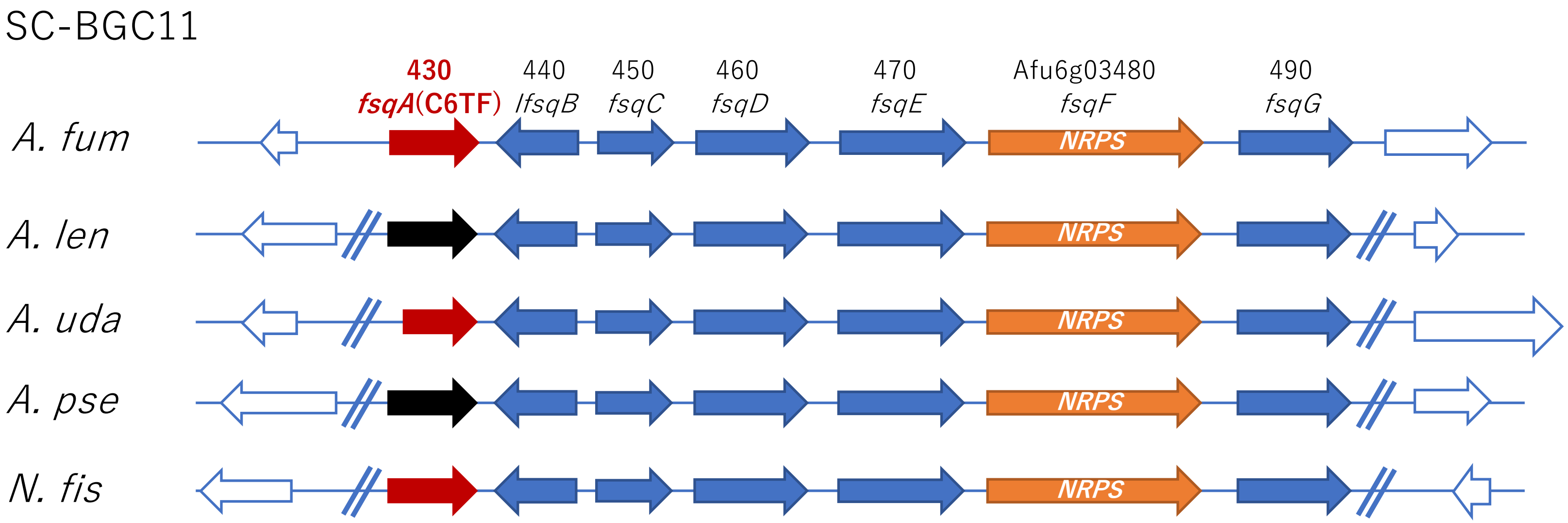

SC-BGC11
430
fsqA(C6TF)
440
IfsqB
450
fsqC
460
fsqD
470
fsqE
Afu6g03480
fsqF
490
fsqG
A. fum
NRPS
A. len
NRPS
A. uda
NRPS
A. pse
NRPS
N. fis
NRPS

**L**

SC-BGC12

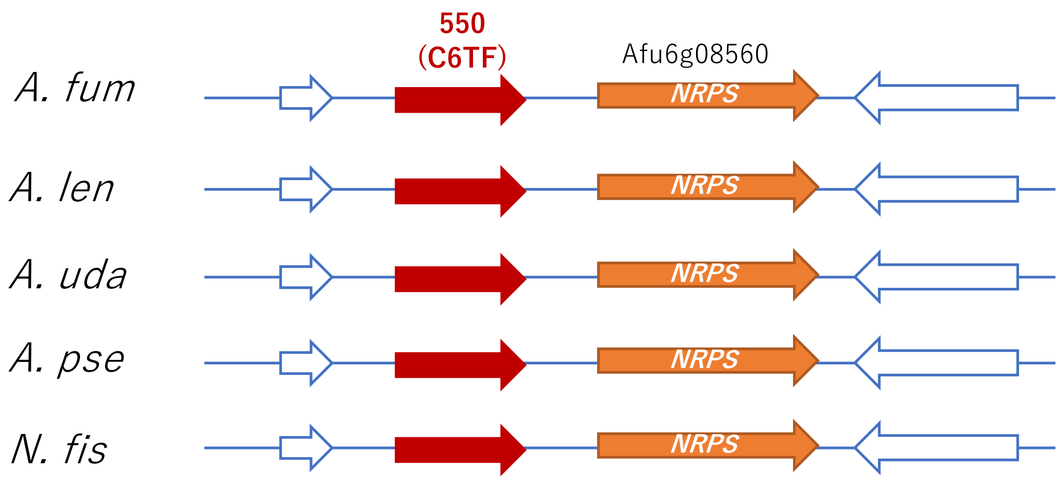

550
(C6TF)
Afu6g08560
A. fum
NRPS
A. len
NRPS
A. uda
NRPS
A. pse
NRPS
N. fis
NRPS

**M**

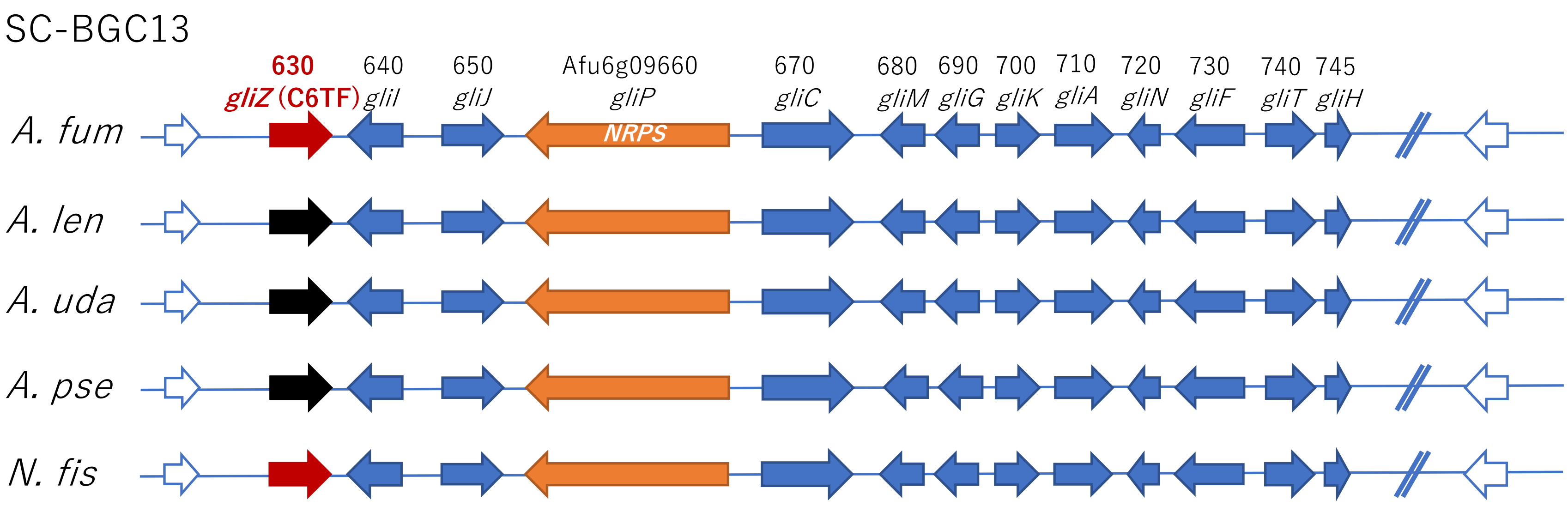

SC-BGC13
630
gliZ (C6TF)
640
gliI
650
gliJ
Afu6g09660
gliP
NRPS
670
gliC
680
gliM
690
gliG
700
gliK
710
gliA
720
gliN
730
gliF
740
gliT
745
gliH
A. fum
A. len
A. uda
A. pse
N. fis

**N**

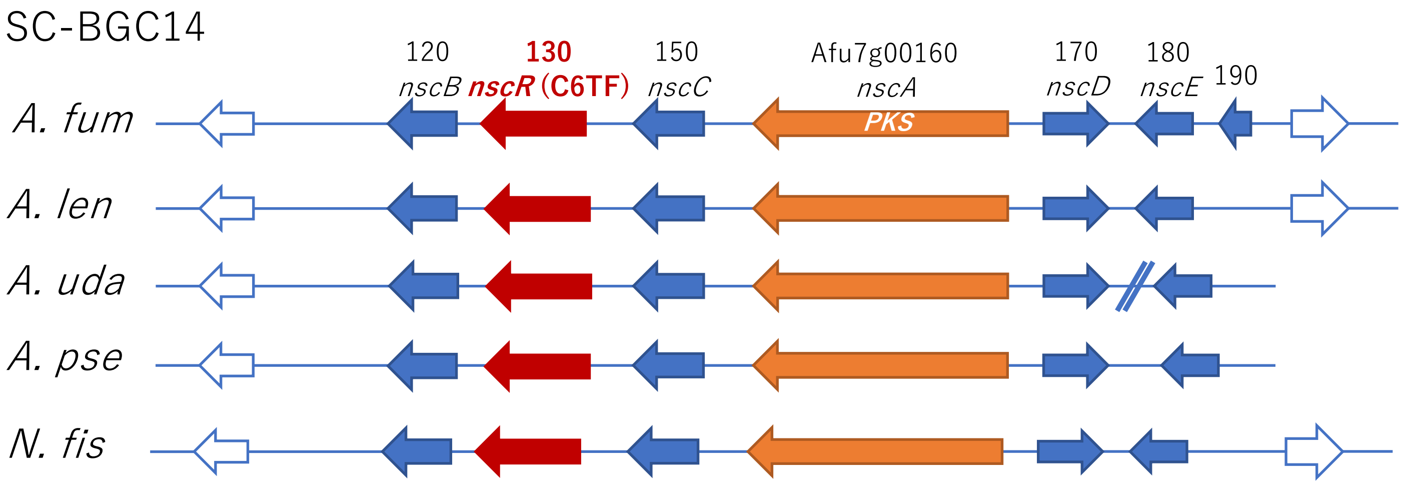
SC-BGC14
120
nscB
130
nscR (C6TF)
150
nscC
Afu7g00160
nscA
PKS
170
nscD
180
nscE
190
A. fum
A. len
A. uda
A. pse
N. fis

**O**

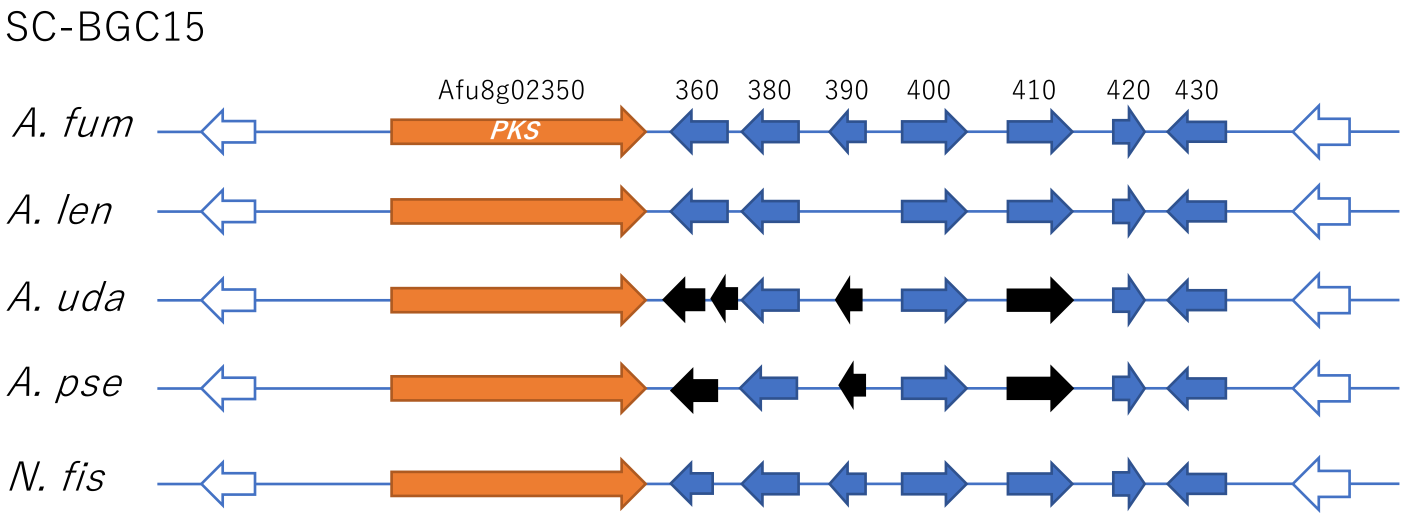
SC-BGC15
Afu8g02350
360
380
390
400
410
420
430
A. fum
PKS
A. len
A. uda
A. pse
N. fis
