## Supplementary figures and images for "Comparative genome and transcriptome analyses revealing interspecies variations in the expression of fungal biosynthetic gene clusters"

### Fig S2

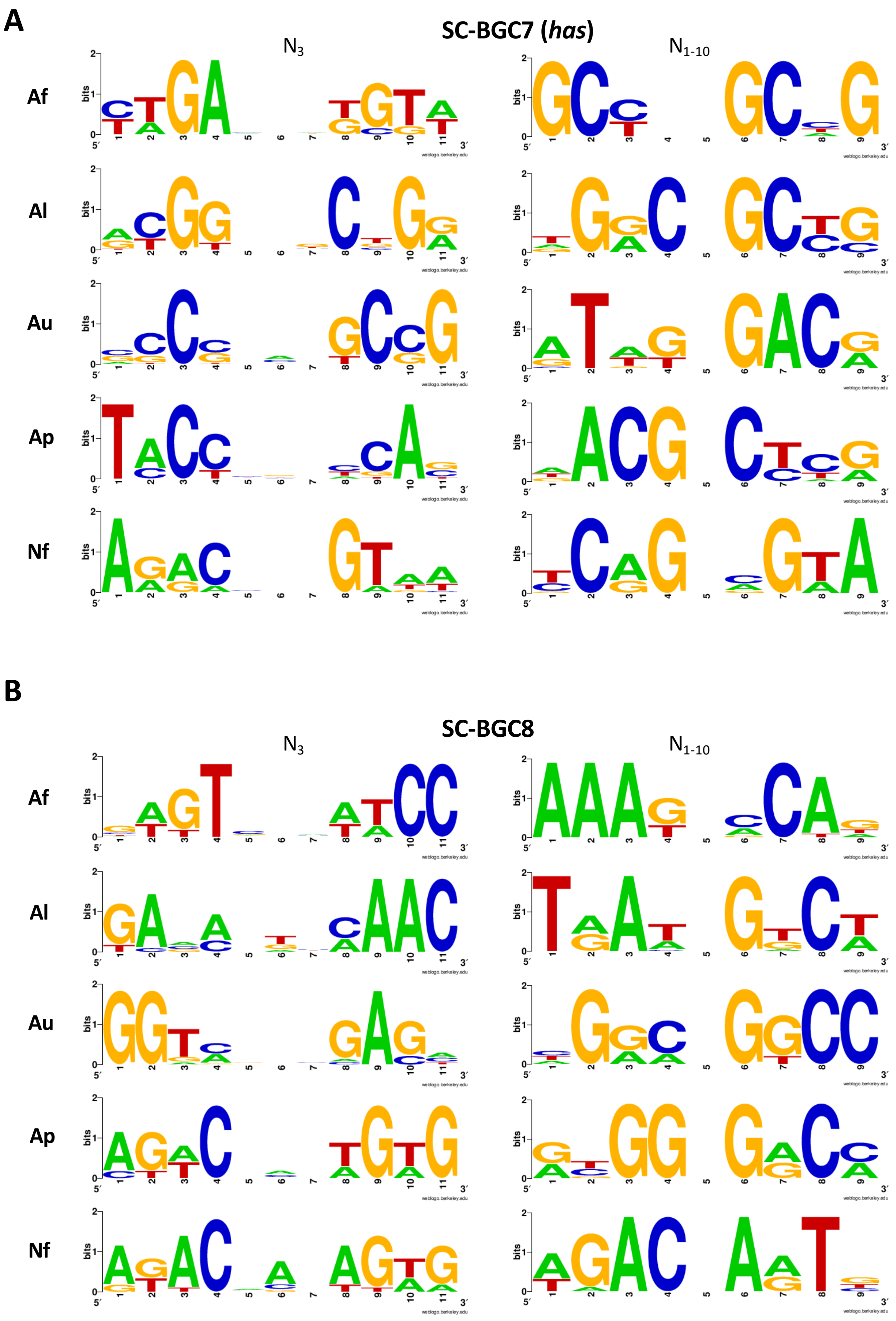

**C**

**SC-BGC9 (*tpc*)**

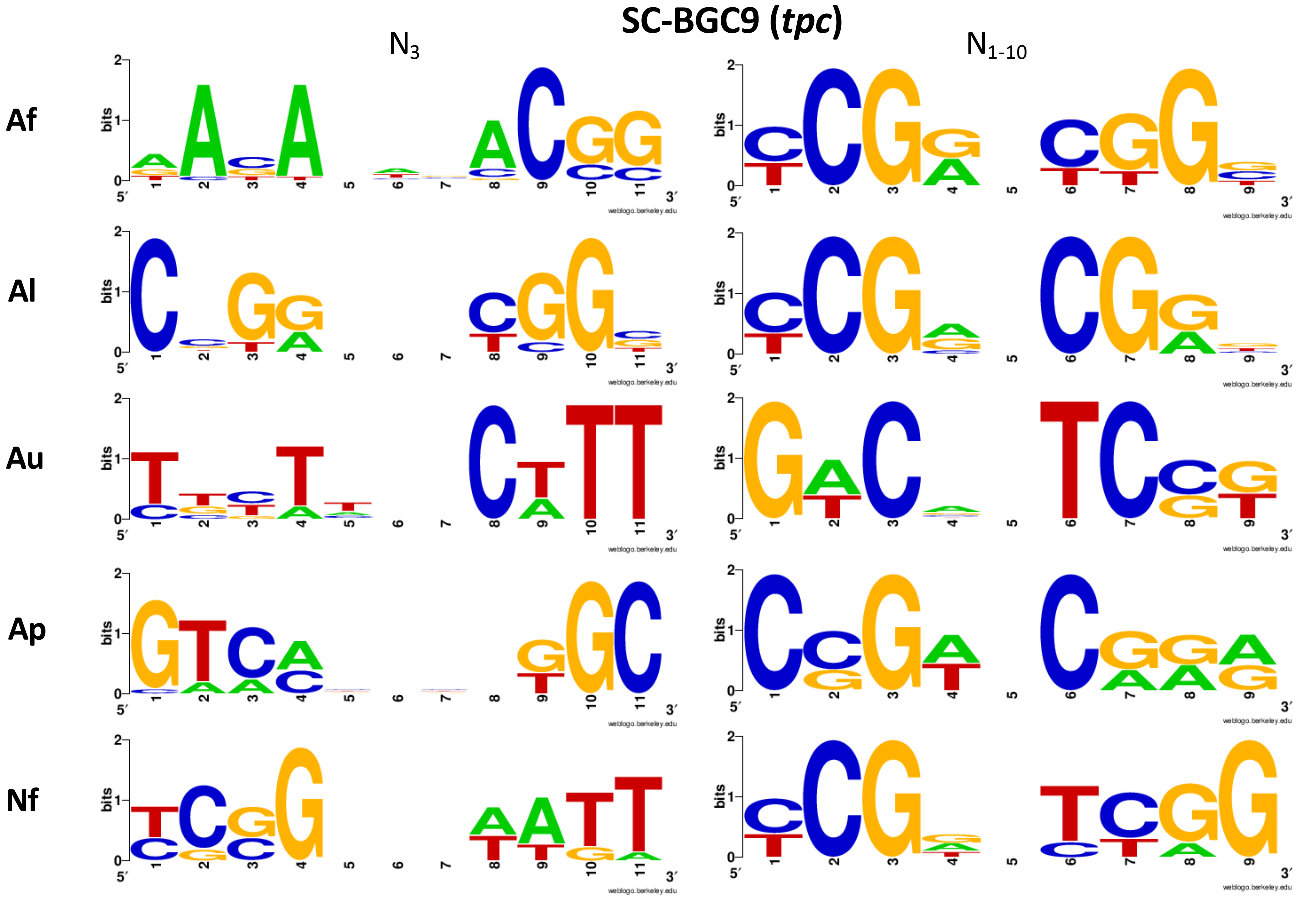


**D**

**SC-BGC10**

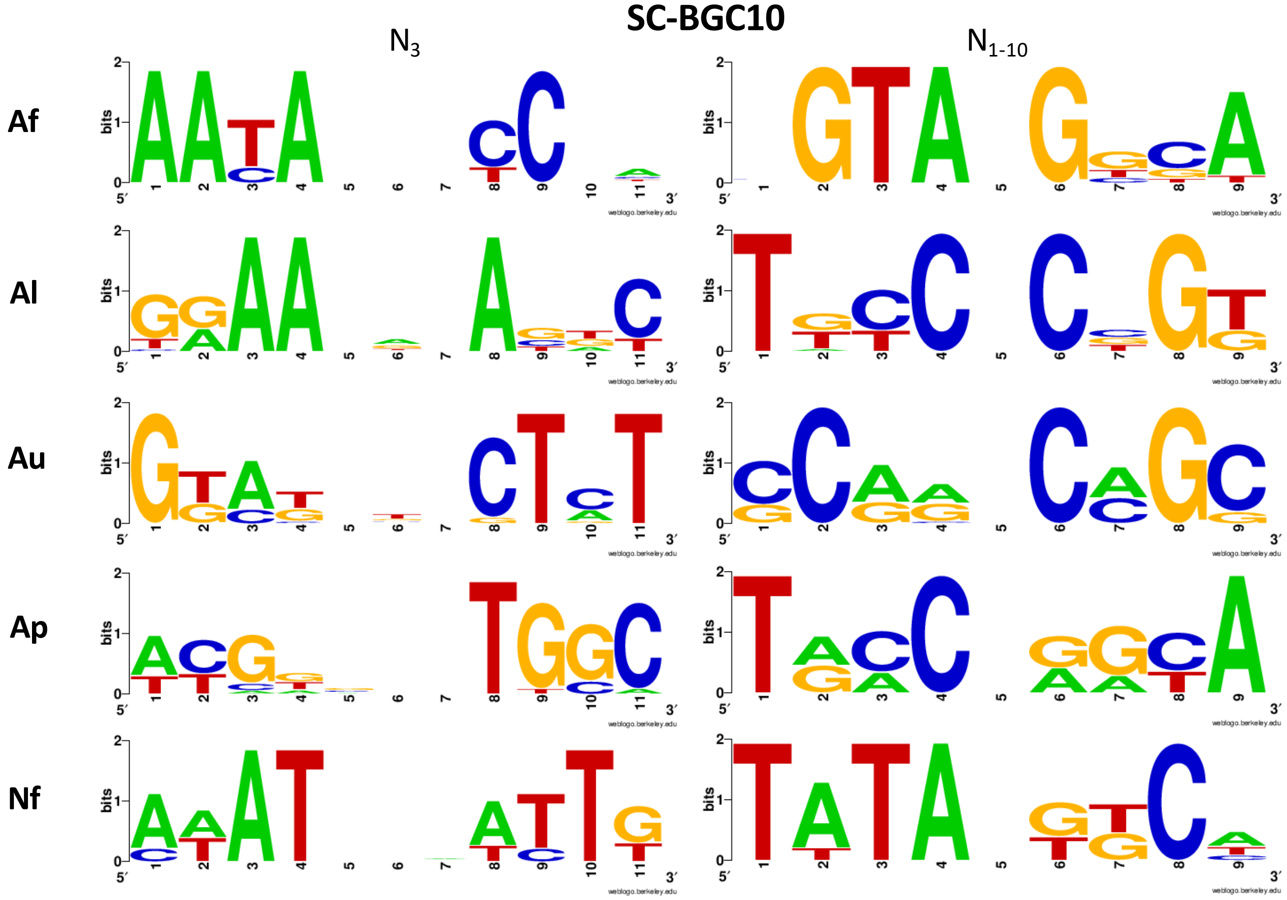

**E**

**SC-BGC11 (*fsq*)**

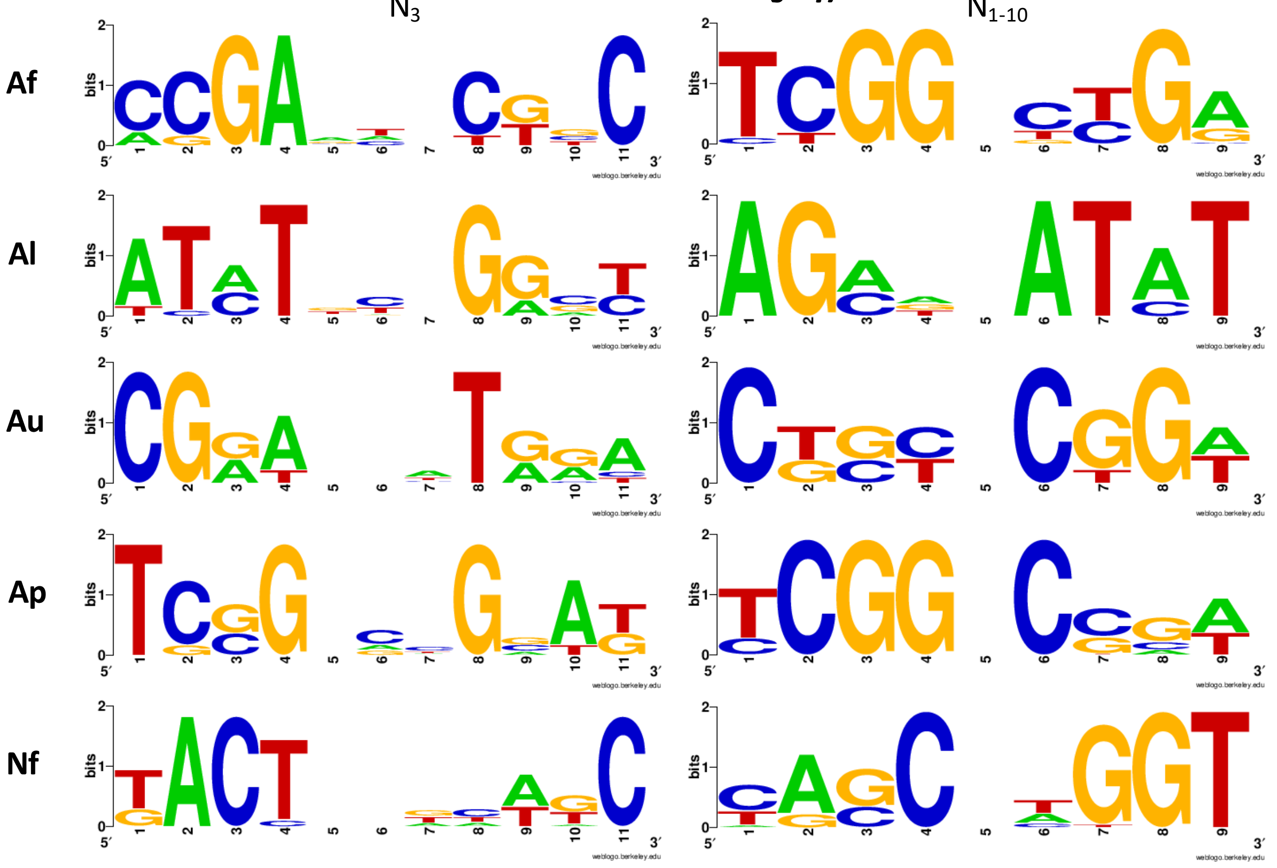


**F**

**SC-BGC12**

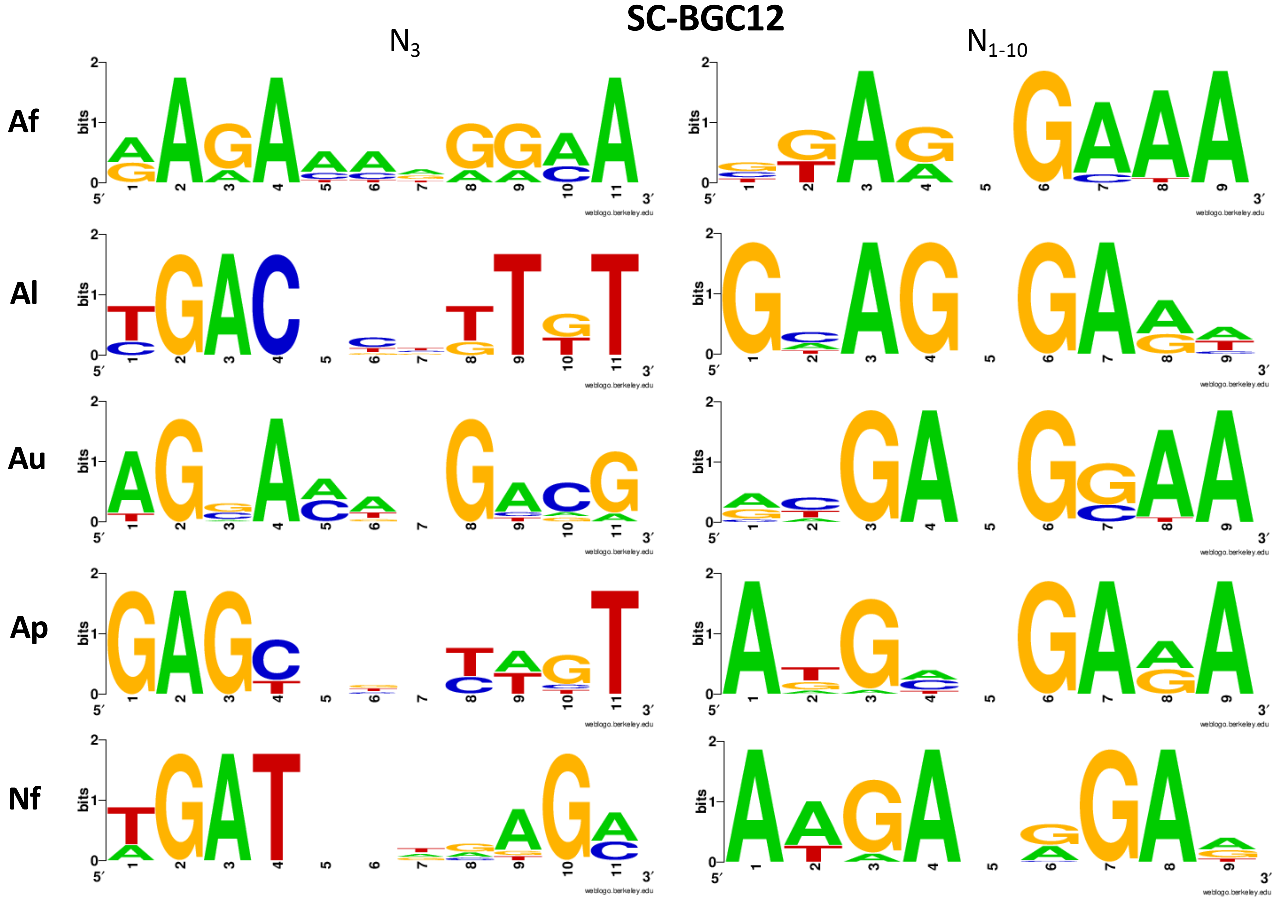

**G**

**SC-BGC13 (*gli*)**

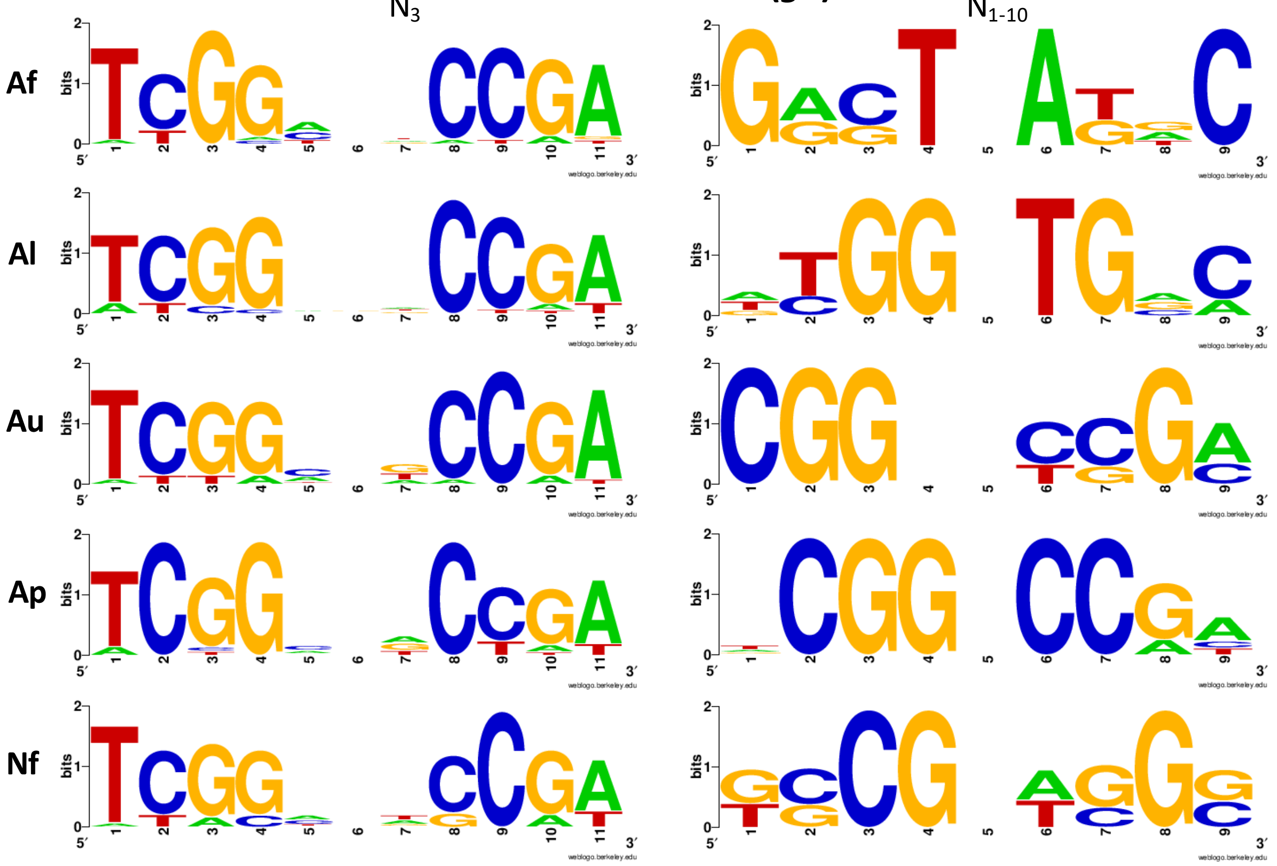


**H**

**SC-BGC14 (*nsc*)**

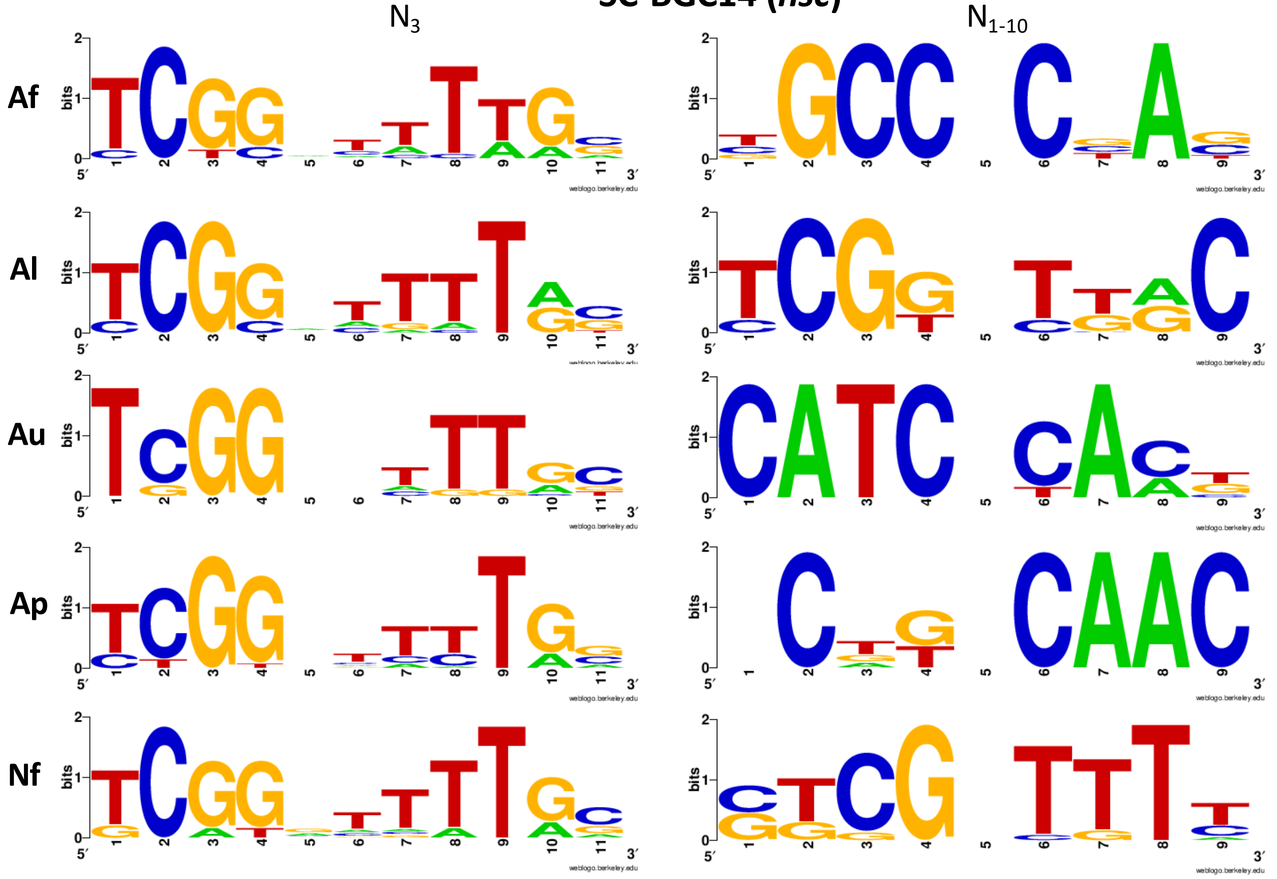


Fig. S2

### Fig S4

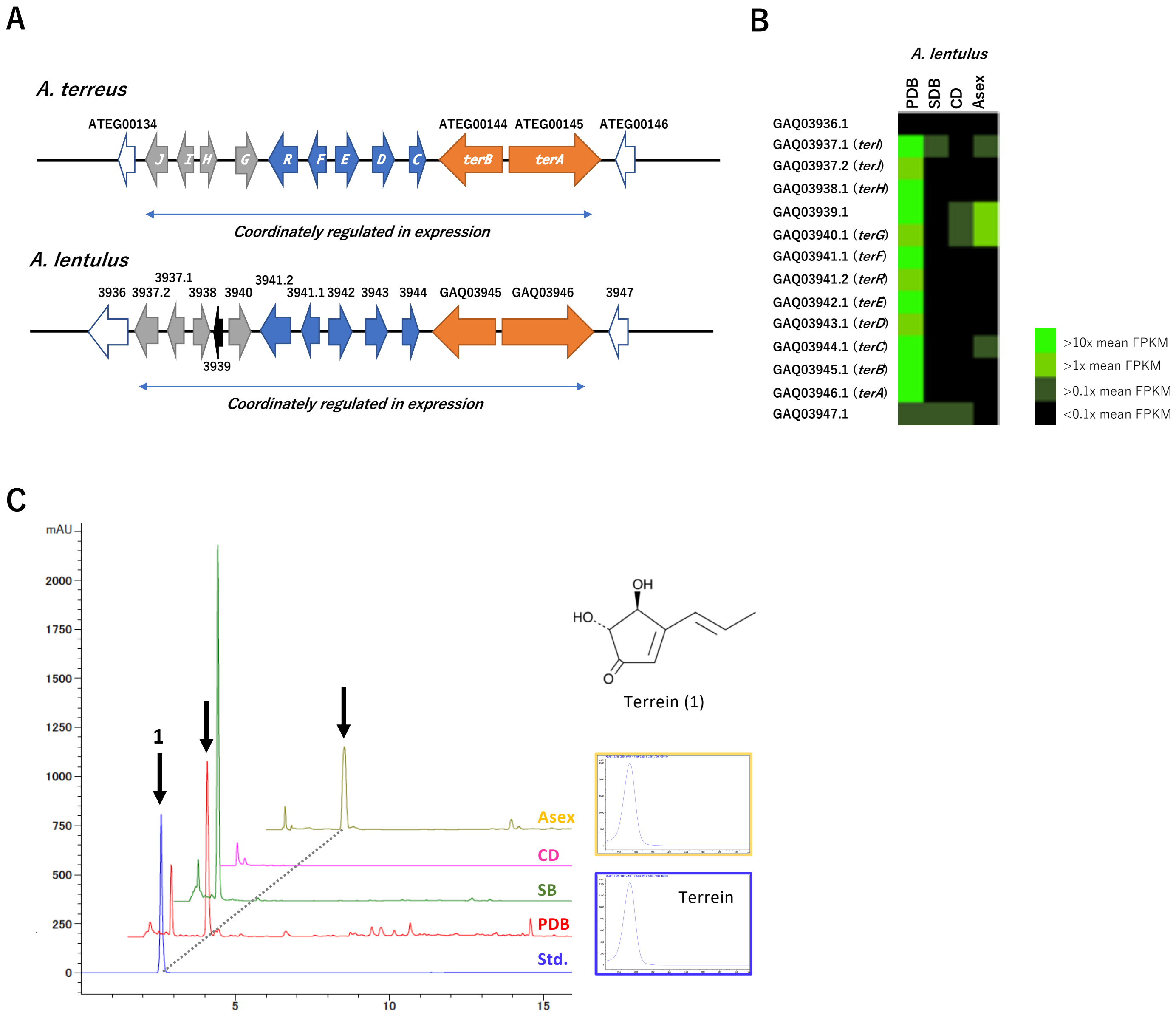
