## Supplementary material for "Comparative genome and transcriptome analyses revealing interspecies variations in the expression of fungal biosynthetic gene clusters": Fig S3

### Alignment of *gliZ* upstream regions (-500)

**A**

```
Afu6g09630 (Af)     ---------TC------ATAGTGAACAACATATCAGCCGCGACTTCAAACATGCATCGAG 45
NFIA_055320 (Nf)    GCCGCAGATTCTGGGATAGAGTGAAGAACATGTCAACATCAACTT--GACATGAACCGAG 58
gene_01174.t1 (Ap)  ----------C---------GT--------TGTCAACATCAACTT--GGCATGACTCTAG 31
gene_00590.t1 (Al)  ------------------------------------------------------------
gene_09462.t1 (Au)  ------------------------------------------------------------

                    AGTCAATTGAGTTTGATTGCGGGATCAGGTGGAGGGACATTGGGC-------------CC 92
                    AGTCAATTGAGTTTGATTGCGGGATCAAGTGGTGCGACATTCGGC-------------CA 105
                    --TCAATTGAGTTTGATTGCGGGACTAGGTGCCGAGACATTCATCGTCCTTTCTTTTCCG 89
                    ----------GTTTGATTGCGGAT-CAAGTGGTCAGACATTTG-------------GCCA 36
                    ----------ATCTAGTGCCGAGA-CATCT--CTCGCCCTTTGTT----------TGCCG 37
                              * *  *  **     *  *     * * **               *

                    TCTGACAGGCGGATCCGCATCGTCAG--GCGGTCCAAAACCGTAAGCTGCATGCGCGAAG 150
                    TCTGACAGGCGGATCCGCATCGTCAG--GCGCTCCAAAACCGTAAGCTGCATGCGCGAAG 163
                    TCTGACAGGCTGATCCACATCGTCAGAGGCGCTCCAAAATTGTAAGCTGCATGCGCGAAG 149
                    TCTGACAGGCGGATCCGCATCATCAG--GCGCTCCAAAACTGTAATCTGCATGCGCGAAG 94
                    TCTGACAAGCGGATCCGCCTCGTCAGAGGCGCTCCAAATCCGTAAGCTGCAGGCGCGAAG 97
                    ******* ** ***** * ** ****  *** ******   **** ***** ********

                    GATTCTGAGGCGCAAA--ACCCCCCTGCTAGTGGTGCTGATTCAGTCAGGACCGACGGAT 208
                    GATTCTGAGGCGCAAA--ACCCCCCTGCTAGTGGTGCTCATT----CAAGACAGACTGAT 217
                    GATTCTGAGGCGCAAGG-AAACCACTGCTAGTGGTTCAGATT----CAGGACGGACCGAT 204
                    GATTTTGAGGCGCAAATGAAG-CCCTGCTAGTGGTGCTCGTT----CAGGACGGACTGAT 149
                    GATTCTGAGGCGCAAGGCAACTCCCTGCTAGTGGTGCTGATT----CAGGACGGACTGAT 153
                    **** **********   *   * *********** *   **    ** *** *** ***

                    CGAGATTGTCAAATGTCAACTGAATCGAAGTCAATCTAGTTGCAA--------ACGTTGT 260
                    TGACATTGTCAA-TGTCAACTGAACCGAAGTCAATTTAGTTGCAATGCGGATCAAGTTGC 276
                    TCACATTGTCA--CGTCGACTGAATCGAGGTCAATTTAGTTGGAGGGCGGATCAAGTCGT 262
                    TGACATTGTCAA-CGTCAACTAGATCGAGGTCAATTTAGTTGGACTGCGGATCAAGTTGA 208
                    TGACATTGTGAA-CGTCGACTGAGTCGAGGTCAATTTAGTTGGAGTGCGGATTAAGTCTC 212
                      * ***** *   *** ***    *** ****** ****** *         * **

                    AACAGACAGTCTCCTCGCTTTCTGACAGGCCAAATCCGCGCAATCAGAGGAGCTGCAAAG 320
                    GACAGACACTCTCCTCGCTTTCTGACAGGCCAAATCAGCGTAATCAAAGGCGCTGCAAAG 336
                    GACAGAGACTCTCCTTGTTCTCTGACAGGCCAGAT----------------GCTGCAAAG 306
                    GACAGACGCTTTCCTTGCTCTCTGACAGGCCAGATGAGCGTCATCAGAGGTGCTGCAAAT 268
                    GACAGAAACTCTCTTGGCTCTCTGACAGGCCAGATCAGCGTCGTCAGAGGTACTGCAGGG 272
                     *****   * ** * * * ************ **                 *****

                    GGTACTCGGCATGCAGGGGGGC-ATTCAGCTGTGCAGAATA------TCCGGCCCCCTTG 373
                    TGTACTCGGCATGCATGGAGAC-ATTCAGGTGTCAAGAATA------CCC----CCCTTG 385
                    CGTAATCGGCATGCACGGAGACCATTCAGGTGT-CAGAATA------CCC-----CCTTG 354
                    -GTACTCGGCATGCATGGAGAG-ATTCAGGTGT-CAGAATAGAATCCCCC-----CCTTG 320
                    -TTAATTGGCATGCACGGAGAC-ATTCAGGTGT-CAGAATA------CCC-----CCTTT 318
                      ** * ******** ** *   ****** ***  ******       **     ****

                    CC------------ACCCGTGTTGCACTG---CGAT----------AGAAGGTTAAACGC 408
                    C-------------ACCAGCA-------G---CGAT----------AGGAGGTTAAAGGC 412
                    CCAGGGTGCTGCAAAGTGATGGTGCAAAG---TGATGGGAGGTTAAAGGAGGTTAAACAC 411
                    CC------------ACCGGTGCTGCAAAA--GCGAT----------AGGAGGTTGAACGC 356
                    CC------------AGTGGTGGTGCAGAGCGATGAT-----------GGAAGGTTAAAGC 356
                    *             *                ***           * ***** **  *

                    GAAC-CTTCTCCGAGGGCGATCGCTGCTAATATTTAAGGGCCGGTAGTCTACCTCTTCCA 467
                    G-----TTCTCCGAGGGCGATTACTTCTAGTATTTAAGGGCCAGTAGTCTACCTCCTCCA 467
                    GTTCTCTCCTCCGAGG-CG-TTGCTGCTAATATTTAAGG-TCGCTAGTCTACCTCCTCCA 468
                    GTTCTC-----CGAGGCCGTTCGGTGCTAATATTTAAGG-TCAGTAGTCTACCACCTCCA 410
                    CTTCTCTTCTCCGAGGC-GTT-GCTGCTAATATTTAAGG-CCAGTAGTCTACCACCTCCA 413
                              *****  * *   * *** *********  *  ********* * ****

                    TGGGATCGCCAAAGAGCAAA-CTACTCGTCAGCG-------------------------- 500
                    TGGGATCGCCAAAGAGCAAA-CTACTCGTCAGCG-------------------------- 500
                    TGGGCCTGACCAA-AGCTAA-CTACTCGTCAGCG-------------------------- 500
                    TGGGGTTGCCAAA-AGCCAAACTACTCGTCAGCGATGGCGACAGCTATGCAGGATGTGTC 469
                    CGGGTCAATCA---AGCCAA-CTACTCGCTAACGATGGCGGCATCTATGCAGGATGTGTC 469
                     ***     *    *** ** *******  * **

                    ------------------------------
                    ------------------------------
                    ------------------------------
                    GCCATCTTTCACCTCGTCTCCGACGTCCACA 500
                    GCCATCTTTTACCTCGTCTCCCACATCCACA 500 (-1 from initiation site)
```

**B**

### Alignment of *gliI* upstream regions (-500)

```
gene_01175.t1 (Ap) ---------------ATTGGCATATCATACAGATGGGCATTCTGGCCATTGATGTCATTG 45
gene_09463.t1 (Au) AGATCAGTCTGGCCGATTGGCATATCATACAGGTGGGCATTCTGGCCATTGATGTCATTG 60
gene_00589.t1 (Al) -------TCTGGCCGATTGGCATATCATACAGATGGTCATTCTGGCCATTGATGTCATTG 53
Afu6g09640 (Af)    ------GTCTGCCCGATGGGCATATCATACAGATGGGCATCCTGGCCATTGATGTCATTG 54
NFIA_055330 (Nf)   ---TCAGTCTGGCCGATTGGCATATCATACAGATGGGCATCCTGGCCATTGATGTCATTG 57
                                      ** ************** *** *** *******************

                   CGAAACCATCCGCGTATCATGTACGGAAAGTCATTGTGGCCGTCGATCAGAGGCACCTCT 105
                   CGAAACCATCCGCGTATCATGTACGGAAAGTCATTGTGGCCGTCGATCAGAGGCACCTCG 120
                   CGAAACCATCCGCGGATCATGTATGGAAAGTCATTATGGCCATCGATCAGGGGCACTTCT 113
                   CGGAACCATCCGCGTATCATGTATGGAAAGTCATTATGGCCGTCGATCAGAGGTACCTCT 114
                   CGAAACCATCCGCGTATCATGTATGGAAAGTCATTATGGCCATCGATAAGAGGTACCTCT 117
                   ** *********** ******** *********** ***** ***** ** ** ** **

                                                                gliJ
                   TTCAGAAGCCTCTGGGCCTGTTCTTCGATCGCGCTCGGAGTCATTCTAGGGCTGGTAAAA 165
                   TTCAGAAGCCTCCGGGCCTGTTCTTCGATCGTGCTCGGAGCCATTCTAG----------- 169
                   TTCAGAAGCCTGCGGGTTTGTTCTTCGATCGCGCTCGAAGTCATTCTAG----------- 162
                   TTCAGAAGCCTCGTTGCCTGCTCTTCGATGGGGCTCGGAGACATTCTTG----------- 163
                   TTCAGAAGTCTCCGTGCTTGCTCTTCGATCGCGCTCGGAGACATTCTAG----------- 166
                   ******** **    *  ** ******** * ***** ** ****** *

                   TCTCTAGCTGAGTTTGGTATAATTTCTAGCTGAATTTGGTATCCTGTCTAGCATGGTGAT 225
                   -------CTATGTGTGGTATAATTCCTAGCTAAGTTTG-TATCTTGTCTAGCATGATGGT 221
                   -------TTAAGTCTGGTATAATTCCTAGCTAAGTTTGGTATTCTGTTTATCATGGAGAT 215
                   --------CATGTCAAGGATTGTTCCTAGCTCAATTTG-TATTCTGTCTTTCACGAAAAC 214
                   --------CATGTCCAGTATTGTTCCTAGCTAAGTTTG-TATTCTGTCTATCATGGAGAT 217
                              **   * **  ** ****** * **** ***  *** *  ** *

                   CTG-TCTTCCATTT-CTT--CGTTGTGTCTCGCGGAGGAC-AATGTACCCTACTTCAGCC 280
                   CTGATATTCCATTT-CTTGTCGTTGGAACTGGCGGAGGAC-AATGTACCCTACTTCAGCC 279
                   TGGATATTCCGTTT-CTT--CGTTGTGTTTTGTGGAGGTCTGATGTACCCTACTTCAGCC 272
                   CTGGTATTCCATTTTCTTGTCGTTGGGTCTG-TGGAG-TC-AATGTACCTTTCTCAAGCC 271
                   CTGATATTCCATTT-CTTGTCGTTGGGTCTGGCGGAG-TC-AATGTACCTCTCTCAAGCC 274
                     * * **** *** ***  *****     *    ****   *   *******   **   ****

                          #1
                   TCTCAATTCGGCCACCGAAATCCAAGTTCCATCCACTCATTCGAATCTCTATGCAAAACA 340
                   TCTTAATTCGGCCACCGAAATCAAAGTTCCACCCACTCATTCGAATTTCTATGCAACACA 339
                   TCTTACTTCGGTGGCCGAAATCCAA-CTTCATCCACTCATTTGCATTTCCATGCAGCACA 331
                   TCGTACTTCGGTGACCGAAA-CCAA-CTGCATCCGC-CCTGTGCATTTCTATGCAACACA 328
                   TCTTACTTCGGTGACCGAAA-CCAA-CTTCATCAAC-CATATGCATTTCTATGCAACACA 331
                   **  * *****    ****** * **  * ** *   * * *  * ** ** *****  ***

                                                  #2
                   TGTTATTGTTGA-TTGGTTTTCGTCAATCTCGGATTCCGAT-ATTCT-----TTTACAAA 393
                   TGTTACTGTTGA-TTCTAATTCGTCGATCTCGGAATCCGAT-ATTCT-----TTTACGCA 392
                   TGTTATCGTTGA-TGTAAATTCGTTCATTTCGGATTCCGAT-TCCTTCCTCATTTACACA 389
                   -GTCGATGGTGAGTTTTGATTCGTCAATTTCGGCCTCCGATTATTAT-----TTTAGACG 382
                   -GTTATTGCTGAGTTTTAATTCGTCGATTTCGGACTCCGATTAAGAT-----TTTACGCG 385
                    **     * *** *      *****  ** ****  ******     *      ****

                   GTAGCTTCAATAGATAGAAGCGTGCT--ATCTTTTCAA-GCAGTTGATCATGACATGCTG 450
                   GACTTTTCAATAGATGGAAACCTGCT--ATCTTCTCGACACTGATGATCATGATACGAGG 450
                   GTTTCCTCAATGGATAGAAACGTGCC--ATCTTTTCGAGACTGCTGATCATGATATGGTG 447
                   GTCCCTTGAACGGATAGAAACGTGCGCGATTCTTT-GAGGCCGTCGATCTTGATGTGACA 441
                   GTCTCTTCAACGGATAGAAACGTGCGCGATTCTTTTGAGGCAGTCGATCTTGATACGACG 445
                   *     * **   *** *** * ***    **   * *   *   * *   **** ***    *

                   A--AGTCCATCTAACAAGTCAGATCGGGTATAACACCAA-CA--------GCGACAACAC 499
                   A--GGCCCATCTAACAAGTCAGGTCAGGTAAAACGCAAA-CA--------GCGACAACAC 499
                   ACAAGTCAATCTAACAAGTCAGGTCAAGTTTAACGCAAAACA--------GAGATAACAA 499
                   A--AGTCCATCTACCAGGTCCGGTCACGTAGAAGGCAAAGCAACCAAGACGAGACAACAC 499
                   A--AGTCCATCTACCAAGTCAGGTCACGTATAACGCA----AACGGAGGCGAGACAACAC 499
                   *   * * ***** ** *** * ** *   ** *   **   *         * ** ****

                   A 500
                   A 500
                   A 500
                   T 500
                   T 500 (-1 from initiation site)
```

**c**

### Alignment of *gliJ* upstream regions (-500)

```
gene_01176.t1 (Ap) -----------CGGACGGACCAGTTCTCCGCCGTCATCATATTGACTCGGTTCTGTTTCG 49
gene_09464.t1 (Au) --AGCTCGCCGCGGACGGACCAGTTCTCCGCCGTCATCATATTGACTCGGTGCTGTTTCG 58
gene_00588.t1 (Al) CGAGCTCGTCTCGGATGGACCAGTTCTCCGCCGTCATCATGTTGACTCGGTGCTGTTTGG 60
Afu6g09650 (Af)    --AGCTCGTCTCGGATGGACCAGTTCTCCGCCGTCATCATGTTGACTCGATGCTGTTTGG 58
NFIA_055340 (Nf)   -----------CGGATGGACCAGTTCTCCGCCGTCATCATGTTGACTCGATGCTGTTTGG 49
                              **** *********************** ******** * ****** *

                   CATTGAGCCGAGACACCTGTCGCATATGCTTGTTGATGAACCATTGGATGTTTGCGCGAG 109
                   CATTGAGCCGAGATACCTGCCGCATATGCTTGTTGGTGAACCATTGGATGTTTGCGCGAG 118
                   CATTGAGCCGAGCGACCTGTCGCATATGCTTGTTGGCAAACCATTGAATGTTTGTGCGAG 120
                   CATTCAGCCGAGACACCTGCCGCATGTGCTTGTGGGTGAACCACTGGATGTTTGTGCGAG 118
                   CATTGAGCCGAGACACCTGTCGCATATGCTTGTTGGTGAACCACTGGATGTTTGTGCGAG 109
                   **** *******  ***** ***** ******* *   ***** ** ******* *****

                                                                         gliI
                   AACGATGTGCCAGGGAGGCAGTAATTGTCTCT------------GATGGCATTG 151
                   AACGATGCGCTAGAGAGGGAGTAATTATCTCAGATG------CAGATGGCATTG 166
                   CACGATGCGCTAG---GGGAGGAAGCGTCTCA------------GATGGCATTT 159
                   CACGATGCGCTAGCAAAGGAGTGATCGTCTCG------------GATGGCATAG 160
                   CACGATGCGCTAGGGAAGGAGTGGTTGTTTCAGATGGCATTTCAGATGGCATAG 163
                    ****** ** **     * **       * **           ********

                   TGTTGTCGC--------TGTTG-GTGTTATACCCGATCTGACTTGTTAGATGGACTT--C 200
                   TGTTGTCGC--------TGTTT-GCGTTTTACCTGACCTGACTTGTTAGATGGGCCT--C 215
                   TGTTATCTC--------TGTTTTGCGTTAAACTTGACCTGACTTGTTAGATTGACTTGTC 211
                   TGTTGTCTCGTCTTGGTTGCTTTGCCTTCTACGTGACCGGACCTGGTAGATGGACTT--T 218
                   TGTTGTCTCGCCTCCGTT----TGCGTTATACGTGACCTGACTTGGTAGATGGACTT--C 217
                   **** ** *        *      *  **   **   ** * *** ** ***** * * *

                   AGCATGTCATGATCAACTGC-TTGAAAAGATAGC--ACGCTTCTATCTATTGAAGCTACT 257
                   CTCGTATCATGATCATCAGTGTCGAGAAGATAGC--AGGTTTCCATCTATTGAAAAGTCT 273
                   ACCATATCATGATCAGCAGTCTCGAAAAGATGGC--ACGTTTCTATCCATTGAGGAAACT 269
                   GTCACATCAAGATCGACGGCCTCAAA-GAATCGCGCACGTTTCTATCCGTTCAAGGGACC 277
                   GTCGTATCAAGATCGACTGCCTCAAAAGAATCGCGCACGTTTCTATCCGTTGAAGAGACC 277
                     *   *** ****  * *  *  *    ** **  * * *** ***  ** *     *

                                       #2
                   TTGTAAAAGAATA-----TCGGAATCCGAGATTGACGAAAACCAA-TCAACAATAACATG 311
                   GCGTAAAAGAATA-----TCGGATTCCGAGATCGACGAATTAGAA-TCAACAGTAACATG 327
                   GTGTAAATGAGGAAGGAATCGGAATCCGAAATGAACGAATTTACA-TCAACGATAACATG 328
                   GTCTAAAATAATAA----TCGGAGGCCGAAATTGACGAATCAAAACTCACCATCGAC-TG 332
                   GCGTAAAATCTTAA----TCGGAGTCCGAAATCGACGAATTAAAACTCAGCAATAAC-TG 332
                      ****     *     *****  **** **  *****      * *** *    ** **

                                                                 #1
                   TTTTGCATAGAGATTCGAATGAGTGGATGGAACTTGGATTTCGGTGGCCGAATTGAGAGG 371
                   TGTTGCATAGAAATTCGAATGAGTGGGTGGAACTTTGATTTCGGTGGCCGAATTAAGAGG 387
                   TGCTGCATGGAAATGCAAATGAGTGGATG-AAGTTGGATTTCGGCCACCGAAGTAAGAGG 387
                   TGTTGCATAGAAATGCACAGG-GCGGATG-CAGTTGG-TTTCGGTCACCGAAGTACGAGG 389
                   TGTTGCATAGAAATGCATATG-GTTGATG-AAGTTGG-TTTCGGTCACCGAAGTAAGAGG 389
                   *  ***** ** ** *  * * *  * **  * ** * ******    ***** *  ****

                   CTGAAGTAGGGTACATT-GTCCTCCGCGAGACACAACGA--AGAAA-TGGAAGA-CAGAT 426
                   CTGAAGTAGGGTACATT-GTCCTCCGCCAGTTCCAACGACAAGAAA-TGGAATATCAGAC 445
                   CTGAAGTAGGGTACATCAGACCTCCACAAAACACAACGA--AGAAA-CGGAATATCCAAT 444
                   CTTGAGAAAGGTACATT-GAC-TCCAC-AGACCCAACGACAAGAAAATGGAATACCAGGT 446
                   CTTGAGAGAGGTACATT-GAC-TCCGCCAGACCCAACGACAAGAAA-TGGAATATCAGAT 446
                   **  **   *******  * * *** * *    ******  *****  **** * *

                   CACCATGCTAGACAGGATACCAAATTCAGCTAGAAATTATACCAAACTCAGCTAGAGATT 486
                   CATCATGCTAGACAAGATAC-AAACTTAGCTAGGAATTATACCACACATAGCTAGA---- 500
                   CTCCATGATAAACAGAATACCAAACTTAGCTAGGAATTATACCAGACTTAACTAGA---- 500
                   TTTCGTGAAAGACAGAATAC-AAATTGAGCTAGGAACAATCCTTGACAT-GCAAGA---- 500
                   CTCCATGATAGACAGAATAC-AAACTTAGCTAGGAACAATACTGGACAT-GCTAGA---- 500
                      * **  * ***  **** *** * ****** **  ** *   **     * ***

                   TTACCAGCCCTAGA 500 (-1 from initiation site)
                   --------------
                   --------------
                   --------------
                   --------------
```

**D** 

#### Alignment of *gliP* upstream regions (-500)

```
Afu6g09660 (Af)     GTCTATGACGACGGAGAGAACCCAGTAGAGAACGGGCCGGGCCGCCACGAGGGCCAGCAC 60
NFIA_055350 (Nf)    ----------ACAGAGAGAACCCAGTAGAGAACAGGCCGGGCCGCAACGAGGGCCAGCAC 50
gene_00587.t1 (Al)  ------------AGACAGAATCCAGTAGAGAACAGGCCGGGCCACAACGAGTGCCAGCAC 48
gene_09465.t1 (Au)  --------------------GCCAGTAGAGAGCAGGTCGGGCCACCACGAGAGCCAGTAC 40
gene_01177.t1 (Ap)  ----------------------------------GTCGGGCCACCACGAGGGCTAGGAC 25
                                                      * ****** * ***** ** ** **

                                                            gliC
                    CATGCAGGGAACGAGTATAGTCAGTGTGAAAGCCATGGCAAGACGGTTTGAAAGTGCCAA 120
                    CAAACATGGGACGAGTATAGTCAGTGCGTAAGCCATGGCGAGGGAGTTAGAAAGTGCCAA 110
                    CAAACAGGGGACGAGTACAGTCAGTGTGTAAGCCATGGTGAGGGAGTTGGAAAGAGCCAA 108
                    CAGACAGGGGAGGAGTATAGTGAGTGTGTAACCCATGATGAATGAGTTGGGAAGGGTCAG 100
                    CAGACAGGGGAGGAGTATAGTCAGCGCATAAGCCATGGCGAGGGAGTTCAAAAGAGTCAA 85
                    **  ** ** * ***** *** ** *   ** *****   *    ***   *** * **

                    A------------CGACGGGTCGAAACCGTAGTGAATAGGGGCTTGTCCATACCTTAACA 168
                    C------------AGTATGGTCGAAACAGCAGTGAATAGGGGCTTGTCGATACCTTAACA 158
                    C------------AGAATGGTCGAAACTGTAGTGAATAGGGGCTTGTCGATACCTTAACA 156
                    C------------AGAATGGTCTGTACAGTAGTGAACAGGGGCTTGCCG-TACCTTAACA 147
                    CGGAAGGGTAAACAGAATGATCGGTACAGTAGTGAATAGGGGCTTGTTGATACCTTAACT 145
                                 *   * **   ** * ****** *********    *********

                                                                          #1
                    GC-AGCGACCGCCGCTGCAAGGCCTACC-----GCTGAAGCCC-TGCTCGGAGGCCGACT 221
                    GC-AGCGACCGCCGCTGCAAGGCCTACC-----GCTGAAGCCC-TGCTCGGAGGCCGACT 211
                    GC-AGCGACCGCCGCTGCATGGCCTACC-----GCTGAAACCC-TGCTCGGAGGCCGACT 209
                    GGGAACGACCGCCGCTGCAGGGCCTATT-----TCTGGAGCCCCTGCTCGGACGCCGACT 202
                    GCAAGAGACCCCGGCTGCAGGGCTTGCCTAGGGGCTGCAGCCCGTGCTCGGAGGCCGACT 205
                    *  *  **** * ****** *** *         *** * *** ******** ********

                      #2
                    TTTCGGAGGCCAAACTCATCC-GGAAAACGGGATAGTGCCTATACCAGGAGTGGCAATAG 280
                    TTTCGGAAGCCAAACCAACCC-GGAAATCGGGATAGTGCCTTTACCAGGAGCGGCAATAG 270
                    TTTCGGAGGCCAAACGAAGCC-GGAAATCGGGATGGTGCCTATACTAGGGGTGCCAATAG 268
                    TTTCGGAGGCCAAACTACCCCTGGTAATCGGGATAGTCCCTAAACTAGGGGTGGCAATAC 262
                    TTTCGGCGGCCAAACCACCCCGGGCAATCGGGACAATGCCTATACTGGGAGTA-CAACAC 264
                    ******  ********    ** ** ** *****   * ***   **   ** *   *** *

                    TGGATGCCAATAGGAACAGCATGTGGAGTCCAATTGAA-TAC-CTCCCACTGGACTGCAT 338
                    TGGATGCCAATAGGAACTGTACGTGGACTCCAAAGAAA-TAC-CCCCCACTGGACTGCAT 328
                    TGGGTGCCAATAGGGACTGTACGTGGAGTCCAAAGGAA-TACGCCCCCGCCATAGTGCAG 327
                    AGGGTACCGATAGGGATTGTACGTGGAGTCCAAAGGAAATATACCCCCGCCATGGTGCAT 322
                    AGGATACCAGTAGGGACTGTAGGTATAGTCCAAAGGAAGTACGCCGCCACTATAGTGCAT 324
                     ** * **  **** *  * * **  * *****   ** **  *  ** *     ****

                    GAG-----------CCATCCCCGGCCATCCCCGAGAGATAGCCATATGAAGCATC---GA 384
                    GAAAGGACGGTTATCCATCCCCGACCATCCCCGAGAGATAGCCATATGAAGCATC---GA 385
                    GAAAGGACGGCCATCCATCCCCGGACATGCCCGACAGATAACCATATGAAGCATC---GA 384
                    GAAGCGGCGGCCATAAGTCTCCGAGC-TCCCCGAGAGATTCGCATATGAAGCATCATCGA 381
                    GAAAGGACGGCCATCCATCCCCGAACCTCTCCGAGAAAAAGGGATATGAAGCATC---GA 381
                    **                 ** ***  * *  **** * *     ************    **

                    CTTACCTCGAAGTCAACCC-C-C--ACCGAAGTTCTCTCAACAGGAAGAAAGTTGGGCAT 440
                    CTTGCCTCGAGGTCAACCT-C-C--AACAAAGTTCTCTCAACAG-AAAAGAGTCGGGCAT 440
                    CTTGCCTCGAGGTCAACCT-CAC--ATCAAACTTCTCTCAACAG-ATAAAAGTCAGGCAT 440
                    CTGCCTCCCAGTTCAGCCTACATTAACCAAACTTCTTTCAACAG-AAAATAGTCAGGCAT 440
                    CTGCCTCCCAGTTCAGCCTACAGCAACCAAACTTCTCTCAACAG-AAAAGAGTCTGGCAT 440
                    **  *  * *  *** **  *     * * ** **** ******* *  * ***  *****

                    GTCTGCTGCTTTTTGAATATCTACTCGATTATTGTCTGGCTCAACCCTACGCTCTGCATC 500 (-1 from initiation site)
                    GTCTGCTCCTTTTTGAATTTGTACTGTATTATTGTCTGGCTCAACCCTACGCTCTTCATC 500
                    GTCTGCTCCTTTTTGAATCTCTACTGTATTATTGTTTGGCTCAACCCTACGCTCTTCATC 500
                    GCCTGCTCCTTTTTAAATATCTACACTATTATTATTTGGCTCAACCCTACGCTCTTCATC 500
                    GTCTACTCCTTTTTGAATCTTTACTATATTATTTTTTGGCTCAACCCTACGCTCTTAAAC 500
                    * ** ** ****** *** * ***    ****** * ******************  * *
```

**E**

### Alignment of *gliC* upstream regions (-500)

```
Afu6g09670 (Af)    CGAGCCGCTCTCATGATCGACTGCCAGCTGGTGTGGTGTCCGAGCGACGGACCGGTCAAA 60
NFIA_055360 (Nf)   ----------TCATGGTCGACTGCCAGCTGGTGGGGCGTCCGAGCGACGGACCGGGCAAA 50
gene_00586.t1 (Al) ------------ATGGTCGACTGCCAGCTGGTGTGGCGTCCGAGCGACGGACCGGGCAAA 48
gene_09466.t1 (Au) --------------------CTGCCAGCTGGTCTGGCGTCCGGGCGACGGACCGGGCAAA 40
gene_01178.t1 (Ap) -----------------------------------GCGTCCGGGCGACGGACCGCGCAAA 25
                                                      * ***** ***********  ****
                                                      gliP
                   AAGCTGGCAGAGGTCGAGCGCTACTACTGATGGCATGATGCAGAGCGTAGGGTTGAGCCA 120
                   AAGCTGGCAGAGGTTGAGGGCTAGTACTGATGGCATGATGAAGAGCGTAGGGTTGAGCCA 110
                   AAGCTGGCAGAGGTCGAGGGCTACTACTGATGGCATGATGAAGAGCGTAGGGTTGAGCCA 108
                   AAGCTGGCAGAGGTCGAGGGCTACTACTGATGGCATGATGAAGAGCGTAGGGTTGAGCCA 100
                   AAGCTGGCAGAGGTCGAGGGCTACTACTGAAGGCATGTTTAAGAGCGTAGGGTTGAGCCA 85
                   ************** *** **** ****** ****** *  ******************

                   GACAATAATCGAGTAGATATTCAAAAAGCAGCAGACATGCCCAACTTTCTTCCTGTTGAG 180
                   GACAATAATACAGTACAAATTCAAAAAGGAGCAGACATGCCCGACTCTTTTC-TGTTGAG 169
                   AACAATAATACAGTAGAGATTCAAAAAGGAGCAGACATGCCTGACTTTTATC-TGTTGAG 167
                   AATAATAATAGTGTAGATATTTAAAAAGGAGCAGGCATGCCTGACTATTTTC-TGTTGAA 159
                   AAAAATAATATAGTAAAGATTCAAAAAGGAGTAGACATGCCAGACTCTTTTC-TGTTGAG 144
                    * ******   *** * *** ****** ** ** ******  *** *   ** ******

                   AGAACTTCGGT--G-G-GGGTTGACTTCGAGGTAAGTCGATG---CTTCATATGGCTATC 233
                   AGAACTTTGTT--G-G-AGGTTGACCTCGAGGCAAGTCGATG---CTTCATATGGCTATC 222
                   AGAAGTTTGAT--GTG-AGGTTGACCTCGAGGCAAGTCGATG---CTTCATATGGTTATC 221
                   AGAAGTTTGGTTAATGTAGGCTGAACTGGGAGGCAGTCGATGATGCTTCATATGCGAATC 219
                   AGAAGTTTGGTTGCTGTAGGCTGAACTGGGAGGCAGTCGATG---CTTCATATCCCTTTT 201
                   **** ** * *    *  ** ***  * *  *  ********   ********     *

                   TCTCGGGGATGGCCGGGGATGG-----------CTCATGCAGTCCAGTGGGAG-GTATT- 280
                   TCTCGGGGATGGTCGGGGATGGATAACCGTCCTTTCATGCAGTCCAGTGGGGG-GTATT- 280
                   TGTCGGGCATGTCCGGGGATGGATGGCCGTCCTTTCCTGCACTATGGCGGGGGCGTATT- 280
                   TCTCGGGGA-GCTCGGAGACTTATGGCCGCCGCTTCATGCACCATGGCGGGGGTATATTT 278
                   TCTCGGAGAGGTTCGGGGATGGATGGCCGTCCTTTCATGCACTATAGTGGCGGCGTACTT 261
                   * ****  * *  *** **               ** ****      * **  *   ** *

                   CAATTGGACTCCACATGCTGTTCCTATTGGCATCCACTATTGCCACTCCTGGTATAGGCA 340
                   TCTTTGGAGTCCACGTACAGTTCCTATTGGCATCCACTATTGCCGCTCCTGGTAAAGGCA 340
                   CCTTTGGACTCCACGTACAGTCCCTATTGGCACCCACTATTGGCACCCCTAGTATAGGCA 340
                   CCTTTGGACTCCACGTACAATCCCTATCGGTACCCTGTATTGCCACCCCTAGTTTAGGGA 338
                   CCTTTGGACTATACCTACAGTCCCTACTGGTATCCTGTGTTG-TACTCCCAGTATAGGCA 320
                      ***** *  ** * *  * ****  ** * **  * ***   * **  **  *** *
                                              #2             #1
                   CTATCCCGTTTTCC-GGATGAGTTTGGCCTCCGAAAAGTCGGCCTCCGAGCAGGG-CTTC 398
                   CTATCCCGATTTCC-GGGTTGGTTTGGCTTCCGAAAAGTCGGCCTCCGAGCAGGG-CTTC 398
                   CCATCCCGATTTCC-GGCTTCGTTTGGCCTCCGAAAAGTCGGCCTCCGAGCAGGG-TTTC 398
                   CTATCCCGATTACCAGGGGTAGTTTGGCCTCCGAAAAGTCGGCGTCCGAGCAGGGGCTCC 398
                   TTGTCCCGATTGCCCGGGGTGGTTTGGCCGCCGAAAAGTCGGCCTCCGAGCACGGGCTGC 380
                      ***** ** ** **     ******* *********************** **  * *

                   AGC-----GGTAGGCCTTGCAGCGGCGGTCGCTGC-TGTTAAGGTATGGACAAGCCCCTA 452
                   AGC-----GGTAGGCCTTGCAGCGGCGGTCGCTGC-TGTTAAGGTATCGACAAGCCCCTA 452
                   AGC-----GGTAGGCCATGCAGCGGCGGTCGCTGC-TGTTAAGGTATCGACAAGCCCCTA 452
                   AGA-----AATAGGCCCTGCAGCGGCGGTCGTTCCCTGTTAAGGTACGG-CAAGCCCCTG 452
                   AGCCCCTAGGCAAGCCCTGCAGCCGGGGTCTCTTGCAGTTAAGGTATCAACAAGCCCCTA 440
                   **          * *** ****** * ****  *     *********     *********

                   TTCACTACGGTTTCGACCCGTCGTTT------------GGCACTTTCAAACCGTCTTGCC 500 (-1 from initiation site)
                   TTCACTGCTGTTTCGACCATACTGTT------------GGCACTTTCTAACTCCCTCGCC 500
                   TTCACTACAGTTTCGACCATTCTGTT------------GGCTCTTTCCAACTCCCTCACC 500
                   TTCACTACTGTACAGACCATTCTGCT------------GACCCTTCCCAACTCATTCATC 500
                   TTCACTACTGTACCGATCATTCTGTTTACCCTTCCGTTGACTCTTTTGAACTCCCTCGCC 500
                   ****** * **    ** *   *   *             * * ***   ***     *    *
```

**F**

#### Alignment of *gliM* upstream regions (-500)

```
gene_00585.t1 (Al) -------------------------GGT-----CGGTTCGCA---TGAGTGAACGAGAGGC 28
NFIA_055370 (Nf)   ---------------------AGTGGT-----CGGTTCGCA---TGAGTGAACGTGAGGC 31
Afu6g09680 (Af)    AGAAATGGCCAAAGCTGCAGGAGTGGT-----CGGTTCGCA---TGGGTGAACGTGAGGC 52
gene_01179.t1 (Ap) --------------------------T-----CGGTTCGCA---TGAGTGAGCGAGATGC 26
gene_09467.t1 (Au) ----------------GGAGGCGTGATTCTTACTCTTCCCACTGTAAATAGA-GTAGTAC 43
                                           *     *   *** **    *    *    *     *

                   CGTCAAACGGGCATGGCAGCGCGTCGCCGG---ATTTGGACACGGCGAGAAGGAGTATG- 84
                   CGTCAAACGGGCATGGCAGCGCGTCGCCGG---TTTTGGACACGGCGAGAAGGAGTATG- 87
                   CGTCAAGCGGGCCTGGCAGCGCGTCGCGGG---GTTTGGCCATGGCGAGAAAGAGTATG- 108
                   CGTCAAACGGGCATGGCAGCGAGTCGCGGG---CTTTGGACACGGCGAGAAGGAGTATG- 82
                   AGTTTTATGAATACAGTTATAC-TTACCAGTGTATCCGTCCACTCGTAGGTTGAGTTTAC 102
                    **     *      *      *  *  *     *  *  **      **    **** *

                                     gliG
                   GAATGTTAGAGGCGTGATCCTTAGTCCTCCCACTATCAATAGAGTAGTGCAGTTTTA-TG 143
                   GAATGTTAGAGGCGTGATTCTTAGTCTTCCTACTGTCTATAA--TAATACAGCTTTA-TG 144
                   GCATGTTAGAGGCGTGATTTTTAGTCTTC-TACTGTATATAA-GTAATTCAGCTT----- 161
                   GAATGTTAGAGGCGTGATTCTTACTCGCCCCATTGTTAATAGAGTAGGACAGTTTTA-TT 141
                   GTTTATTCTAGGT-TGATTACTAGTGCCC--ACCA-GAATAAGCCAA-ACATTTCTACTA 157
                   *  * **   ***   ****   ** *    *   *      ***    *    **   *

                   CAATACAGTTATATCACTGAATCTATCCCTTTAATCGTATGTTGATTCTAGGTTGCTTAC 203
                   CAATACCGTTATACCACTGAATCTATTCCTTTAATCGTATGTTGATTCTAGCTTGCTTAC 204
                   CGA-------ATACCACTGAATTTATTCC--GAATCTTCTGTTGAT-CTGGGTTGCTCCC 211
                   CAATACAGTTATTCCAACCAATTTATCCGTTTAATCGTAGGTGGATTCTAGGCTGCTTAC 201
                   CGCTAGAAATAGTTGTCTAGACA---CCGTTTGGGCGTA-GTGGTTTGTTGGCTGACAAC 213
                   *          *          *       *        * *  ** * *   * *  **    *

                   TACTTAGCCCGATGTCTTGGCCCTGGGGTTGACCAGCCCTACGGTTCTACCTCGACTACA 263
                   TTCCTAGCCTGATGTCTTGGCCCTGGG-TTGACCAGCCCTACGGTCCTACGTCGACTGCT 263
                   TCCCTAGCCCGATGTCTTGGCCCTGGG-TCGACCAGCCCTACGGTTCTACCGCGACTACA 270
                   TAGCTA-CCCGACGGCTTGGCCCCGGGGTCGACCAGCCCTACGGCACTATCTCCACTACA 260
                   TAGTTA-CCCGATGTCTTGGCCCTGGG-TTGACCCGCCTCAGGGCCCTACTTCGACTACA 271
                   *   ** ** ** * ******** *** * **** ***  * **  ***   * *** *

                   GTATCCT--AGTATTT-CGCGCCTATGGAAGATTCTAGATTAAGTAGATCGGT-AATTTA 319
                   GTATCCT--TGTATTC-CGCGCCTATGGAAGATTCTAGGTTAAGTAGATCAAT-AATTTA 319
                   GTATGC---GGTATTT-CGCGCTTATG-AAGATTCTGGACGCATTCAGTCAGT-AATTTA 324
                   GTATCCTCCTGTATTTTCACGC-TATGGAAGATTCTGGATTATGTCGATCATTCAGTATC 319
                   GAATCCT---GTCTTTTCTCGCCTATGGAAGATTCTGGTTCATGTCAATCATTCAGTATC 328
                   * ** *     ** **  * *** **** ******** *       *    **  * * * *

                                                                            #1
                   CTTATTCGATCGATATAT-CAACTTGGTCACCGATTAAGTGCAATATTAGAGCTCGGCGA 378
                   CTTATTCGATCGATATATGCCAGTTGGTCACCGAATACGTGCAGTAATACAGCTCGGCGA 379
                   CTGATCCCATCGAGAGAT-TAAGTTGGTCACCGAATAGAACCATCAAGGTAGTTCGGAGA 383
                   TCTATTTCATGGATATAT-CCAGTTTGTCGCCGAATAGAATCATGATTATAGTTCGGCGA 378
                   TCAATTTGATCGATATAT-CCAGTTTGTCGCCGAGTAGAATCATGATTATAGTTCGGAGA 387
                       **    ** ** * **   * ** *** **** **     **  *     ** **** **

                   CCGAAAAGGCGATATGTCTCCATATGTCTCCGACAGTGTCGAGTATATGTATCCGAGGT- 437
                   CCGAAAAGGAGATATGTCTCCATATGTCTCCGACAGCGTCGAGTATATGTATCCGAGCG- 438
                   CCGAAAAGTGGCTATGTCTCCATATGTCTCCGACAGTGCTCAGTATATGTATCCGAGTG- 442
                   CCGAAATGCAGATATGTCTCCATATGTCTCCGAGAGTGTCGAGTATATGTATCCGAGCG- 437
                   CCGAAAAGGAA----------AAATGTCTCCGAGAGAGTCGAGTATATGTATCCGAGTGG 437
                   ****** *              * ********** ** *    ****************

                   CTCGCAATCAACAAGTATGCAATCCCAGACATCAACCACAGGTTTACAAACAAAGAGATA 497
                   CTGGCA-TCAACAAGTATGCAATCCCAGACATCAAGCACAGATTTACAAACAAAGAGATA 497
                   CTCACA-TCAACAGGTATGCAATCCTAGACCTCAAGCA-AAGCTAACGAGCA---AGAGA 497
                   CTCGCAATCAACAAGTGTGCAATCCCACAGACTAAGCACAGTTTTACGAAGAAAGGCACA 497
                   CTCGCCATCAACAAGTGTACAATCCCACAGACTAAGCACAGTTTCAAGAAGAAAGACACA 497
                   **   *  ****** ** * ****** * *    ** ** *   * *   *   *     * *

                   ACC 500
                   AAG 500
                   ACC 500
                   ACA 500
                   GCA 500 (-1 from initiation site)
```

**G**

### Alignment of *gliG* upstream regions (-500)

```
Afu6g09690 (Af)    ----------AGAAGCGGGTGAATCGCCCAGTCGCAGGGTGGTTTTCTTCTCGCGGGCAA 50
NFIA_055380 (Nf)   GAACGTCGCGACGAGCGGGTGAATCGCCGAGTCGTAGGGTGGTGTTCTTGTCTGGGGCGA 60
gene_00584.t1 (Al) -----------TGAGCGGGTGAATCGCCCAGTCGGAGAGTGGTTTTCTTGTCAGGTGCAA 49
gene_01180.t1 (Ap) ---CGTCACGAGGAGCCTGTTCGTCTCCAAGTCGTAGCGTCGTTTTCTTGTCTGGTGCAA 57
gene_09468.t1 (Au) ------------GAACGGGTGAGTCGCCCAGTCGTATTGTTGTTTTCTTGTCTGGTGCAA 48
                                  * *  **    ** ** ***** *  ** ** ***** **   * ** *

                   AGGGGGCGACGCTGGCAAAGGAAGGTTCCGCGTACGGGATGCCAAAGATGTCGAAGGTCA 110
                   AAGGGGCGACGCTGGCAAACGAAGGCTCAGCGTACGGGATGCCAAAGATGTCAAAAGTCA 120
                   AGGGGGCGACGCTGGCAAACGAAGGCTCAGCGTACGGGATGCCAAAGATGTCAAAGGTCA 109
                   AAGGGGCGACGCTCGCAAACGAAGGCTCAGCGTACGGGATACCGAAGATGTCGAAAGTCA 117
                   AAGGGGCGACGCTCGCAAACGAAGGCTCCGCGTACGGGATGCCGAAGATGTCGAAAGTCA 108
                   * *********** ***** ***** ** *********** ** ******** ** ****

                   GGTAGTGCGAGGGGACAATGACGGCCTTGATATCCAGTGCTTTGATGCCTCGATTCTCCA 170
                   GGTAGTGGGAGGGGACAATAACAGCCTTAATATCCAGTGCTTTGATGCCTCGATTTTCCA 180
                   AGTAGTGCGAGGGGACAATGACAGCCTTGATATCCAGGGCCTTGATGCCTCGATTTTCCA 169
                   GGTAGTGAGAGGGGACAACAACTGCCTTGATATCCAAGGCCTTAATGCCTCGATTTTCTA 177
                   GGTAGTGCGAGGGGACAACCACTGCCTTGATATCCAGCGCTTTAATGCCCCTATTCTCTA 168
                    ****** **********  ** ***** *******  ** ** ***** * *** ** *

                   GTACCGAGAGGCGTAAGTTGGAGCCGTAGGCAAAGTACCAGATTCCCCCATGAGGGTCTT 230
                   GTACAGATAGGCGTAAGTTGGAGCCGTAGGCAAAGTACCAGATTCCCCCGTGAGGGTCTT 240
                   GTACTGATAGTCGTAAGTTGGAGCCGTAGGCAAAGTACCAGATGCCCCCCTGAGGGTCCT 229
                   GCACAGATAGCCGTAGGTTTGAGCCGTAAGCAAAGTACCAGATTCCCCCGTAGGGGTCTT 237
                   GCACAGATAGGCGTAGGTTAGAGCCGTAGGCAAAGTACCAGATGCCCCCGTGAGGGTCTT 228
                   * ** ** ** **** *** ******** ************** ***** *  ***** *

                                    gliK
                   GCAACGCTGCTTTCCCCATGTTGCTTGCTCTTTGCGT-TCTGTAAGCAGGGGGACCAGGG 289
                   GCAACGTTGCTTTCTTCATGTTGTTTGCTCTTTGCGT-TCTGTAAACGGGGCGACCAGGG 299
                   GCAACATCGCTTTCCCCATGTTGTTTGCTCTTTGCGT-CCAGTAAACGGGATAACCAGCA 288
                   GCAACATCGCTTTCCCCATGTTGTTTGCTCTTTGCACGTCTGTAAGCAGCGCAACCAGGG 297
                   GCAACGTCGCTTTCCCCATGTTGTCTGCGCCTTGCACGTCAGTAA-CAGGGTGAACAGGG 287
                   *****   ******   *******  *** * ****     * **** * *     * ***

                                                             #1
                   AATTGTCTCTCTATCAATTTTAGCTACTGCTCAACGCACCGACGGCTTCCGAAAATTCGG 349
                   AATTATCTCTCTATTCACTTTAGCTACTTCCCAACGTACCGACGGCCTCCGAAAATTCGG 359
                   GATTATCTCTCTGTCCACTTTAGCTACTTCTCGAGGCACCGACGGCCTCCGAAAATTCGG 348
                   AAATATCTTTCTATCCACTGTATCTACGTCTCAACGCACCGACGGCGTCCGAAAATTCGG 357
                   AATTATCTCTCTACCCTCTTTAGCTACTTCTCAACGCACCGACGGCGTCCGAAAAATCGG 347
                    * * *** ***      * ** ****  * * * * ********* ******** ****

                   ATGGCGCGTTGGGGGGGGGGGGGGGGGGTGGCTACAAGAGGCG----GTTGGCCATAGA 405
                   ATGGCGCAT--------------GGGTGGGTGGCTACAAGAGGCT----GTTGGCTATAGA 402
                   ATGGCGCGT-------------GGGTGGGTGGCTACAAGAGGCT----GTTG--TATACA 389
                   ATGGCGTGT-------------GGGATGG---CTACAGGAGGCTGTTTGTTGCCTATAGA 401
                   ATGGCGTTT-------------GGGATGGTGGCCAGAGGAGGCTATTTGTTGCCTACACA 394
                   ******  *              ***  **   * * * *****     ****   * * *

                   TATAGCAGG-------GCCAAAAG-CCACAGATATTGTA--TGCCCCTACCCTTGCTAA- 454
                   TATAGGAGG-------GCCAGGGGACCACAGATATCGTT--TGCCCCTACCCTTGCTAAA 453
                   TATAGGAGA-------GC-GGGGGACCACAGATATCGTT--TGCGTCTACCCTTGCTAAC 439
                   TATAGGATA-------GCCAGAGGACCGCAGACATCGTTGCTGCCCCTACCATTGCTAAC 454
                   TATAGGGTAACCGGTAGCCAGAGGACCGCTGACATCCTTGCTGCCCCTACGCCTGCTAAA 454
                   *****            **      * ** * ** **  *   ***  ****   ******

                   ---------------CTTCACCAACCATTACCAGTTCTTGGACTGTTCCACACTCGTCAG 499
                   C--------------TTTCACACACTATTCTCAGTTCTCTGAACGTTCTACACTCGTCGA 499
                   CTACCCTCGCTGACCTTTCACAAACTATTCTCAGTTCTCTAATCATTCTATATCCGTGAA 499
                   T--------------AC-CACAACCTATTTTCAATTCTCTAATTGCCCTATACCCGTGAA 499
                   C--------------ATTCACTACCTATTT-CAATTACCTAATCATTCTAAACACGTGAA 499
                                     ***   * ***  ** **     *     * * *  ***

                   A 500
                   A 500
                   A 500
                   A 500
                   A 500 (-1 from initiation site)
                   *
```

#### H Alignment of *gliK* upstream regions (-500)

```
NFIA_055390 (Nf)   GTCCAAGATGGCGGGCACCATCTTGTGAGGATTGATCTCGCTGAACCAGGGGGCAGACAC 60
Afu6g09700 (Af)    ---------GGCGGGCACCATCTTGTGGGGATTGATCTCGCTGAACCAGGGGGCAGACAC 51
gene_00583.t1 (Al) ------------GGGCACCATCTTGTGAGGATTGATCTCGCTGAACCAGGGCGCGGACAC 48
gene_09469.t1 (Au) -----------------CCATTTTGTGAGGGTTGATCTCGCTGAACCAGGGAGCCGACAC 43
gene_01181.t1 (Ap) ----AGAATGGCGGGCACCATCTTGTGAGGATTGATCTCGCTGAACCAGGGCGCAGACAC 56
                                    **** ***** ** ******************** ** *****

                   CTTTTCGACGACGTAGATATCATGAGGAAGTCCGAGCTCTTCGAGTAGGATCAACGGTTT 120
                   CTTTTCGACGACATAGATATCGTGAGGGACTCCGAGCTCTTCGAGGAGAATCAACGGTTT 111
                   CTTTTCAACGACGTAGATGTCATGAGGAACTCCGAGCTCTTCGAGTAGGATCAACGGTTT 108
                   CTTTTCGACGACATAGATGTCATGAGGCACTCCGAGCTCCTCGAGCAAGATCAGTGGTTT 103
                   CTTTTCGACAACGTAGATGTCATGCGGCACTCCGAGTTCCTCGAGCAAGATCAACGGTTT 116
                   ****** ** ** ***** ** ** ** * ****** ** ***** *  ****  *****

                   GACGGTGTTGTGCGTGGACGTGGCGGTGCCTGTCGTC-ACT----TCATGAGTGGAGGG- 174
                   GACGGTGTTGTGTGTGGACGTGGCGGTGCCTGTAGTTGACT----GCATGAGAGGAGA-- 165
                   GACGGTGTTGTGCGTGGATGTGGCGGTGCCTGAAGCC-ACT----CCATGAGTGGAGAGT 163
                   GACGGTGTTGTGCGTGGAGGTGGCGGTACCTGTAATCAGGTTAACCTATGAGTGGAGGG- 162
                   GACGGTGTTGTGCGTGGATGTGGCGGTGCCTGCAGCCAAG---GCGTGAGGG-GGGGGG- 171
                   ************ ***** ******** ****                 * * ** *

                   TTTGCCGATGAGTTCGACTCGACTCACCTTTGACCACGAAAAGCACCAGCCTGTTTACTA 234
                   CTTGCTGATGCGT-CGACTCGACTCACCTTTGACCACGAAAAGCACCAACCTGTTGACTA 224
                   TTTGACGATGAGTTCAACTCGACTCACCTTTCACCACGAAAAGCACCAGCTTGTTTACTA 223
                   CTTGAAGGTCAGTTCCACCAGACTCACCTTTCACCACGAAAAGGACCAGTCGGTTTACCA 222
                   CTTGACGATCAATTTGACTCGACTCACCTTTCACCACGAAAAGCACCAGCCGGTTTACCA 231
                    ***  * *   *   **  *********** *********** ****    *** ** *

                                                gliK
                   CAAGATCGGAGGGTCGTTCGGACATTTCGACGAGTGTAGAACGTTCAGAGAACTGAGAAT 294
                   CAAGATCGGAGGGTCGCTCGGACATTCTGACGAGTGTGGAACAGTCCAAGAACTGGTAAT 284
                   CAAGATCGGAGGGTCGTTCGGTCATTTTCACGGATATAGAATGATTAGAGAACTGAGAAT 283
                   CAAGGTCTGATGGGCGTTCGGTCATTTTCACGTGTTTAGAATGATTAGGTAATTGAAA-T 281
                   CAAGATCTGACGGGCGTTCGGTCATTTTCACGGGTATAGGGCAATTAGAGAATTGAAAAT 291
                   **** ** ** ** ** **** *****   ***  * * *     *     ** **  * *

                   AGTGTGTGAAAGT--------------TTAGCAAGGGTAGGGGCA--AACGATATCTGTG 338
                   GGTTGGTGAAG----------------TTAGCAAGGGTAGGGGCA--TACAATATCTGTG 326
                   AGTTTGTGAAAGGTCAGCGAGGGTAGGTTAGCAAGGGTAGACGCA--AACGATATCTGTG 341
                   AGGTAGTGAATGT--------------TTAGCAGGCGTAGGGGCAGCAAGGATGTCAGCG 327
                   AGGTTGTGG-TAG--------------TTAGCAATGGTAGGGGCAGCAACGATGTCTGCG 336
                    *   ***                    ******  ****  ***   *  ** ** * *

                   GTCCCCTGGCCCTC-------CTATATCTATAGCCAACA----GCCTCTTGTAGCCACCC 387
                   G-CTTTTGGCCCTG-------CTATATCTATGGCCAACC----GCCTCTTGTAGCCACCC 374
                   GTCCCCCG-CTCTC-------CTATATGTATA--CAACA----GCCTCTTGTAGCCACCC 387
                   GTCCTCTGGCTACCGGTTACCCTATATGTGTAGGCAACAAATAGCCTCCTCTGGCCACCA 387
                   GTCCTCTGGCTATC-------CTATATCTATAGGCAACAAACAGCCTCCTGTAGCC---A 386
                   * *    * *           ****** * *   ****     ***** * * ***

                                                   #1
                   ACCC-------------ATGCGCCATCCGAATTTTCGGAGGCCGTCGGTACGTTGGGAAG 434
                   CCCCCCCCCCCCCCCCCAACGCGCCATCCGAATTTTCGGAAGCCGTCGGTGCGTTGAGCAG 434
                   ACCC-------------ACGCGCCATCCGAATTTTCGGAGGCCGTCGGTGCCTCGAGAAG 434
                   TCCC-------------AAACGCCATCCGATTTTTCGGACGCCGTCGGTGCGTTGAGAAG 434
                   TCCC-------------ACACGCCATCCGAATTTTCGGACGCCGTCGGTGCGTTGAGACG 433
                    ***             *  ********** ******** ********* * * * *   *

                   TAGCTAAAGTGAATAGAGAGATAATTCCCTGGTCGCCCCGTTTACAGA-ACGCAAAGAGC 493
                   TAGCTAAAATTGATAGAGAGACAATTCCCTGGTCCCCCTGCTTACAGA-ACGCAAAGAGC 493
                   TAGCTAAAGTGGACAGAGAGATAATCTGCTGGTTATCCCGTTTACTGG-ACGCAAAGAGC 493
                   TAGCTAAAGAGGGTAGAGAGATAATTCCCTGTTCACCCTG-TTACTGACGTGCAAGGCGC 493
                   TAGATACAGTGGATAGAAAGATATTTCCCTGGTTGCGCTGCTTACAGACGTGCAAAGAGC 493
                   *** ** *      *** *** * *   *** *    * * **** *    **** * **

                   AAACAAC 500
                   AAGCAAC 500
                   AAACAAC 500
                   AGACAAC 500
                   AAACAAC 500 (-1 from initiation site)
                   *  ****
```

### Alignment of *gliA* upstream regions (-500)

```
NFIA_055400 (Nf)    --GTCTATTCTATGCG-TGCATGGAAGCATGAAGAACGAGTGAGTACCGTGTAGAACTCA 57
gene_00582.t1 (Al)  TGGTTTATCCT--GCG-TGCATGGAGGCATGAAGAACGAGTGAGTACCTTGTAGCACTCA 57
gene_01182.t1 (Ap)  ------TTCGTATGCGGTGAATGGAAGCATGAAGAACAAGTGAGTACCTTGTAGAACTCA 54
gene_09470.t1 (Au)  ------------TGCGGTAAATGGAAGCATAAAGAACGAGTGAGTACCTTGGAGAACTCA 48
Afu6g09710 (Af)     ---------CTACGTG-CGTATGGAAGGACGAAGAACGAGTGAGTACCTTGCA-AACGCA 49
                                 * *    ***** * *  ****** ********** ** *  ** **

                    ACAGATGCATGTACACTTTGTCAGTGT--CCACATGTACTT-GTATTATAATGAAACAGG 114
                    GTAGATGCATCCACACTTCGTCAGTGC--CTACATGTACTTTGTATTATAATGAAAAAGG 115
                    ACCGATACGTCTACGCTTCATC-GTGT--CTGCATATGATC-A--GTATAAT-AAAAAGG 107
                    ATAGATACATCTACACTTCATC-GTGTTTCTGCATATAGTC-ATAGTACATT-AAAAAGG 105
                    CCGGACGCAT-------------GTGC--CTGAAGACAATG--------AAT---ACAGC 83
                       **  * *             ***    *    *      *          * *    * **

                    AGACAGTGT--CTAAATTAGATGCTG-ATTGGCTTACGGACCGTCCCTTGCCCTAA---- 167
                    ACAGAATATA-CAAAATAGGATGCTG-ATTGGCTGACGGGCCGTCCCTTGCCCTAG---- 169
                    AGGGAATCTAATAAA-CGCGCCATTGGATTGGCTGACGGGCCGTCCCTTGCCCTAG---- 162
                    AGAGAATCTAATAAAACGTGCCATTGGATTGGCTGACGGGCCGACCCATGCCCTAG---- 161
                    AGCCAGCGT--CTAAATTAGATGCTG-ATTGGCTTCCGGGCCGTCCCTTGCCCTAAAGCA 140
                    *   *   *    **    *    ** *******  *** *** *** *******

                    -----ATGAGGACGTGAGGTTGTTCGGAGACTTAACGGAGACTTTGCCGCCACGCCGAAT 222
                    -----ATGAGG--------TTGTTCGGAGACTTAACGGAGACTTTGCCGCCACACCGAAT 216
                    -----TTGAGG--------TTGTTCGGAGACTTAGCGGAGACTTTGCCGCCACGACGAAT 209
                    -----TTGAGG--------GTGTTCGGAGACTTAGCGGAGACTTTGCCGCCAAGCCGAAT 208
                    GAAACTTGAGG--------TTGTTCGGAGACTTAACGGAGACTTTGCCGCCACGCCGAAT 192
                          *****         ************** ****************   *****

                    CACAGCTGATGATG------CTGC---------AGACAGTCAG--ATGAGCAGATTTCTT 265
                    CACAGCGGATGATG------CTGC---------AAACAGTCAGAGATGAGCAGATTTTTT 261
                    CAGATGGGGTGATG------CTGC---------AAACAGGCCGGAATGAGCAGATTT-CT 253
                    CAGATGGGGTGACG------CTGC---------AAACAGTCCAAGATGAGCAGATTTTCT 253
                    CACAGCGGATGATGAAGATGCTGCGGCGGCGGTAGACAGTCAG--ATGAACCGGTTTCTT 250
                    ** *   * *** *      ****         * **** *    **** * * ***  *

                    TCAGT---------TCAGCAGGGATCGGCATTAGCCAGCCAGGTCGGAATTGTAAATAAA 316
                    TCAGTC--------TGAGCAGGGATCTGCAGTAGCCAGCCAGGCCGGAATTGTAAATAGG 313
                    TCAGTT--------CCAACAGAGATCTGCATTAGCGAGCCATGTCGGCTTTC-AAATGAG 304
                    TCAGAA--------TTAGCAGGGATCTGCATTAGCCAGCCATGTAGGCATTGGAAATGAG 305
                    CTAGTTCTTTTTGTTCAGCAGGGATCTGCAGGAGCCAGCCAGGTCGGAATTGGAAATGAA 310
                      **            * *** **** ***  *** ***** *  **  **  ****

                    A-ACTGCTGACAA-GT-AGCGAGTCTAGAAGGGCTTC-TTAGCTAAGAT-GCCAACACTA 371
                    A--CTCCTGACAA-GT-AGCGAGTCTAGAAGGGCTTC-TTAGCTAAGAT-GCAAACACGA 367
                    A-TCTGCTGACAA-GTGAGCGAGTCTAGAAGGGCTTC-TTAGCTAATATGGCCAACACTA 361
                    A-ACAGCTGACAATATAAGCGAGTCTAGAAGGGCTTCCTTAGCTAAGATCGCCAACACTA 364
                    AGACTGCTGACAA-GT-ACCGAGTCTAGAAGGGCGGC-TTAGCTAAGAT-GCCAACACTT 366
                    *  *  *******  * * ***************  * ******** ** ** *****

                    -GATGCATGAAAGTCTCCATGTCTCCGATGTCTGAATAGATATAGATGCGTCTCTTG--- 427
                    -GATGCATGAAAGTCTCCATGTCTCCGATATCTCTATAGATATAGACGCCCCTTTCGAC- 425
                    -GATGCATGAAAGTCTCCATGTCTCCGATGTC-CTATAGATATAGATGCGTCTCC-G--- 415
                    -GATGCATGAAAGTCTCCATGTCTCCGATGTCTCTGTAGATATAGATGCGTCGCT-GCTG 422
                    TGACGCATGAAAGTCTCCATGTCTCCGATGTCTGAGTAGATATAGATGCGTTTCGTGCGG 426
                     ** ************************ **    ********** **       *

                    -CTGTCTTCCACTGCT--CACTCATGTTTACAATC-ATCTTCAAGT---------CAATT 474
                    -TTGTCCTCCACTGCTCGCACTCATGTCTACAATC-ATCTTCAAGT---------CAATT 474
                    GCTATCGTCCACTGCTCGCACTGTTGTCTA--GTCCATCGTCAAGTCAATTGAATCAATT 473
                    GCTATCCTCCACTGCTTGTACTCATTTCTACAGTCCATCATTAAGT---------CAATT 473
                    TCTGTTTGTTGCTACT-GT-CTCCGGTCTACCATT-ATCGTCACGTCA--------AATC 475
                      * *      ** **    **    * **   *  *** * * **          ***

                    AAACATCCTTAC-TGACATCAATCATC 500
                    AACCATTTCTAC-CGACATCAAGCATC 500
                    GAACATCTCTACTTTACATCAATCATC 500
                    GAACATCTTTCTGTCACATCAATCATC 500
                    CAGC-TCTCTAC-TGACATCGACCATT 500 (-1 from initiation site)
                     * * *   *     ***** * ***
```

**J**

### Alignment of *gliN* upstream regions (-500)

```
                     gliF
gene_09471.t1 (Au) ---------------AAGCCGAGGTTGAATTCCGGCCGAGATTCTAGATTTA-------- 37
gene_01183.t1 (Ap) ---------ATGCTGAGGCGGAGGTCGAATTCCGGCCGAGATTCTAGG---A-------- 40
gene_00581.t1 (Al) -----------TCGTAAACGGAGG--AAGTCCC-----AGATATCAAGC--A-------- 32
NFIA_055410 (Nf)   ------------CTGAGGCGGAGGTCGAGTTTAGGCCGAGAGTATAGGCGTATACAGCCA 48
Afu6g09720 (Af)    TCTTCCCGGATGCGGGGGCGGAGGTGGAGTTGAAGCCACGAGTATAGGTGTATACTGCGC 60
                                     * ****   * *         **    *     *

                   GCGAGGTGTTTATTCTGTCGTAGTG-TTGGGATTACTTGATTCGCTTTGAAAA-T-GAGG 94
                   G-GCGGTTGGTAGTATCTTGTAGCGGTTTAGATTACTTGATTTGCTTTGAAAA-TTGAGT 98
                   A-GTGGTTTATTCCATGCCTTTGGGGTTGAAGTGACTTGGCTGGCTTTGAAAAATTGAGA 91
                   GAGGGTGCTTTTCTACGTCGTAGCATTG-AGGTTACTTGGCTTGCTTTGAATT--TGAGG 105
                   GAGAGTGCTTTCTGACATCGTAGCGTTGGAGGCTCCCTGGCTTCC--------------- 105
                     * *     *         * *   *        * **  *  *

                   TTAGAGCGCTACTTTACACTGAAAT----------GACATAAAATTAGAACAAAATGTGT 144
                   TTACAGCGTTGCTTTACCGTGGAATAGTGTCGAACGATATGAAATTAGATCAATGTGTGT 158
                   TTTCAGCGCTCCTTTACTGTGGGATAGTATCAAACAATATGAAATTAGAACAA-ATGTAT 150
                   TTACAGCGGTACTTA--CGTCAAATTGTATTGAAGAATCTGAAATTAGAACAAAATGTGT 163
                   ------------------GTGAAATTGTATTGACCGATCTAAAATTAGAACCAAATGTGT 147
                                      *   **           *  * ******** * *  *** *

                   CCG---------GAGTATACT-TTGAAGCAAG---------CTTTGATACCAGATGTGTC 185
                   CCG---------GAGTATACT-TTGAAGCAAG---------TTTTGATATCAAATGTGTC 199
                   CCGAATGTATCCGAGCATATCATTAACACGAGGTCTATCGCCTTTGAT--CAT-TATGTC 207
                   CCG---------GAGTATAAA-TTGAAGCAAGTTCCTCGTTTTG-GATGCCAA-TATGTC 211
                   CCG---------GACTATAAA-TTGAAGCAAGTTCCTTGGTTTGTGATGCCAA-TATGTC 196
                   ***         **  ***   ** *  * **          *  ***  **  * ****

                   ATGA-TTAATTTCCATTGTAAGCCTGGCCCTAGTCTACAGCTTCATCGGCCGACATGAGA 244
                   ATGA-TCCATTTCTATTGTAAGCCTGGTCCGAGTCTACAGCTTCATCCGCCGAGATGAGA 258
                   ACGAATTCATTTCTATTGTAGGCTTCGTCCGAGTCTACAGCTTCATCCGCCGAGATGAGA 267
                   ATGAAT-CATT-CTATTGTAAGATTGGTCCCAGTGTACAGCTTCATCCGCCGAGATCAGA 269
                   CTGGTTTCATG-CTACTGTAGGATTGGCCCCAGTGTACAGCTTCATCCGCCGAGATGAGA 255
                     *  *  **  * * **** *  * * ** *** ************ ***** ** ***

                   CCCCGAGTGCTACTACGCCGATCGTAAGGTCCAATGCATTCAC-AAATGGCATGAGCAAA 303
                   CCCCGAGTGCTGCTACGCCGATCGTAAGATCGAATGCATTCAA-AAATAACCTGCAGAAA 317
                   CTCCGAGTGCTACTACTTCAATCGTAAAATCGAAAGCATTCAACAAATGTCATGGCCAAG 327
                   CTCCGAGTGCTACGAAACTA-TCGTACGATTGAATGGATTCAT-AAACAAATTGAACAAA 327
                   CTCCGAGTGCCACGAAACCAATCGTAGGATTGAATGGATTCAT-GAACAAAGTGAACAAT 314
                   * ********  * *      *****   *  ** * *****   **     **   **
                                                        #1
                   GCCGAGACCCCTGCGGTCACCAAAATTTCTCGATTTCGGTCTCCGAAACGGGTGGAGTGT 363
                   GCCGTAGCTCCTGTGGTCATTGGGATT-CTC-ATTTCGGTCTCCGAAACTGGTGGAGCAT 375
                   GCCGTGACCCATACGGTCATTTAGATT---CAATTTCGGTGTCCGAAACGGGTGGAGTGT 384
                   GCCATGACGGATACGGTCATTGAGATT---CGATTTCGGAGACCGAAACGGGTGGAGTGT 384
                   -CCATGGCAGATACGGTCATTCAGATT---CGATTTCGGAGACCGAAACGAGTGGAGTGT 370
                    **    *   *  *****     ***   * *******   *******  ******  *

                   TCGACATGTCTGATGAACGCATCCATGGTCTATATATACCTATGTAGTCCTTGTACGCCG 423
                   TCGACATGTCCGATGGATGCATCCATGGTCTATATATA----TGTAGTTGTTGTATTCCA 431
                   TCGACAT-------GAACG------TGGTCTATATA-AGTGGTGTA-TCTATCTATACCA 429
                   TCCATATGCCTGATGAAGGTATC--GTGTCTATATATA--TACGTGGTCTATCTATACCG 440
                   TCAATATGCCTGA-GAGGGGATCC-GGGTGGATATATA--TACACGGTCCCTGTATATCG 426
                   ** * **       *   *        **  ***** *         *   * **   *

                   TGTTGAA-AGGCAA---CATTCATCTAGGACCTGTCCAACACTCACCATTGTCCTTGACC 479
                   GGTCGAA--GTCAA---CATTCATCTAAGACGTGCCCAA-ACTCATTACTGTCT---ACC 482
                   GGTTCAA-AGTCAA---CAGTCATCTACGACGTGCCCAA-ACTCATCACTGTCT---ACC 481
                   -GTTGAGCAG------CATCTAACCTGTA--------AACAACCATTACTGTGT---ACC 482
                   AGTTGAGCAGACGAATCCACTAACCCATCGC-TGTCCAGCACCCATATCCGTGT---GCC 482
                    **  *   *          * * *            *  *  **     **      **

                   AGATAACTGCAAACCGCCACC 500
                   AG---ACTGCAAACCGCCACC 500
                   AGA-AACTGCGA-CTGCCACC 500
                   AGA-AATTGCGA-CTGC-ATC 500
                   AGA-AACTGCGA-CTGC-ATC 500 (-1 from initiation site)
                   **   * *** * * ** * *
```

**K**

#### Alignment of *gliF* upstream regions (-500)

```
Afu6g09730 (Af)     -----CGGGTTGCGGTGGTCCCATCCGAGGACGTTGTGCATGTGCTGGGTCTTTGCATTG 55
NFIA_055420 (Nf)    ---GCGGGGTTGCGGTGGTCCCATCCGAGGACATTGTGCATGTGTTGGGTCTTCGCATTG 57
gene_00580.t1 (Al)  TCGGCGGGGTTGCGGTGGTCCCATCCGAGGACATTGTGCATGTGTTGGGTCTTTGCATTG 60
gene_09472.t1 (Au)  -----------------------TCCGAGGACGTTGTGCATGTGTTGGGTCTTTGCATTG 37
gene_01184.t1 (Ap)  ----------------------ATCCGAGGACGTTGTGCATGTGTTGGGTCTTTGCATTG 38
                                           ********* *********** ******** ******

                    CGGTAGACGCCAGAGTCGAAGACGACTGCCGTGTGCAGTTGACGGGCCAGGCCGGTGGCT 115
                    CGATACACGCCAGAGTCGAAGACGACAGCCGTGTGTAGTTGACGAGCGAGGCCAGTGGCT 117
                    CGGTACACGCCAGAGTCGAAGACGACTGCTGTGTGTAGTTGACGAGCCAGGCCGGTGGCT 120
                    CGATACACGCCAGAGTCGAAGACGACCGCCGTGTGTAGCTGGCGAGCCAGGCCAGTGGCC 97
                    CGGTACACGCCAGAGTCGAAGACGACTGCCGTGTGCAGCTGGCGAGCCAGGCCGGTTGCT 98
                    ** ** ******************** ** ***** ** ** ** ** ***** ** **

                    GTCGAGAGACCAGCGGGACCGGCGCCGATGATGAGCACGTCGACGAGCAGGGCTCCGTTG 175
                    GCCGAGAGACCAGCCGGACCGGCGCCGATGATGAGCACGTCGACGAGGAGGGCTCCGTTG 177
                    GTTGACAGACCAGCCGGACCGGCGCCGATGATGAGCACGTCGACGAGGAGGGCTCCGTTG 180
                    GTTGAGAGACCTGCCGGACCTGCGCCGATGATGAGCACGTCGACGAGGAGCGCTCCGTTG 157
                    GTCGAAAGACCAGACGGACCTGCGCCGATGATGAGCACGTCGACGAGGAGCGCTCCGTTG 158
                    *  ** ***** *  ***** ************************* ** *********

                                         gliT
                    GAGAGTAGTTTGCCGATCGACATGGTTGTGGTAT-GCGCGAGAGTAGTGGGATGCCAGAA 234
                    GATAGTAGTTTGCCGATCGACATGTTTGTGGTAT-GAGAGCAAGTATTTAAATGCAAGAA 236
                    GATAGTAGTTTGCCGATCGACATGTTTGTGGTAT-TCAAGATGGTAAGTTTATGCAAGAG 239
                    GATAGTAGTTTGCCGATCGACATGTCTGTGGTATTGTGAGATTGTGATTGTATGGAAGAG 217
                    GCTAGTAGTTTGCCGATCGACATGTCTGTGGTATTGTGAGATAGTAATTGTATAGGAGAG 218
                    *  *********************  ********     *   **       **   ***

                    --ATTACCAAAAGAAAAGAA----AAACCAGAATGAAAGAGCCT--------CGTCGGGG 280
                    --ATGACGAAGAGAACAGAA----AAAA-AGAAAGTAAGAGCCT--------CGTTTGGG 281
                    --ATTAACAAGAGAAAAGAA----ATGA-AGAAAGAAAGAGCCT--------CGTTTGCG 284
                    TTATTATCCAGAGAAAAGGA--TTAGAAGAAAATGTAAAAATTTTGCATCGACGGTGAGG 275
                    --ATTACGCAGAGAAAAAAAAATTAAAAGAAAATCTAAAATTTTTGGATCAACGATGAGG 276
                      ** *   * **** *  *    *    * **   ** *   *         **    *

                                                                     #1
                    TGGTAT------TTATATAC---GGCGCATGGGTTCGGACACTTTCTTCTCGGCATCCGA 331
                    TCGTAT------TTATATAC---CTCAGATGGGTTCGGACACTTTTTTCTCGGCATCCGA 332
                    TCGTAT------TTAAATAC---CTCAGATGAGTTCGGACACTT--TTCTCGGGATCCGA 333
                    TTGCCTGTCTTTTTATATACACCTCCAGA---GCTCGGGGACTT--TCCTCGGCAGCCGA 330
                    TTGCGTGTCAGTCTATATAC---CTGAGATTGGCTCGGAAACTT--TTGTCGGCAGCCGA 331
                    * *  *        ** ****         *   * ****  ****   *  **** * ****

                    AAAATGTTTCCGCGGAATGGGCCAATGGTCATGCAATTGATCTATAAGACACATGTTGC 391
                    AAAA-TGTTTCCGCGGGATGGGCCAATGGCCATGCAATTGATGTATAAGACGCATGTTGC 391
                    AAAA-TGTTTCCGCGGGATGGGCCAATCGCCATGCAAGCGATGTATAAGACGCATGTTGC 392
                    AAAA-TGTTTCCGCGGGATCGGCCAATCGCCATGCAAGTGATGTATAAGACACAGGTTGC 389
                    AAAA-TGTTTCCGCGGGGTGGGCCAATCGCCAGGCAAGTGATGTATAAGACGCAGGTTGC 390
                    **** ***********  * ******* * ** ****  *** ******** ** *****

                    ACCACATTCGGATGGACCGAGTAGAATTAAATAGAGTCGATGGGGAATTGATTGCAGAAT 451
                    GCCACATTCAAAAGAACCAAGTAGAATTACATATAGCCCATGGAGAGTCGATTACAGAAT 451
                    GCCATGTTTGAAAGAATCAAATAGAATTAAATATAGTTCATGGAGAACTGCCCATAGAAT 452
                    GCCACGTTCAAAAGAAACAAATAGAATTACATATAGTTCACGGAGAGCTGACCCGATAAT 449
                    GCCATATTCGAATGAATCAAATAGAATTACATATAGTTCATGG-GAGCTCACCTGATAAT 449
                     ***  **    * * * * * ******** *** **   * ** **         * ***

                    TGAATTGAATTCC-C-CAAAGCAAAGAGAAA-GAGAAGTGAAAGCACGCGCC 500
                    TGAATTAAACCTCGC-TGAAGCAAAGAGAGA-TAGATCTGAC-GCACTCGCC 500
                    TGAATTGAGCCTCTCTCAAAGCAAAGAGTGA---GATCTGAA-GCACTCCCC 500
                    TGAATTCAACCTCGCTTGAACCAAAGAGCGACGCGATCGGAA-GCACACGCC 500
                    CGAATTGAACCTCGCTTGAACCAAAGAGAGACGAGATTCGAA-GCACACGCC 500 (-1 from initiation site)
                     ***** *    * *   ** *******  *   **   **  **** * **
```

**L**

### Alignment of *gliT* upstream regions (-500)

```
Afu6g09740 (Af)    -----TGCGAGTACCTTGGCATGGCCCTCCTCCAGCGTTTTCACAATATCTTGATCACCA 55
NFIA_055430 (Nf)   ---GTTGCGAGTACCTTGGCATGGCCGTCCTCCAGCGTTTTCACAATGTCTTGATCACCA 57
gene_00579.t1 (Al) GTTGTTGGGAGTACCTTGGCATGGCCTTCCTCCAGCGTTTTCACAATATCCTGATCACCA 60
gene_01185.t1 (Ap) -----------------------GCCCTCCTCCAGGGTTTTCACAATGTCTTGATCACCA 37
gene_09473.t1 (Au) ----------------------GACCCTCCTCCAGTGTTTTCACAATGTCTTGATCACCA 38
                                           ** ******** *********** ** *********

                   ACACTCTGCAGGACCGGCAAGTCCAGCGCCTTTAGACTCTTTGTCTTTTGCAAAAGCAGT 115
                   ACACTTTGCAGGACCGGCAAGTCCAACGCCTTTAGACTCTTTGTCTTTTGCAAAAGCAGC 117
                   ACACTTTGGAGAACCGGCAAGTCGAGTGCTTTCAGACTTTTTGTCTTTAGCAAGAAGAGT 120
                   ACACTTTGCAGAACCGGCAAGTCCAGCGCCTTCAGACTCTTTGACTTTTGCAAAATCAGT 97
                   ACACTTTGCAGAACCGGCAAATCCAGCGCCTTCAGACTCTTTGTCTTTTGCAAAATTAGT 98
                   ***** ** ** ******** ** *  ** ** ***** **** **** **** *  **

                   CGATATACCCAGTACAGCAGGCCGACTCCAAACGAGACCGCGACGGCGGTCCGCACGGAC 175
                   CGATATACCCAGTAGAGCAGGCCGACTCCAAACGAGACCGCGACTGCGGTTCGCAGGGAC 177
                   CGATATACCCAGTATAGCAGACCGACCCCAAACGAGACCGCGACTGCAGTTCGCAGGGAC 180
                   CGATATACCCAGTATAGCAGACCGACCCCAAACGAGACTGCGACTGCAGTTCGCAAAGAC 157
                   CGGTATACCCAGTATAGCAGGCCAACGCCAAAAGAGACTGCGACTGCAGTTCGCAGGGGC 158
                   ** *********** ***** ** ** ***** ***** ***** ** ** ****  * *

                   gliF
                   TGGGGGAGGTAGACTTGGTCCATGGCGCGTGCTTTCACTTCTC-TTTCTCTTTGCTTTG- 233
                   TGGGGGAGGTAGACTTGGTCCATGGCGAGTGCGT-CAGATCTA-TCTCTCTTTGCTTCA- 234
                   TGGGGGAGGTAGACTTGGTCCATGGGGAGTGCTT-CAGATCT---CACTCTTTGCTTTGA 236
                   TGGTGGAGGTAGACTGTGTCCATGGCGTGTGCTT-CCGATCGCGTCGCTCTTTGGTTCAA 216
                   TGGTGGAGGTAGACTGTGTCCATGGCGTGTGCTT-CGAATCTCGTCTCTCTTTGGTTCAA 217
                   *** ***********  ********* * **** * *   **      ******* **

                   GGGAA-TTCAATTCAATTCTGCAATCAATTCCCCATCGACTCTATTTAATTCTACTCGGT 292
                   GCGAGGTTTAATTCAATTCTGTAATCGACTCTCCATGGGCTATATGTAATTCTACTTGGT 294
                   GAGAGGCTCAATTCAATTCTATGGGCAGTTCTCCATGAACTATATTTAATTCTATTTGAT 296
                   GCGAGGTTGAATTCAATTATCGGGTCAGCTCTCCGTGAACTATATGTAATTCTATTTGTT 276
                   GCGAGGTTCAATTCGATTATCAGGTGAGCTC-CCATGAACTATATGTAATTCTATTTGAT 276
                   * **   * ***** *** *          ** ** *   ** *** ******** * * *

                   CCATCCGAATGTGGTGCAACATGTGTCTTATAGATCAATTGCATGACCATTGGCCCATTC 352
                   TCTTTTGAATGTGGCGCAACATGCGTCTTATACATCAATTGCATGGCCATTGGCCCATCC 354
                   TCTTTCAAACATGGCGCAACATGCGTCTTATACATCGCTTGCATGGCGATTGGCCCATCC 356
                   TCTTTTGAACGTGGCGCAACCTGTGTCTTATACATCACTTGCATGGCGATTGGCCGATCC 336
                   TCATTCGAATATGGCGCAACCTGCGTCTTATACATCACTTGCCTGGCGATTGGCCCACCC 336
                    * *   **   *** ***** ** ******** ***   **** ** * ******* *   *

                                   #1
                   CGCGGAAACATTTTTTCGGATGCCGAGAAGAAAGTGTCCGAACCCATGCGCCG---TATA 409
                   CGCGGAAACATTTT-TCGGATGCCGAGAAAAAAGTGTCCGAACCCATCTGAGG---TATA 410
                   CGCGGAAACATTTT-TCGGATCCCGAGAA--AAGTGTCCGAACTCATCTGAGG---TATT 410
                   CGCGGAAACATTTT-TCGGCTGCCGAGGA--AAGTCCCCGAGC---TCTGGAGGTGTATA 390
                   CGCGGAAACATTTT-TCGGCTGCCGACAA--AAGTTTCCGAGCCAATCTCAGG---TATA 390
                   ************** **** * ****  *  ****  **** *   *     *    ***

                   TAAAT------ACCACCCCGACG--------AGGCTCTTTCATTCTGGTTT--TTCTTTT 453
                   TAAAT------ACGACCCAAACG--------AGGCTCTTACTTTCTT-TTT--TTCTGTT 453
                   TAAAT------ACGACGCAAACG--------AGGCTCTTTCTTTCTT-CAT--TTCTTTT 453
                   TAAAAAGACAGGCAACCTCACCGTCGATGCAAAATTTTTACATTTTCTTCTAATCCTTTT 450
                   TAGACTGACACGCAACCTCATCGTTGATCCAAAAATTTTAGATTTTCTTTTAATTTTTTT 450
                   ** *         * **      **         *    * **    ** *    *   *   * **

                   --CTTTTGGTAAT--TTCTGGCATCCCACTACTCTCGCGCA-TACCACAACC 500
                   --CTCTTCGTCAT--TTCTTGCATTTAAATACTTGCTCTCA-TACCACAAAC 500
                   --CTCTTGTTAAT--CTCTTGCATAAACTTACCATCTTGAA-TACCACAAAC 500
                   --CTCTGGATAATAACTCTTCCATACAATCACAATCTCACAATACCACAGAC 500
                   TTCTCTCTGCGTAAT--CTCTCCTATACAATTACTATCTCACAATACCACAGAC 500 (-1 from initiation site)
                     ** *   * **    ***    **       **    *    * *******   *
```

**M**

### Alignment of *gliH* upstream regions (-500)

```
NFIA_055440 (Nf)   TCCCTTCCATGAGACCAGTGTGCCTGGAGTATTTGCCGTGGGGACTGTGCCACGCCGTT 60
Afu6g09745 (Af)    GCCCTTCTATGAGACCAGTGTGCCCGGAGTATTTGCCGTGGGGATTGTGCCACGCCGTT 60
gene_00578.t1 (Al) --------ATGAGACCAGTGTACCTGGAGTATTTGCAGTCGGCGATTGTGCCACGCCGTT 52
gene_09474.t1 (Au) -------------ACCAGTGTGCCCGGAGTATTCGCAGTGGGGATTGTGCCACACCGTT 47
gene_01186.t1 (Ap) -----------AGACGAATGTGCCCGGAGTATTTGCCGTAGGGGATTGTGCCACACCATT 49
                                ** * *** ** ******** ** ** ** ** ******** ** **

                   AAAGGCTGTCACCCCGGCGGTGGCGATGGGATCTTTGGCTGCTGGAGGTCTAGTGGCTCA 120
                   AAAGGCCGTCACGCCCGCGGTGTCGATGGGATCTTTGGCCGCTGGCGGTCTCGTGGCTCA 120
                   AAAGGCCGTCACCCCGGCGGTGGCGATGGGATCTTTGGCTGCTGGAGGTCTGGTTGCTCA 112
                   GAAAGCTGTCACCCCCGCGGTATCAATGGGATCTTTGGCTGCTGGTGGTCTGGTTGCTCA 107
                   GAAGGCTGTCACTCCTGCGGTGGCGATGGGATCTTTGGCTGCGGGAGGTCTGGTTGCCCA 109
                    ** ** ***** ** *****  * ************** ** ** ***** ** ** **

                   GTTGCAGGCTCAGTCATTGCCGGAGTTTCGTCTCGATCAGGAGCTATAGGAGTTTGTTAT 180
                   GCTGCAGGCTCAGGCATTGCCGGAGTTTCGTCTCGATCAGGAGCTATAGAAGTTTGTTAT 180
                   GTTGCAGGCTCAGCCATTGCCGGAGTTTCGTCTCGATAAGGAACTATAGGAGTTTGTTGT 172
                   GCTGCAGGCTCAGCCATTGCCGGAGTTTCGTCTCGATCAGGAACTGTAGGAGCATGTTAT 167
                   GCTGCAGGCTCAGCCATTGGCGGAGTTTCGTCTCGATCGGGAACTGTAGGAGCATGTTAT 169
                   * *********** ***** ****************  *** ** *** **  **** *

                   gliT
                   AG--TTGTACATAGTT---TTTAGTT---TC-CGAGGTTCAATTGAA--ATCAAGTGACT 229
                   AG--CTGTACATAAATAGCTGTACATACGTC-TTAGATTCGCTAGGTTCATGAATTGACT 237
                   AG--TTGTACATAGTTT-TTCTAGTT---TCGCGAGGTTCAATTGAA--ATCAAGTGACT 224
                   AGAGTTGTACATAGTT---TCTAGCT---TTGTGAGGTTCAATCAA---ACCAAGTGACT 218
                   A-ATTTGTACATAGTT---TCTAGCT---TTGTGAGGTTCAATTAA---ACCAAGTGACT 219
                   *     ********  *   * **  *   *    ** ***  *      *  ** *****

                   TTCCTTATTTCTGTCATTA-GTTCTAAATCGTGAGTTATCAATCTCCAA-AGATCCGTAA 287
                   TTCCTTA---CTGCCAGGA-TTCGAGAATTGTGAGGCACCA-TCTCCAA-AGATCTGTGA 291
                   TTCCTTATC-CTGCGAATAGTTCCAGATTCGTGAGCTATCTGTCTGCAAGAGATTCGTAC 283
                   TTCCTATTTCCTGC-AGTAGCTTGAGCCTAGTGAGCTGTCCTTCTGAAAGAGATTCGTAA 277
                   TTCCTGTTTCCTGC-AGTAGCTTGAGCATAGTGACCTATCCGTCTGAAAGAGATTCCTAA 278
                   *****      ***  *  *  *       * ****     *  ***  ** ****   *

                   CAAA-TAAATAAGAATGGAAGACGCAAAGAAACT-TTCTC-CGG----TTGGG-CAGGTC 339
                   CAAA-TAA----GAGTA-AAGACGCAAGTAACCT-TTCTC-CAG----TTGG--CAGGTG 337
                   CGAAGTAGATGAGAATC-AAGACGCGGAGATGCA-GGCGCACAAGAC-CTGGCGCCGGTC 340
                   CAAACTTGATGAGAATGAAAGATGCATCGAAAAAAGTCGCTCAGGTCCACGGTGGTGACC 337
                   CAAATTAGACGAGAATGAAAGACGCAAAGAAAGA-GTCGCTCAGGTCAACGGTAGTGATA 337
                   * ** *       ** *   **** **     *       * * *         **    *

                   TGAACCTCAACGTTGAAAT-CGCTCGCTGTGGATAGCTTCATATCCCGCCGGTGGTACA- 397
                   TGAACCTTGACGTTGAAAT-GGCCCCCTGTGGATAGCTTCATATCCCGCCGGTGGGATA- 395
                   TGAGACTTCATATTGCATTTGGCTCGCCGTGGATAGCTATATATCCCACCGGTGATACA- 399
                   TGAGACTCGATGCTAAAAT-AGCTCGCCATGGAAAGCTTCATATCACGCCGACGGTACAG 396
                   TGAGACTCGATGCTAAAAT-AGCTCGCCGTGGATAGCTTTATATCACGCCGACGGTACAG 396
                   ***  **  *    *  * *  ** * *  **** ****  ***** * ***  *  * *

                         #1
                   AACTCGGCGACCGAAAACTACGGCATTCCTCGGAGAACGCCTCC-TATATCATAATATAA 456
                   AAATCGGCGACCGAAAACTGCGGCATTCCATGGATAGGGTCGCCGTTTATTATAATATAA 455
                   AAATCGGCGACCGAAA-CTGCGGCATGACTCTGATGGAGACACA-TATATCATAATATAA 457
                   AAATCGGCGACCGAAA-CTGCGGCGTTACTCAGACAGACATCCCATATATGATAATATAA 455
                   AAATCGGCGACCGAAA-CTGCGGCATTACTCAGACAGAGACCCCATATATGACAATATAA 455
                   ** ************** ** **** *  *   **        *  * *** * *******

                   CACCCCCA-GACCTCAGACAGATTCTCAACAACAGCAAC-ATCACC 500
                   CACCCCCAAGACGGCAGACAGATCTTCAACGACAGCAAC-AGCACC 500
                   CACCCCCAGAACG-CAGACA-ACCCTCAACAACAGCAAC-ATCACC 500
                   TGCCCCCAGAATG-CGGACAAATCCTCAATGACAGCAACTATCACC 500
                   TACCCCCAGAATG-CGGACAACTCCTCAATGATAGCAACTATCACA 500 (-1 from initiation site)
                     ******  *   * ****     ****  * ****** * ***
```

Fig. S3
